# Gene-set Enrichment with Regularized Regression

**DOI:** 10.1101/659920

**Authors:** Tao Fang, Iakov Davydov, Daniel Marbach, Jitao David Zhang

## Abstract

**Motivation:** Canonical methods for gene-set enrichment analysis assume independence between gene-sets. In practice, heterogeneous gene-sets from diverse sources are frequently combined and used, resulting in gene-sets with overlapping genes. They compromise statistical modelling and complicate interpretation of results.

**Results:** We rephrase gene-set enrichment as a regression problem. Given some genes of interest (*e.g.* a list of hits from an experiment) and gene-sets (*e.g.* functional annotations or pathways), we aim to identify a sparse list of gene-sets for the genes of interest. In a regression framework, this amounts to identifying a minimum set of gene-sets that optimally predicts whether any gene belongs to the given genes of interest. To accommodate redundancy between gene-sets, we propose regularized regression techniques such as the *elastic net.* We report that regression-based results are consistent with established gene-set enrichment methods but more parsimonious and interpretable.

**Availability:** We implement the model in *gerr* (gene-set enrichment with regularized regression), an R package freely available at https://github.com/TaoDFang/gerr and submitted to *Bioconductor.* Code and data required to reproduce the results of this study are available at https://github.com/TaoDFang/GeneModuleAnnotationPaper.

**Contact:** Jitao David Zhang, Roche Pharma Research and Early Development, Roche Innovation Center Basel, F. Hoffmann-La Roche Ltd. Grenzacherstrasse 124, 4070 Basel, Switzerland.

## Introduction

Gene-set enrichment analysis sheds light on biological functions of gene lists derived experimentally or computationally. Popular choices include Fisher’s exact test, GSEA (Subramanian *et al.*, 2005), and CAMERA (Wu and Smyth, 2012), but many other tools using various statistical models and procedures are available (Rahmatallah *et al.*, 2014; de Leeuw *et al.*, 2016; Rahmatallah *et al.*, 2016; Geistlinger *et al.*, 2019). Methodologically, they can be classified into self-containing and competitive methods (Goeman and Bühlmann, 2007). Practically, they are applied in contexts where the user wishes to gain biological insight from a set of Genes of Interest (GoI hereafter) among all possible genes of consideration (*background* hereafter).

Most available methods are based on two implicit assumptions. First, only one set of GoI is tested for enrichment of gene-sets a time. If there are multiple sets, each set is tested independently and the results are combined. Second, enrichment is tested one gene-set a time, assuming independence between gene-sets. There are good reasons for both assumptions. Treating both sets of GoI and gene-sets independently simplifies software implementation and allows analysis even for one set of GoI with one single gene-set. Computational techniques such as parallelization (Sergushichev, 2016) and approximation (Zhang *et al.*, 2017) can speed up execution under these assumptions. Finally, when the gene-sets are derived from a single data source, the redundancy between the gene-sets, *i.e.* the proportion of shared genes between two gene-sets, can be small. In such cases, the gene-sets may indeed be treated as independent from each other.

Both assumptions, particularly the independence between gene-sets, are however often violated in practice. Gene-sets from different sources are commonly aggregated for enrichment analysis, and consequently redundant gene-sets are often called as hits. For instance, DAVID (Huang *et al.*, 2009), a popular web tool for gene-set enrichment analysis, aggregates by default gene-sets that reflect disease association (e.g., OMIM, Hamosh *et al.*, 2005), functional category (e.g., UniProt keywords, The UniProt Consortium, 2019), Gene Ontology (Ashburner *et al.*, 2000, The Gene Ontology Consortium, 2018), pathways (e.g., KEGG pathway, Kanehisa *et al.*, 2016) and protein domains (e.g. Interpro, Mitchell *et al.*, 2019). Additionally, users can append other gene-sets. Gene-set enrichment analysis is then performed for one set of GoI uploaded by the user, using a modified version of Fisher’s exact test running on each gene-set independently. The same approach, where gene-sets are aggregated and tested independently, is followed by most published studies and methodologies.

While some gene-sets hardly overlap with each other, others do share a substantial proportion of genes. For instance, genes associated with the keyword *chemotaxis* in UniProt are highly redundant with genes associated with the biological-process terms *cell chemotaxis* and *inflammatory response* in GO, because chemokine signalling underlie all three concepts. If a set of GoI is indeed enriched of chemokine-relevant genes, all three gene-sets are likely reported as hits. When gene-sets overlap, the independence assumption underlying statistical modelling is compromised (Tarca *et al.*, 2012; Maleki and Kusalik, 2018). Besides, the redundancy complicates human interpretation because a common set of genes underlie many hit gene-sets, some with very different names.

One way to assist interpretation is to cluster gene-sets with similar compositions *post hoc.* For instance, the DAVID Gene Functional Classification Tool is based on the kappa statistics, a similarity measure of gene-sets based on gene-set composition, and a fuzzy heuristic multiple-linkage partition algorithm to cluster gene-sets that are similar with each other into so-called *annotation clusters* (Huang *et al.*, 2007). The results are useful because they organize gene-sets reflecting identical or relevant biological aspects together, and the resolution of the clustering can be modulated by user-defined parameters. Nevertheless, users still have to examine gene-sets within each annotation cluster to derive a high-level understanding of pathways and gene-sets that are enriched. This does not scale when many annotation clusters are identified or when many sets of GoI are to be tested simultaneously.

For instance, we recently analyzed results from a community challenge to assess network module identification methods across complex diseases (Choobdar *et al.*, 2019) and had to annotate more than three hundred consensus modules that were reported. It would be slow, error-prone, and not reproducible to manually curate results of gene-set enrichment analysis to remove redundancy for such a large number of sets of GoI.

We hence asked whether it is possible to perform large-scale, parsimonious, and interpretable gene-set enrichment analysis with minimal human intervention. We wished to explicitly model redundancy between gene-sets using a single unified statistical framework. This led us to paraphrase gene-set enrichment as a regression problem. The key insight is that we can treat gene-sets as binary feature vectors of background genes with the value of one if a gene belongs to the gene-set and zero otherwise.

From this perspective, the problem of gene-set enrichment in a set of GoI can be seen as a regression problem with many correlated features (gene-sets in this case) as predictors, and the membership of GoI as a binary response variable, with the value of one if a gene belongs to the GoI and zero otherwise.

Many regularized regression algorithms are able to identify a parsimonious set of independent variables, including the *Lasso* method (Tibshirani, 1996), *Ridge* regression (Hoerl and Kennard, 1970), or linear support vector machines (Hastie *et al.*, 2004). Below we demonstrate the principle and feasibility using the elastic net (Zou and Hastie, 2005), an established statistical procedure that combines regularization and feature selection. We show that regularized regression is able to derive biologically meaningful and succinct lists of gene-sets that reflect biological functions enriched in GoI, without *post hoc* human interference.

## Algorithm and Implementation

We provide a schematic representation of the *gerr* algorithm in Figure 1 and the mathematical description below. We let *G* denote the GoI and *B* the background. We assume that *G* ⊆ *B*, *i.e.* any gene in *G* must be also in *B*. We let *Y* denote a binary vector indexed by genes in *B*: *y_g_* = 1 if and only if *g ∈ G*.

**Figure 1:**
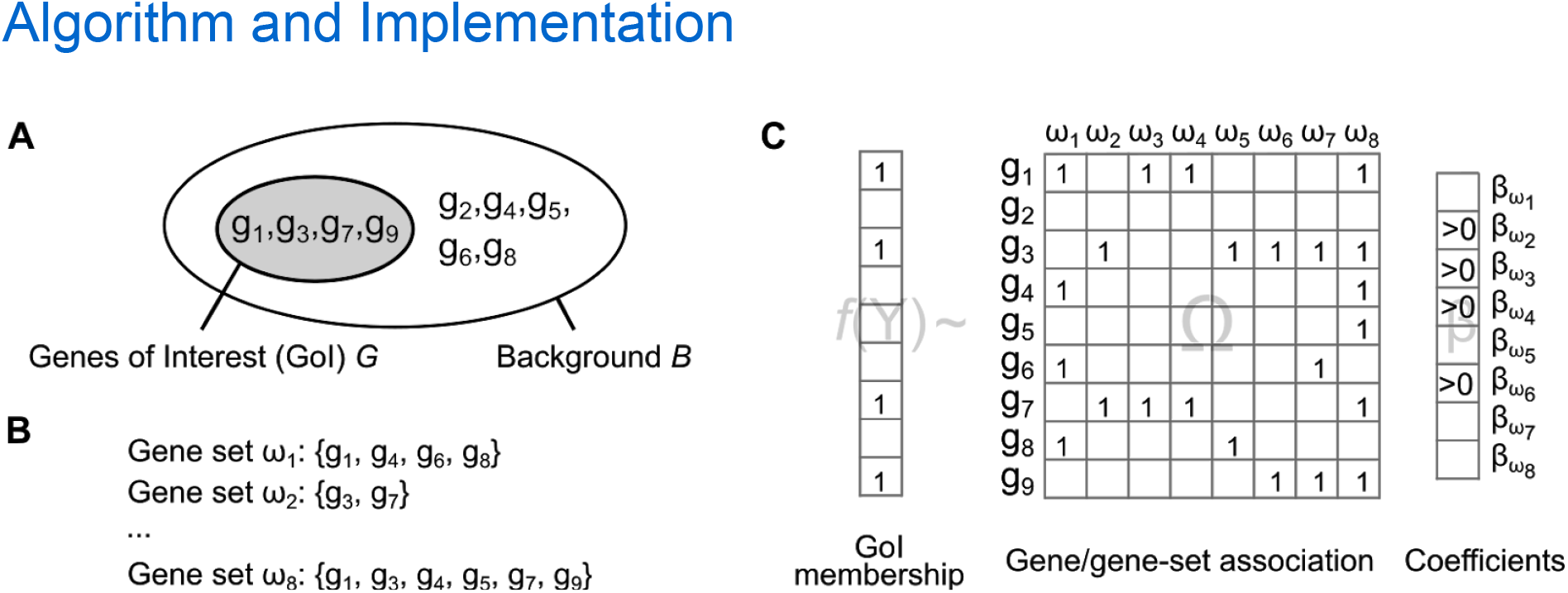
Schematic representation of the *gerr* algorithm. We present a toy example with four genes of interest in a background of nine genes (panel A), and eight gene-sets (panel B). The gene-enrichment with regularized regression (*gerr*) algorithm formulates the task of gene-set enrichment analysis as a regression problem with GoI membership as a binary target variable of background genes, and gene/gene-set association as (partially correlated) independent variables, using a link function *f* that can be specified by the user (panel C). Feature selection is achieved by identifying gene-sets with coefficients that are non-zero, more specifically that are positive for over-representation. In the toy example, gene-sets ω_2_, ω_3_, ω_4_, ω_6_ are selected. Empty cells in column vectors and in the matrix have the value of zero.

We let Ω denote the set of gene-sets that we use to annotate *G*, with *n* gene-sets ω_1_ ω_2_, …, ω_*n*_ as elements. In its simplest form, each gene-set ω_*i*_ (*i* = 1,2, …, *n*) is expressed as a binary vector indexed by genes in *B*: ω_*ig*_ = 1 if and only if *g* ∈ ω_*i*_. Equivalently, we use Ω to denote the gene/gene-set association matrix with genes in rows and gene-sets in columns.

The statistical model of generalized linear regression (Agresti, 2015) has the form of

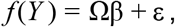

where function *f* denotes the inverse of the link function, which in the case of linear regression is the identity function, and in the case of logistic regression is the logit function that maps from the domain of [0,1] to the real-number domain *R. Y* is the column vector of GoI membership, β denotes the column vector of coefficients of gene-sets, and ε denotes the error term that is assumed to be independently and identically distributed following the normal distribution. A similar definition of the problem was used by Mi *et al.* (2012). The important distinction, however, is that we use regularization to mitigate non-independence between gene set, while Mi *et al.* applied logistic regression to rank gene-sets, not to select a relevant and non-redundant set.

Previous studies (Hellevik, 2009) and our observations (supplementary document 1) show that the results of linear models are both stable and meaningful. Therefore we use the linear model below and set it as the default in *gerr*, though the logistic model is equally valid and supported by our software.

Regularization techniques allow us to derive a parsimonious set of gene-sets. Examples include (1) *Lasso*, a type of *L1* regularization, which, loosely speaking, shrinks coefficients of less important features to zero and randomly picks one variable if two variables are identical, (2) *Ridge*, a type of *L2* regularization, which shrinks coefficients of all variables and halve the coefficient if two variables are identical, and (3) the *elastic net,* a hybrid of *L1* and *L2* regularization, controlled by the hyperparameter α. *Elastic net* combines advantages of both *Lasso* and *Ridge* approaches: it estimates non-zero values for coefficients of correlated features while setting coefficients for features of low importance exactly to zero. We use an elastic net with α =0.5 implemented in *glmnet* (Friedman *et al.*, 2010). Users can control the sparsity of results by adjusting the hyperparameter.

We set the constraint that the coefficients must be non-negative, namely, we ignore under-representation and only consider over-representation of gene-sets, where genes in a gene-set are more frequently present in GoI than random as specified by the null model. The key results are gene-sets with positive values of the coefficient β.

Practically, the gene-sets can be derived from any data source. For the purpose of demonstration, we use a union set derived from GO and the Reactome pathway database (Fabregat *et al.*, 2018).

We implement the regularized regression model, the GO and Reactome gene-set collection, as well as a number of helper functions in the open-source R package *gerr* (gene-set enrichment with regularized regression). For gene-sets identified by regression, *gerr* also returns the enrichment analysis result using Fisher’s exact test for comparison. In case that selected gene-sets are originated from GO and/or the Reactome database, which implement graph (tree-like) data structures, *gerr* returns the distance from the selected node to root nodes as well as the subtree structure, which may help users understand the biological context of the selected gene-sets. Other functionalities are documented in the vignettes of the package.

## Results

### Model verification and performance evaluation

We perform simulation studies to verify the model and evaluate the performance of *gerr.* We highlight the key concepts and results below and expose the details in supplementary documents 2 and 3.

#### Simulations with MSigDB

We use five-hundred randomly selected gene-sets from MSigDB (Subramanian *et al.*, 2005) for simulations. The gene-sets are of varying sizes, ranging from dozens to thousands of genes, and some of them share common genes.

To verify the *gerr* model, we iterate over all gene-sets and for each gene-set, we assign its member genes as GoI and use *gerr* to identify enriched gene-sets. An ideal model would report the selected gene-set as the only positive hit, which is the case for *gerr* in 88% of the five-hundred cases. In the remaining cases, *gerr* always returns the true-positive hit and no more than three false-positive hits (maximum false positive rate per simulation 3/500, or 0.006).

We perform the same analysis using the one-sided Fisher’s exact test (testing for over-presentation), with FDR-correction by the Benjamini-Hochberg method (Benjamini and Hochberg, 1995). Previous work suggests that FDR is the best choice when gene-sets are likely to be related (Khatri and Drăghici, 2005; Hackenberg and Matthiesen, 2008). This procedure, called FET+FDR hereafter, returns many more false-positive hits than *gerr* (Figure 2A, the median false positive rate per simulation 7/500, or 0.014).

**Figure 2:**
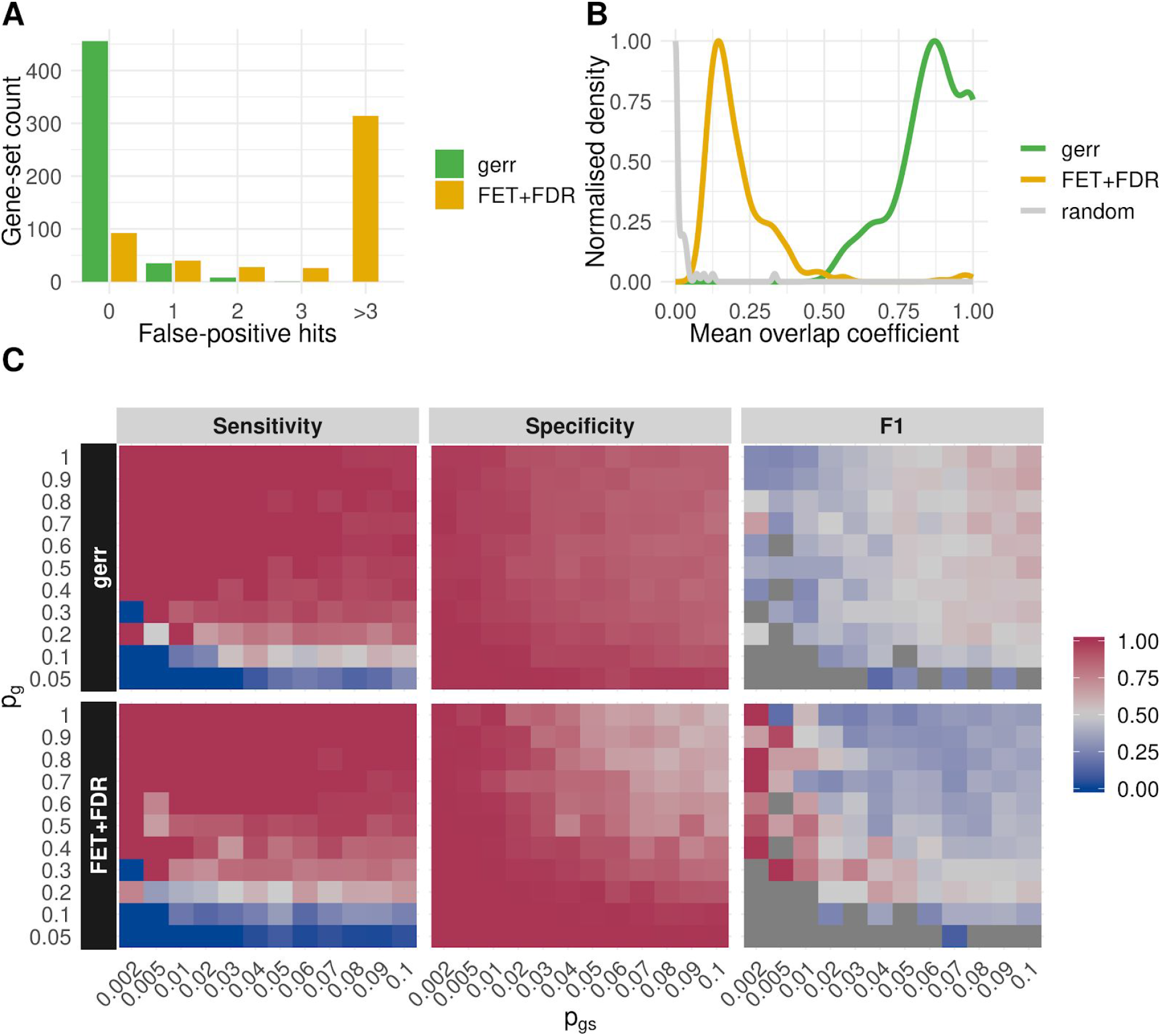
Simulation studies suggest that *gerr* offers comparable or superior performance compared with the established method, one-sided fisher’s exact test with FDR correction (FET+FDR). (A) Both *gerr* and FET+FDR achieved 100% sensitivity in model verification. However, FET+FDR returned up to hundreds of false-positive hits (median: 7), whereas *gerr* returned only up to 3 false positive hits (median: 0). (B) Mean overlap coefficient between false positive hits and the true positive hit is substantially higher for *gerr* (green) than for FET+FDR (orange) or randomly drawn gene-sets (grey). (C) Sensitivity (true positive rate, or TPR), specificity (1-false positive rate), and *F*_1_ score (harmonic mean of precision and sensitivity) of *gerr* and FET+FDR for varying combinations of parameters. The parameters include *p_gs_*, the probability of a gene-set contributing to GoI, *p_g_*, the probability of a gene in the gene-set being chosen as a member of GoI, and *p_n_*, the probability of background gene being chosen as a member of GoI. We show here the results when *p_n_* is fixed as 0.001. The results are comparable when *p_n_* varies between 0 and 0.1 (supplementary document 2).

False-positive hits of *gerr* are mainly caused by gene-sets showing high overlap with the true-positive hit. The average overlap coefficient, defined as |*A* Π *B*|/min(|*A*|, |*B*|) for sets *A* and *B*, between false positive hits and the true positive hit is substantially higher for *gerr* than for FET+FDR, and than the expected values if gene-sets are drawn randomly (Figure 2B). This suggests that *gerr* is more conservative with regard to genes in GoI that are not in the gene-set of consideration, which explains the lower false positive rates.

Taken together, model verification suggests that *gerr* achieves high sensitivity and specificity in gene-set enrichment.

Next, we evaluate the specificity, sensitivity, and precision of the *gerr* model in a probabilistic framework. We assume a generative model of GoI, where the probability of *g* ∈ *G* is modelled by 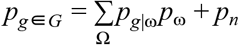, an additive model with two components. The first component specifies that first gene-sets and then member genes are randomly sampled with the probability *p*_ω_ and *p*_g|ω_, respectively. The second component specifies that background genes are chosen to constitute GoI with the probability *p_n_*, which is a gene-set independent noise term (the subscript *n* stands for *noise*). The total probability is defined between the range [0,1].

We assume that both *p*_ω_ and *p*_g|ω_ are random variables following Bernoulli distributions and that a single noise term *p*_n_ applies to all genes. Under these assumptions, we vary the values of *p*_ω_, *p*_g|ω_, and *p_n_* and assess the sensitivity, specificity, and *F*_1_ score of the *gerr* model and of the FET+FDR procedure, respectively (Figure 2C). The results suggest that *gerr* is robust against the noise component of the generative model of GoI, and shows generally higher sensitivity, specificity, and precision than FET+FDR.

In short, simulation studies suggest that *gerr* works reasonably well if a set of GoI is constructed either by genes of a single gene-set or by the proposed additive model with a noise term. In reality, genes of interest are usually generated from a much more complex model than the proposed additive model. Nevertheless, the simulation studies suggest that the performance of *gerr* may rival or even exceed the performance of well-established FET+FDR procedure.

#### Simulations with Gene Ontology

While the gene-sets used in the simulation study above are not organized in any structure, some gene-sets, such as those derived from GO, are organized in a graph structure. Under such circumstances, it is possible to reduce redundancy between enriched gene-sets using graph decorrelation (Alexa *et al.*, 2006; Grossmann *et al.*, 2007). To assess the performance of *gerr* with such gene-sets, we compare its performance with various graph-decorrelation techniques implemented in the *topGO* package (Alexa and Rahnenführer, 2019).

For these simulations, we randomly select fifty gene-sets and all their parent gene-sets from GO, which resulted in 536 gene-sets. We account for the parent-child relationships of the ontology (see details in supplementary document 3). Otherwise, the setup is identical to the simulation with MSigDB gene-sets described above.

Simulation results show that *gerr* demonstrates the highest sensitivity and specificity (Table 1). When the inclusion of genes from the selected gene-sets in GoI is stochastic, *gerr* performs comparably or better than graph-decorrelation methods (supplementary document 3). The results are remarkable because, unlike other methods in comparison, *gerr* does not use the graph-structure information. Nevertheless, *gerr* is able to cope with the high redundancy of gene-sets.

**Table 1.** True positive rates and false-positive rates in *GO* simulations, when all genes from the selected gene-set are present in GoI. Best performances are shown in bold.

| Gene-set enrichment method | True Positive Rate | False Positive Rate |
| --- | --- | --- |
| *gerr* | **1.000** | **0.00958** |
| *FET+FDR* | **1.000** | 0.134 |
| *elim* (Alexa *et al.*, 2006) | 0.424 | 0.0171 |
| *parentchild*<br>(Grossmann *et al.*, 2007) | 0.920 | 0.0809 |
| *weight* (Alexa *et al.*, 2006) | 0.494 | 0.0117 |
| *weight01* (Alexa and Rahnenführer, 2019) | 0.422 | 0.0168 |

#### A case study with consensus modules identified by a community effort in the DREAM challenge

Given the good performance of *gerr* in simulations, we further demonstrate its value on a real-world data set from a community-wide data challenge that some of us organized (the Disease Module Identification DREAM Challenge, Choobdar *et al.*, 2019). In this challenge, participating teams from all over the world applied different methods to detect disease-related gene modules from diverse human molecular networks, including protein-protein interaction, signalling, co-expression, cancer-related and homology-based networks. The collective effort of over 400 challenge participants resulted in a unique compendium of modules for the different types of molecular networks considered, from which robust consensus modules were derived that outperformed the best individual methods (Choobdar *et al.*, 2019).

While most modules partly reflect known pathways or functional gene categories, which they reorganize and expand with additional genes, other modules may correspond to yet uncharacterized pathways. The consensus modules thus constitute a novel data-driven pathway collection. However, for this community resource to be useful, we have to assign meaningful functional annotations to the identified consensus modules. To this end, we consider the consensus modules as GoI and test enrichment with all gene-sets from GO and Reactome using both FET+FDR and *gerr.*

We find that FET+FDR generally result in a large number of enriched gene-sets for each module, making it difficult to understand the characteristic functions of modules. In contrast, *gerr* allows us to readily identify much more specific functions for modules. For example, for modules derived from protein-protein interaction networks based on the STRING database (Szklarczyk *et al.*, 2017), FET+FDR returns a median of 54 GO and Reactome gene-sets per module, while *gerr* returns a median of 9 gene-sets (Figure 3A). Numbers of gene-sets returned by FET+FDR and *gerr* are positively correlated, suggesting that *gerr*, similar with *FET+FDR*, returns more gene-sets for modules with more heterogeneous functions (Figure 3B, Pearson’s correlation coefficient ρ = 0.70).

**Figure 3.**
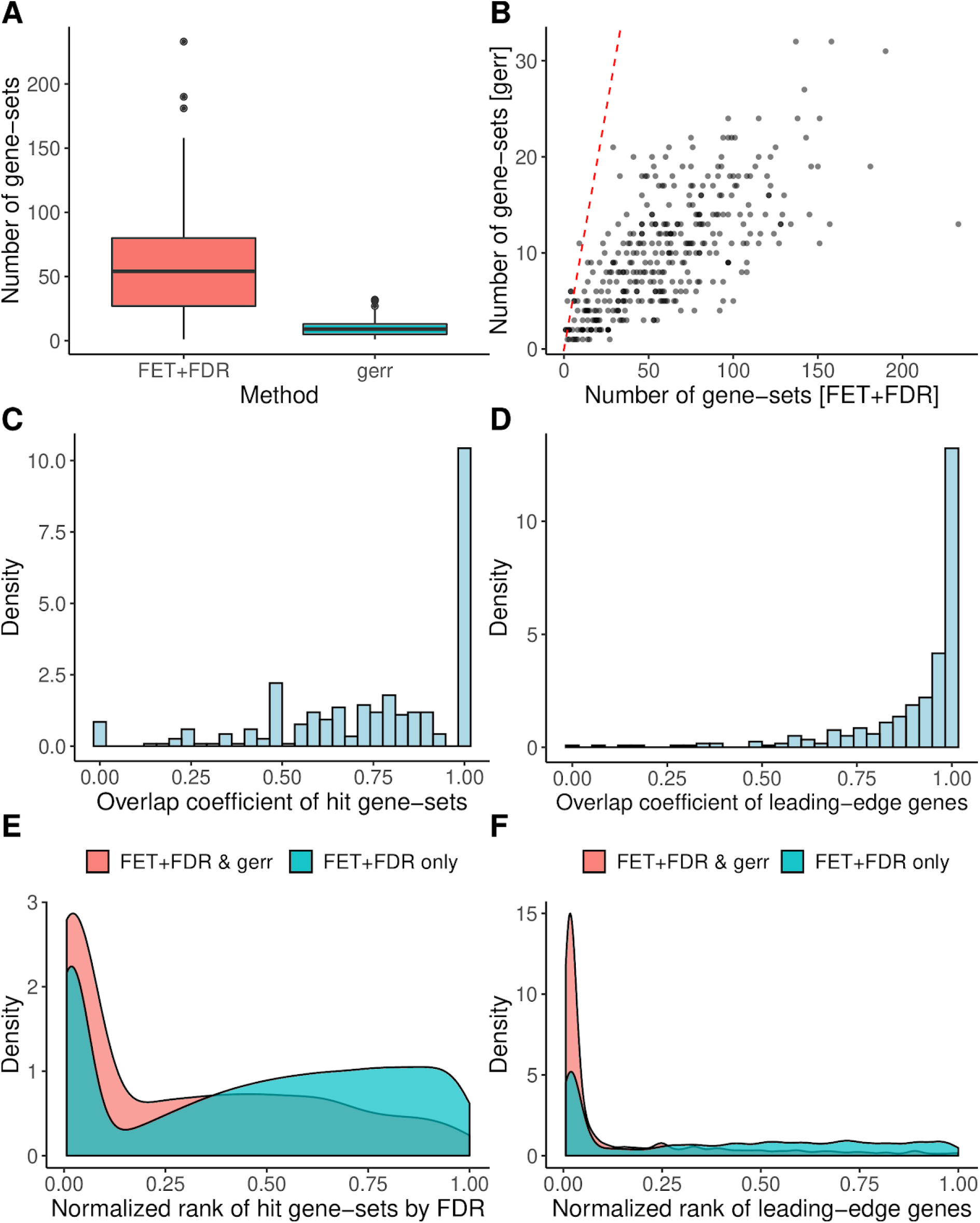
*gerr* annotates network modules identified in the DREAM challenge with parsimonious gene-sets that are consistent with the results of the FET+FDR approach. We apply both *gerr* and FET+FDR to 377 consensus disease-related gene modules detected from the STRING network (Choobdar *et al.*, 2019), and report here summary statistics. (A) Whisker-box plot of the number of hit gene-sets for each module. The horizontal bar in the box plot indicates the median value, the boxes span between the first and the third quartile, outliers that are beyond 1.5 times the interquartile range (IQR) are plotted individually as dots. (B) Scatter plot of the number of hit gene-sets returned by both methods for each module. The red dashed line represents *y = x*. (C-D) Overlap of hits identified by FET+FDR and *gerr* across modules. The distribution of overlap coefficients between FET+FDR and *gerr* hit gene-sets is shown in panel C, and that of leading-edge genes, *i.e.* genes that belong to at least one hit gene-set, is shown in panel D. The overlap coefficients are between 0.5 and 1 for most modules, suggesting that the hits returned by *gerr* are often a subset of hits returned by FET. (E) Normalised rank between 0 (top-ranking by ascending FDR values) and 1 (bottom-ranking) of gene-sets that are selected by both *gerr* and FET+FDR (red), and gene-sets that are selected by FET+FDR alone (green). Gene-sets selected by *gerr* tend to enrich towards the top of the ranking. (F) The enrichment becomes even stronger if we consider the normalised rank of genes. Leading-edge genes are ranked by the lowest FDR value of the gene-sets it belongs to. Gene-sets identified by *gerr* are enriched of genes that are associated with gene-sets with low FDR values identified by FET+FDR, which suggests that *gerr* enriches biological information despite sparse solutions.

The biological information represented by the gene-sets selected by both approaches is consistent. We assess the consistency both manually, by curating the titles and associated descriptions of the enriched gene-sets, and computationally, by examining the overlap between the GO hits of both approaches using the overlap coefficient (Figure 3C). The strong overlap for most modules suggests that *gerr* essentially captures a subset of the gene-sets identified by FET+FDR.

Next, we consider the overlap at the level of *leading-edge* genes. Following the convention of GSEA (Subramanian *et al.*, 2005), we define leading-edge genes as the genes within GoI that belong to at least one hit gene-set. We evaluate for every module the overlap coefficient of leading-edge genes identified by both approaches (Figure 3D). We find that *gerr* effectively identifies a subset of leading-edge genes identified by FTE+FDR for most modules.

Furthermore, *gerr* does not only report gene-sets that have the lowest FDR values but also report gene-sets that are not among the top hits of FET+FDR (Fig. 3E). These non-top ranking gene-sets complement the top-ranking ones in their composition and function, as visualized by the enrichment of leading-edge genes (Fig. 3F). In addition, hits of *gerr* contain higher proportions of GoI genes compared with hits of FET+FDR (supplementary document 4, figure 1). And finally, for some modules, *gerr* i dentifies gene-sets that are not selected by FET+FDR but potentially relevant for the biological function of the module (supplementary document 4, figure 2).

Taken together, our analysis of modules from the DREAM Challenge shows that *gerr* is able to identify a succinct list of gene-sets that is representative of the biological functions associated with genes of interest.

## Conclusions and discussions

In this paper, we present a novel perspective on the problem of gene-set enrichment. By considering genes of interest as a binary response variable of background genes and gene-sets as features, we transform gene-set enrichment to a regression problem, thus explicitly allowing dependencies between individual gene-sets. We propose the *gerr* algorithm and provide an implementation based on the elastic-net regularization technique in a user-friendly R package. We demonstrate the value of *gerr* both using simulated data, where it shows comparable or superior sensitivity and specificity than canonical enrichment methods. In a real-world case study, *gerr* returns parsimonious gene-sets that help us to understand the biological functions of network modules from the Disease Module Identification DREAM Challenge.

The motivation and outcome of *gerr* are quite different from established gene-set enrichment analysis techniques. Most methods seek to uncovering all gene-sets that are over- or under-presented in GoI exhaustively. In contrast, *gerr* aims at identifying a minimum set of representative gene-sets. Clearly, the two approaches are complementary and it may be desirable to apply both: while canonical methods give a comprehensive view of all enriched gene-sets, *gerr* is able to select few specific gene-sets that are most characteristic for the GoI. The latter is particularly valuable for human interpretation and comprehension of gene-set enrichment analysis results.

When many gene-sets are used, a sparse solution to the enrichment problem has been so far only possible if the gene-sets are organized in a tree structure that can be exploited by the gene-set enrichment analysis method, *e.g.* topGO (Alexa *et al.*, 2006). Otherwise, most methods assume independence between gene-sets, an assumption that is often invalid due to the ever-increasing volume of biological knowledge that is embodied in heterogeneous, partially redundant gene-sets. The *gerr* algorithm, in contrast, applies to both structured gene-sets, such as GO and Reactome, and loosely structured or unstructured gene-sets, such as those in the MSigDB database and the CREEDS database (Wang *et al.*, 2016) that are constructed by manual or semi-automatic curation of multiple datasets. Given that technologies such as single-cell omics (Sturm *et al.*, 2019), spatial omics (Rodriques *et al.*, 2019) and functional genomic screening (Haney et al., 2018; Pluvinage *et al.*, 2019) are enriching our knowledge in cell-type and cell-state-specific gene expression and function at an unprecedented rate, we envision that the number of gene-sets available for enrichment analysis will continue to grow. Most of them will be likely unstructured, at least for a period of time, necessitating methods like *gerr* that properly account for redundancy. Surprisingly, even when using gene-sets that are structured in a graph, *gerr* is able to mitigate the high correlation between gene-sets without explicitly using the graph structure.

Another major advantage of a regression approach to gene-set enrichment is that it is highly flexible and can be naturally extended to model data in diverse ways. In this work, we use linear regression to demonstrate the feasibility, though logistic regression is equally legitimate. In this vein, different types of dependent variables can be modelled by choosing an appropriate link function (McCullagh and Nelder, 1989). For instance, when more than one GoI sets are present, multinomial regression can be used to select gene-sets that are characteristic for each GoI. In addition, it is possible to model gene-level summary statistics instead of binary membership of GoI, as demonstrated by Frost and Amos (2017). Similarly, it is possible to use continuous variables instead of binary vectors as independent variables to model associations between genes and gene-sets. Last but not least, covariates can be modelled in the framework of linear regression. It is straightforward to control for biases, such as gene length, by incorporating them into the model, as demonstrated by Mi *et al.*, (2012). In summary, we believe that the great flexibility of generalized linear models (Dobson and Barnett, 2008) allows developers and users to extend the scope of *gerr* to be used in many areas of bioinformatics and genomics analysis where gene-set level interpretation is helpful.

## Acknowledgements

We thank Nikolaos Berntenis and Patrick Roelli who read earlier versions of the manuscript and made valuable suggestions. We also thank Manfred Kansy, Martin Ebeling, Fabian Birzele, Corinne Solier, and Thomas Singer for the support of the work. The work benefits from discussions within the Bioinformatics and Exploratory Data Analysis (BEDA) team. The internship of Mr Tao Fang was sponsored by the Pharmaceutical Sciences department of Roche Pharma Research and Early Development, Roche Innovation Center Basel.

