## Supplementary document 1 for "Gene-set Enrichment with Regularized Regression"

### Gene-set enrichment with regularized regression: comparing linear versus logistic regression

**May 30, 2019**

### 1 Background

In the manuscript *Gene-set Enrichment with Regularized Regression*, we propose using regularized regression to model the relationship between $Y$, a binary dependent (target) variable indicating membership of genes in a set of genes of interest (GOI hereafter), and $\Omega$, a matrix of binary variables indicating membership of genes in gene-sets that are potentially overlapping or even identical with each other.

Classically, binary target variables are often modeled by logistic regression. Alternatively, they can also be modeled by simple linear regression (Agresti 2019), even when the target variable is a dichotomy, namely either $0$ or $1$ (Hellevik 2009).

In this document, we illustrate how the two types of modelling can be constructed with `gerr`, the software package that we published along with the manuscript. In addition, we compare the results of elastic-net regression using either the linear regression or the logistic regression.

Throughout the document, we consider a small list of genes as genes of interest.

```
gene_list <- c("TRPC4AP","CDC37","TNIP1","IKBKB",
               "NKIRAS2","NFKBIA","TIMM50","RELB",
               "TNFAIP3","NFKBIB","HSPA1A","NFKBIE",
               "SPAG9","NFKB2","ERLIN1","REL",
               "TNIP2","TUBB6","MAP3K8")
```

### 2 Reguarlized linear regression

The link function of the generalized linear regression is specified by, the `family` parameter in the `glmnet` and `cv.glmnet` functions in the `glmnet` package. First, we construct a linear regression model, using the Gaussian family (`family="gaussian"`).

```
gaussRes <- regression_selected_pathways(gene_input=gene_list,
                                         family="gaussian",
                                         alpha=0.5)
```

The results of linear regression are stable, in the sense that once the random-number generator runs in a predefined state by calling `set.seed` function, running the model several times will return the same results.

```
gaussRes$selected_pathways_names
## $`R-HSA-1810476`
## [1] "RIP-mediated NFkB activation via ZBP1"
##
## $`R-HSA-5603029`
## [1] "IkBA variant leads to EDA-ID"
##
## $`R-HSA-933542`
## [1] "TRAF6 mediated NF-kB activation"
```

#### 2.1 Visualizaton of the feature-selection process of the linear model

We use the `plot` function in the `glmnet` package to visualize the feature-selection process of the model.

```
plot(gaussRes$model)
```

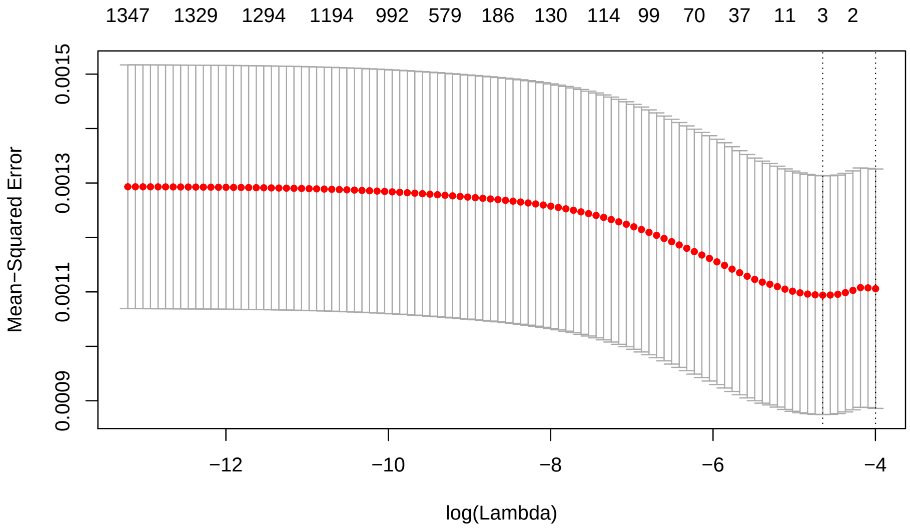

Interestingly, we observe that the mean squared error (MSE) first decreases with increasing number of features (gene-sets in this case) selected, and then increase. This can be read from the plot above, from right to the left. This is apparently different from most prediction tasks, where MSE decreases with more features are used.

I believe that this is due to the dichotomy nature of both dependent and independent variables. With more and more gene-sets are used for prediction, there will be more true-positive results, namely genes are correctly predicted to be in the set of genes of interest (GOI). However more false-positive results are also expected, namely genes outside the set of GOI are wrongly predicted to be members of GOI. Therefore, we use cross validation to select the optimal regularization parameter $\lambda$ in order to identify a minimal set of gene-sets that describe GOI.

#### 2.2 Identifying the leading-edge genes

Leading-edge genes were used in the GSEA paper to indicate genes that contribute significantly to the enrichment scores (Subramanian et al. 2005). Here, we use the term to indicate genes that are correctly identified by the regularized regression algorithm as members of GOI. These true-positive genes are so-to-say supporting the model.

By comparing the predicted `y` value (GOI membership) with the observed values, we notice that the prediction is far from perfect. Perhaps due to the sparsity of the GOI membership (~1/1000), the model is supported by a few genes within GOI while many other genes within GOI are wrongly predicted to have the same response variable as genes out of GOI.

```
gaussPred <- predict(gaussRes$model, newx=gaussRes$x, s="lambda.min")
table(predY=gaussPred, obsY=gaussRes$y)
```

```
##                       obsY
## predY                      0     1
##   0.00100161704903782 17142    15
##   0.00906961694711607     8     0
##   0.113712195513041       3     0
##   0.121780195411119       0     1
##   0.182826081733001       4     3
```

Below are the genes that are supporting the model. They apparently are associated with NF-$\kappa$B signalling pathway. This is consistent with the selected gene-sets by the regularized regression model.

```
# genes that drive the prediction using the Gaussian model
# they are apparently NF-$\kappa$B relevant genes
rownames(gaussRes$x)[which(gaussPred>0.12 & gaussRes$y==1)]
## [1] "IKBKB"  "NFKBIA" "NFKBIB" "NFKB2"
```

And below are the genes that are likely false-positives, namely these genes are not within GOI, but the model predicts otherwise. Indeed, these genes are also genes associated with the NF-$\kappa$B pathway.

```
# genes that drive the prediction using the Gaussian model
# they are apparently NF-$\kappa$B relevant genes
rownames(gaussRes$x)[which(gaussPred>0.12 & gaussRes$y==0)]
## [1] "CHUK"  "NFKB1" "RELA"  "IKBKG"
```

### 3 Reguarlized logistic regression

Instead of simple linear regression, we can also use logistic regression to model the dichotomy of the dependent variable. Though the concept is similar to the linear regression, we found that the solution is not stable. Running the model several times will give partially overlapping but different results.

It is worth mentioning though that empirical observations suggest the results are similar to the linear-regression case. And if the model is repeatedly running many times, the selected gene-sets are observed to converge towards the selected gene-sets using the simple linear regression.

```
binomRes <- regression_selected_pathways(gene_input=gene_list,
                                         family="binomial",
                                         alpha=0.5,
                                         type.measure="deviance")
length(binomRes$selected_pathways_names)
## [1] 3
```

#### 3.1 Visualization of the feature-seletion process of the logistic regression model

```
plot(binomRes$model)
```

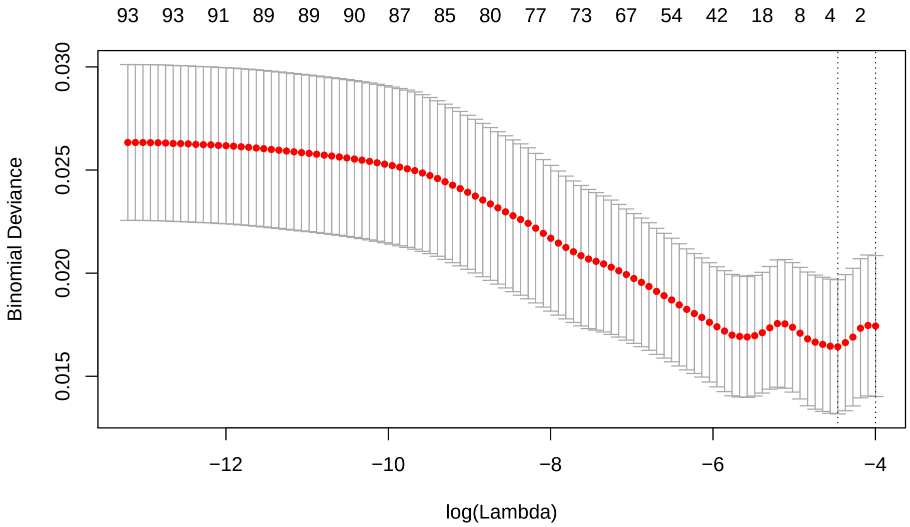

I think that the instability (at least partially) comes from the fact that the binomial deviance is not strictly convex but with two local minima (see an example above). Interestingly, if the function `cv.glmnet` is repeatedly called, after three times only one local minima is represent, where three gene-sets are identified, two out of which are identical with the Gaussian fitting results. This could potentially point to a convergence problem.

The manuscript describing the `glmnet` package (Friedman, Hastie, and Tibshirani 2010) discussed several important implementation details that may be relevant for this observation, among others given that $p >> N$, $\lambda$ cannot be run all down to zero, and that the Newton algorithm is not guaranteed to converge without step-size optimization, and the code in `glmnet` does not implement any checks for divergence. I believe that the observation of instability in the case of logistic regression deserves further study. For the current implementation, I believe it makes sense to stick to the simply linear regression for simplicity and robustness.

#### 3.2 Prediction performance and leading-edge genes

The prediction results of the logistic regression model are quite similar with those of the linear model.

```
binomPred <- predict(binomRes$model, newx=binomRes$x, s="lambda.min")
table(predY=binomPred, obsY=binomRes$y)
##                      obsY
## predY                     0     1
##   -6.87311421762438 17148    15
##   -6.51123053631526     2     0
##   -2.59527225002329     3     1
##   -1.82759996003652     4     3
```

Indeed, the same set of leading-edge genes are identified.

```
# genes that drive the prediction using the binomial model
# they are apparently NF-$\kappa$B relevant genes
rownames(binomRes$x)[which(binomPred>(-3) & binomRes$y==1)]
## [1] "IKBKB"  "NFKBIA" "NFKBIB" "NFKB2"
```

And below are the genes that are likely false-positives, namely these genes are not within GOI, but the model predicts otherwise. Note that beyond the genes associated with the NF-$\kappa$B pathway that were identified by the linear model, a few other genes, including *RIPK3*, *RIPK1*, and *ZBP1*, are reported as well, which have been also reported to be associated with or relevant for the NF-$\kappa$B pathway (for instance see (Yatim et al. 2015)).

```
# genes that drive the prediction using the binomial model
# they are apparently NF-$\kappa$B relevant genes
rownames(binomRes$x)[which(binomPred>(-3) & binomRes$y==0)]
## [1] "CHUK"  "NFKB1" "RELA"  "IKBKG" "RIPK3" "RIPK1" "ZBP1"
```

### 4 Conclusions

In this document, we show how to construct regularized linear and logistic regression models using the `gerr` package, and compare the results of the two types of models.

We observed that the linear regression provides stable solutions that are consistent with the results of regularized logistic regression. This was not only the case for the examples shown above; we tested multiple sets of genes of interest and made similar observations.

Based on these results, in the current implementation of the `gerr` package, we use the cross-validation version of the elastic-net regression using the `gaussian` family (linear regression) as the default option, though the `binomial` family can be specified by the user as well.

### 5 R session info

```
sessionInfo()
## R version 3.6.0 (2019-04-26)
## Platform: x86_64-pc-linux-gnu (64-bit)
## Running under: Linux Mint 18.3
##
## Matrix products: default
## BLAS:   /usr/lib/openblas-base/libblas.so.3
## LAPACK: /usr/lib/libopenblasp-r0.2.18.so
##
## locale:
##  [1] LC_CTYPE=en_GB.UTF-8       LC_NUMERIC=C
##  [3] LC_TIME=en_GB.UTF-8        LC_COLLATE=en_GB.UTF-8
##  [5] LC_MONETARY=de_CH.UTF-8    LC_MESSAGES=en_GB.UTF-8
##  [7] LC_PAPER=de_CH.UTF-8       LC_NAME=C
##  [9] LC_ADDRESS=C               LC_TELEPHONE=C
```

```
## [11] LC_MEASUREMENT=de_CH.UTF-8 LC_IDENTIFICATION=C
##
## attached base packages:
## [1] stats     graphics  grDevices utils     datasets  methods   base
##
## other attached packages:
## [1] glmnet_2.0-18    foreach_1.4.4    Matrix_1.2-17    gerr_0.99.19
## [5] BiocStyle_2.13.0
##
## loaded via a namespace (and not attached):
##  [1] Rcpp_1.0.1         bookdown_0.10      codetools_0.2-16
##  [4] lattice_0.20-38    digest_0.6.18      grid_3.6.0
##  [7] magrittr_1.5       evaluate_0.13      stringi_1.4.3
## [10] rmarkdown_1.12     iterators_1.0.10   tools_3.6.0
## [13] stringr_1.4.0      igraph_1.2.4.1     xfun_0.7
## [16] yaml_2.2.0         compiler_3.6.0     pkgconfig_2.0.2
## [19] BiocManager_1.30.4 htmltools_0.3.6    knitr_1.22
```

### References

Agresti, Alan. 2019. *An Introduction to Categorical Data Analysis*. 3rd Edition. Wiley.

Friedman, Jerome, Trevor Hastie, and Rob Tibshirani. 2010. "Regularization Paths for Generalized Linear Models via Coordinate Descent." *Journal of Statistical Software* 33 (1): 1–22. https://www.ncbi.nlm.nih.gov/pmc/articles/PMC2929880/.

Hellevik, Ottar. 2009. "Linear Versus Logistic Regression When the Dependent Variable Is a Dichotomy." *Quality & Quantity* 43 (1): 59–74. doi:10.1007/s11135-007-9077-3.

Subramanian, Aravind, Pablo Tamayo, Vamsi K. Mootha, Sayan Mukherjee, Benjamin L. Ebert, Michael A. Gillette, Amanda Paulovich, et al. 2005. "Gene Set Enrichment Analysis: A Knowledge-Based Approach for Interpreting Genome-Wide Expression Profiles." *Proceedings of the National Academy of Sciences* 102 (43): 15545–50. doi:10.1073/pnas.0506580102.

Yatim, Nader, Hélène Jusforgues-Saklani, Susana Orozco, Oliver Schulz, Rosa Barreira da Silva, Caetano Reis e Sousa, Douglas R Green, Andrew Oberst, and Matthew L Albert. 2015. "RIPK1 and Nf-$\kappa$B Signaling in Dying Cells Determines Cross-Priming of Cd8+ T Cells." *Science* 350 (6258): 328–34.
