## Supplementary document 2 for "Gene-set Enrichment with Regularized Regression"

### Gene-set enrichment with regularized regression: simulation studies

**June 04, 2019**

### 1 Background

In the manuscript *Gene-set Enrichment with Regularized Regression* (gerr in short), we propose using regularized regression to model the relationship between $Y$, a dichotomous dependent variable indicating membership of genes in a set of genes of interest (GOI hereafter), and $\Omega$, a matrix of dichotomous variables indicating membership of genes in gene-sets that are potentially overlapping or even identical with each other.

In this document, we perform simulation studies to demonstrate the sensitivity and specificity of the gerr method, using the software package of the same name that we published along with the manuscript and the elastic net implemented in the R software package glmnet. Throughout the analysis, we will use the default parameters of gerr, using the elastic net of Gaussian-family linear regression, with $\alpha = 0.5$.

### 2 Gene-sets used for simulations

We wish to use real-world gene-sets that are commonly used by the community for the simulation study for gerr, because synthesized gene-sets may have distributions of sizes, defined by the number of unique genes and overlapping patterns that depart from real-world gene-sets. Therefore we decided to use curated gene-sets provided by the MSigDB database (Category C2) for this purpose. These gene-sets are curated from online pathway databases, publications in PubMed, and collected knowledge of domain experts. Therefore they are not structured in tree structures like gene-sets from Gene Ontology (GO) or from the Reactome pathway database.

```
# gerrRootDir is the root directory of the gerr package
simGenesetsRdata <- file.path(gerrRootDir, "data/simGenesets.RData")
if(!file.exists(simGenesetsRdata)) {
  library(msigdbr)
  simTibble <- msigdbr::msigdbr(species="Homo sapiens",
                                category = c("C2"))
  simGenesAll <- with(simTibble, tapply(human_gene_symbol, gs_name, unique))
  simGenesets <- sample(simGenesAll, 500)
  save(simGenesets, file=simGenesetsRdata)
} else {
  load(simGenesetsRdata)
}
simLen <- sapply(simGenesets, function(x) length(x))

# bgGenes is a vector of unique background genes
bgGenes <- unique(unlist(simGenesets))
# gsMatrix is a gene by gene-set matrix
gsMatrix <- Matrix(sapply(simGenesets, function(x) as.integer(bgGenes %in% x)),
                   sparse=TRUE,
                   dimnames=list(bgGenes, names(simGenesets)))
```

The gene-sets were retrieved using the msigdbr package. For the purpose of simulation, we randomly sample 500 gene-sets from all gene-sets of the category C2 (N=500). We resave the data in the gerr package so that future updates in either the MSigDB database or the msigdbr package will affect the simulation results.

#### 2.1 Distribution of gene-set sizes

The histogram below shows the distribution of gene-set size in the sub-sampled set of gene-sets.

```
{
  hist(simLen, xlab="Number of unique genes",
       breaks=50, freq = FALSE,
       col="lightblue", main="Gene-set size distribution")
  lines(density(simLen, from=0), col="#004495", lwd=2)
}
```

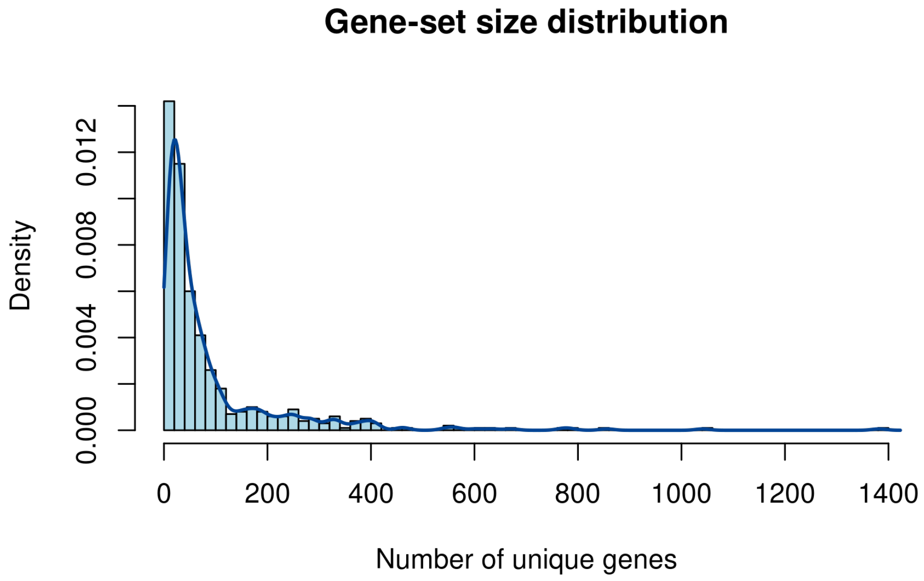

The median gene-set size is 38, with heavy tail on the right side.

#### 2.2 Distribution of pairwise overlap coefficients between gene-sets

Next, we investigate the degree of redundancy among these gene-sets. We use the overlap coefficient, defined by $|A \cap B|/min(|A|, |B|)$ between two sets $A$ and $B$, to measure this.

When the gene-sets are represented as a dichotomous matrix, pairwise overlap coefficients can be calculated efficiently by using matrix operations. The functions below are extracted from the ribiosUtils package, which is currently being submitted to the CRAN repository.

```
uniqueNonNA <- function(x) {
  x <- x[!is.na(x)]
  res <- unique(x)
  return(res)
}
columnOverlapCoefficient <- function(x, y=NULL) {
  if(!is.matrix(x)) x <- as.matrix(x)
  if(is.null(y)) y <- x
  if(is.matrix(y)) y <- as.matrix(y)
```

```
  stopifnot(nrow(x)==nrow(y))
  storage.mode(x) <- storage.mode(y) <- "integer"

  tmatProd <- t(x) %*% y
  xCount <- apply(x, 2, function(xx) sum(xx!=0))
  yCount <- apply(y, 2, function(yy) sum(yy!=0))
  tmatPmin <- outer(xCount, yCount, pmin)
  res <- tmatProd/tmatPmin
  if(is.null(y)) {
    diag(res) <- 1L
  }
  dimnames(res) <- list(colnames(x), colnames(y))
  return(res)
}
listOverlapCoefficient <- function(x, y=NULL, checkUniqueNonNA=TRUE) {
  if(checkUniqueNonNA) {
    x <- lapply(x, uniqueNonNA)
    if(!is.null(y)) {
      y <- lapply(y, uniqueNonNA)
    }
  }

  if(is.null(y)) {
    elements <- unique(unlist(x))
    mat <- sapply(x, function(xx) as.integer(elements %in% xx))
    res <- columnOverlapCoefficient(mat)
  } else {
    elements <- unique(c(unlist(x), unlist(y)))
    mat1 <- sapply(x, function(xx) as.integer(elements %in% xx))
    mat2 <- sapply(y, function(xx) as.integer(elements %in% xx))
    res <- columnOverlapCoefficient(mat1, mat2)
  }
  return(res)
}
```

With these functions, we can calculate pairwise overlap coefficients between gene-sets.

```
simPairwiseOverlap <- columnOverlapCoefficient(gsMatrix)
utOver <- simPairwiseOverlap[upper.tri(simPairwiseOverlap, diag=FALSE)]
{
  hist(utOver, xlab="Pairwise overlapping cofficient between gene-sets",
       breaks=50, freq = FALSE,
       col="orange",
       main="Overlapping coefficient distribution")
  lines(density(utOver, from=0), col="red", lwd=2)
}
```

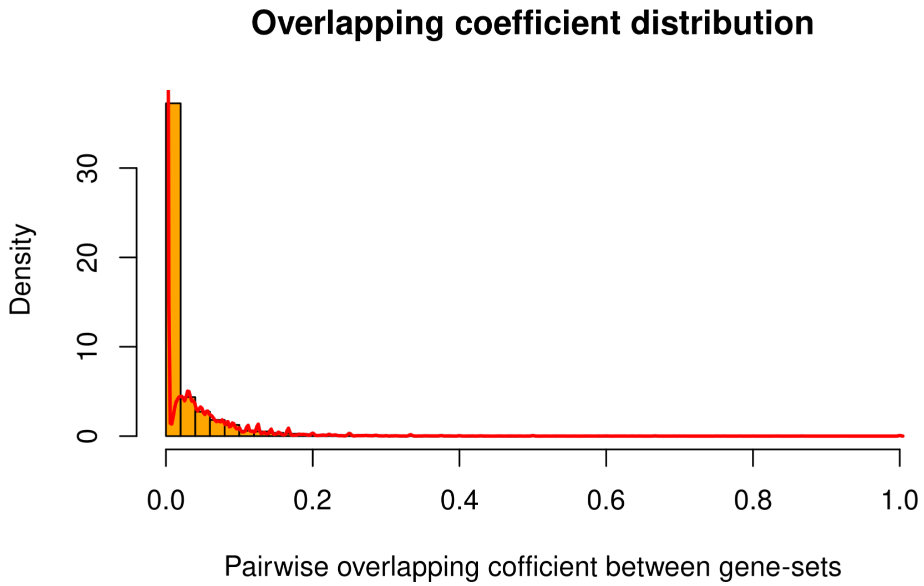

We calculated the overlap coefficient for all unique pairs of gene-sets, and its distribution is shown in the histogram above. While no overlapping genes are identified between 69.8% pairs of gene-sets, overlapping genes are found in 30.2% pairs of gene-sets, with an median overlap coeffiecient of 0.047 and median absolute deviation (MAD) of 0.035. In about 0.5% of the cases, the overlap coefficient is equal or larger than 0.5. It suggests that while the assumption of independence between gene-sets may hold true when few gene-sets are used, there will be violations when many gene-sets are used for testing.

### 3 Model verification

We first verify that gene-set enrichment with regularized regression (`gerr`) performs as expected using the simplest simulation that is possible: we assign genes of one gene-set as GOI, and test whether we can recover the gene-set using `gerr`.

#### 3.1 A small-scale verification with few cases

In the code chunk below, we verify that the model works in the sense that the gene-set that is used as source of GOI is correctly recovered, using randomly selected 10 gene-sets.

```
set.seed(1)
selInd <- sample(seq(along=simGenesets), 10)
selGsName <- names(simGenesets)[selInd]
selGs <- simGenesets[selInd]
selRes <- lapply(selGs, function(gs)
  regression_selected_pathways(gene_input=gs, gene_pathway_matrix = gsMatrix))
foundCounts <- sapply(seq(along=selInd),
                      function(i) length(selRes[[i]][[1]]))
isRecovered <- sapply(seq(along=selInd),
                      function(i) selGsName[i] %in% names(selRes[[i]][[1]]))
stopifnot(all(isRecovered))
```

```
table(foundCounts)
## foundCounts
## 1 2
## 8 2
```

The `foundCounts` variable indicates the number of gene-sets whose coeffient is positive. In this particular example, we have 8 cases where only one gene-set is selected, and thanks to the observation that the input gene-set is all recovered (`isRecovered` is all true), the selected gene-set by `gerr` must be the input geneset.

There are two cases where two gene-sets are returned. The code below examine how the selected gene-sets are related.

```
mt1Return <- foundCounts>1
mt1ReturnOverlapCoef <- sapply(which(mt1Return), function(i) {
  selGnames <- names(selRes[[i]]$selected_pathways_names)
  poe <- listOverlapCoefficient(simGenesets[selGnames])
  mean(poe[upper.tri(poe, diag=FALSE)])
})
print(mt1ReturnOverlapCoef)
## [1] 0.5757576 0.7083333
```

It turns out in both cases, the selected gene-sets (one of which is the input gene-set) are partially redudant, with an overlap coefficient higher than 0.5. This makes sense because when two dependent variables are similarly correlated with the target variable, they are given similar weights by the elastic net.

#### 3.2 The full-scale verification with all gene-sets

Below we run the simulation for all gene-sets. This is not executed by default because of running time (about four minutes in a Linux in Virtual environment with 4G memory and one core Intel i5 CPU), but the idea is the same as the small verification step above.

```
verificate <- function(genesets, gsMatrix, index) {
  stopifnot(index %in% seq(along=genesets))
  selGsName <- names(genesets)[index]
  selGs <- genesets[index]
  selRes <- lapply(selGs, function(gs) {
    regression_selected_pathways(gene_input=gs, gene_pathway_matrix = gsMatrix)
  })
  selInd <- sapply(seq(along=index),
                   function(i) selRes[[i]][[1]])
  isRecovered <- sapply(seq(along=index),
                   function(i) selGsName[i] %in% names(selRes[[i]][[1]]))
  res <- list(results=selRes, selInd=selInd, isRecovered=isRecovered)
  return(res)
}
```

```
modelVerifRdata <- file.path(gerrRootDir,
                             "data/modelVerification.RData")
if(!file.exists(modelVerifRdata) || params$overwriteCache) {
```

```
  system.time(verifRes <- verificate(simGenesets,
                                     gsMatrix,
                                     index=seq(along=simGenesets)))
  verifPaths <- lapply(verifRes$results,
                       function(x) x$selected_pathways_names)
  verifKeyRes <- list(isRecovered = verifRes$isRecovered,
                      selInd = verifRes$selInd,
                      selected_pathways_names=verifPaths)
  save(verifKeyRes, file=modelVerifRdata)
} else {
  load(modelVerifRdata)
}
```

We first confirm that each gene-sets used as a GOI was successfully rediscovered by `gerr`.

```
all(verifKeyRes$isRecovered) ## all recovered
## [1] TRUE
```

Next we query the frequency of cases where `gerr` returned more than one gene-set.

```
verifSelCount <- sapply(verifKeyRes$selInd, length)
## in 44 cases of 500 gene-sets, more than one gene-set was selected
summary(verifSelCount>1)
##    Mode   FALSE    TRUE
## logical     456      44
```

We found that in 44 cases out of 500, one than one gene-set is returned by `gerr`. The gene-sets that were not used as GOI but selected by `gerr` may be regarded as *false positive hits*. However, when we examine the similarity of these gene-sets and the true positive hit using the overlap coefficient (and the average when more than one false-positive hits are available), we found that these *false-positive hits* are in fact very similar compared with the true-positive hit. And the similarity is much higher than what is expected when a random set of gene-sets are chosen and their average overlap coefficient is calculated.

As a comparison, we run one-sided Fisher's exact test and adjust the p-values by Benjamini-Hochberg method (the FET+FDR procedure) using the same strategy, setting one gene-set as GOI a time.

First we implement the FET+FDR stratgy.

```
fetFdr <- function(goi, bgGenes) {
  goi <- uniqueNonNA(goi)
  bgGenes <- uniqueNonNA(bgGenes)
  bgDiffGoi <- setdiff(bgGenes, goi)
  fetRes <- sapply(simGenesets, function(x) {
     selHits <- length(intersect(x, goi))
     nonselHits <- length(goi) - selHits
     selNonhits <- length(x) - selHits
     nonselNonhits <- length(bgDiffGoi) - selNonhits
     mat <- matrix(c(selHits, nonselHits, selNonhits, nonselNonhits),2,2)
     res <- stats::fisher.test(mat, alternative = "greater")$p.value
  })
  ff <- p.adjust(fetRes, "BH") ## we denote fetFdr from here on as ff
```

```
  return(ff)
}
```

Next we run the verification step using FET+FDR.

```
fetFdrVerifRdata <- file.path(gerrRootDir,
                             "data/fetFdrVerification.RData")
if(!file.exists(fetFdrVerifRdata) || params$overwriteCache) {
  fisherVerif <- lapply(seq(along=simGenesets), function(i) {
    goi <- simGenesets[[i]]
    ffRes <- fetFdr(goi, bgGenes)
    isHit <- ffRes<0.05
    hits <- names(simGenesets)[isHit]
    resp <- ffRes[hits]; names(resp) <- hits
    if(length(resp)>1) {
      cosel <- setdiff(hits, names(simGenesets)[i])
      poe <- listOverlapCoefficient(list(goi),
                                    simGenesets[cosel])
      meanPoe <- mean(poe)
    } else{
      meanPoe <- NA
    }
    res <- list(hits=resp, meanPoe=meanPoe)
    return(res)
  })
  save(fisherVerif, file=fetFdrVerifRdata)
} else{
  load(fetFdrVerifRdata)
}
```

It is easy to confirm that FET+FDR has a sensitivity of 100%, namely all gene-sets used as input of GOI are rediscovered.

```
all(sapply(seq(along=simGenesets),
           function(i) names(simGenesets)[i] %in% names(fisherVerif[[i]]$hits)))
## [1] TRUE
```

We first compare the number of false-positive hits of both methods.

```
gerrFpCounts <- verifSelCount - 1
fetFdrFpCounts <- sapply(fisherVerif, function(x) length(x$hits)-1)
verifFpCounts <- data.frame(Geneset=names(simGenesets),
                            method=factor(rep(c("gerr", "FET+FDR"),
                                              each=length(simGenesets)),
                                          levels=c("gerr","FET+FDR")),
                            FP=c(gerrFpCounts, fetFdrFpCounts))

fetFdrCol <- "#E6AB02"
randomCol <- "#CCCCCC"
gerrCol <- "#4DAF4A"
veriFpPlot <- ggplot(verifFpCounts,
       aes(x=cut(FP, c(-Inf,0,1,2,3,Inf),
```

```
                 labels=c("0", "1", "2", "3", ">3")),
           fill=method)) +
  scale_fill_manual(values=c(gerrCol, fetFdrCol),
                    name="") +
  geom_bar(position=position_dodge2(preserve="single")) +
  xlab("False-positive hits") +
  ylab("Gene-set count") +
  theme(axis.text.x=element_text(angle = 0))
print(veriFpPlot)
```

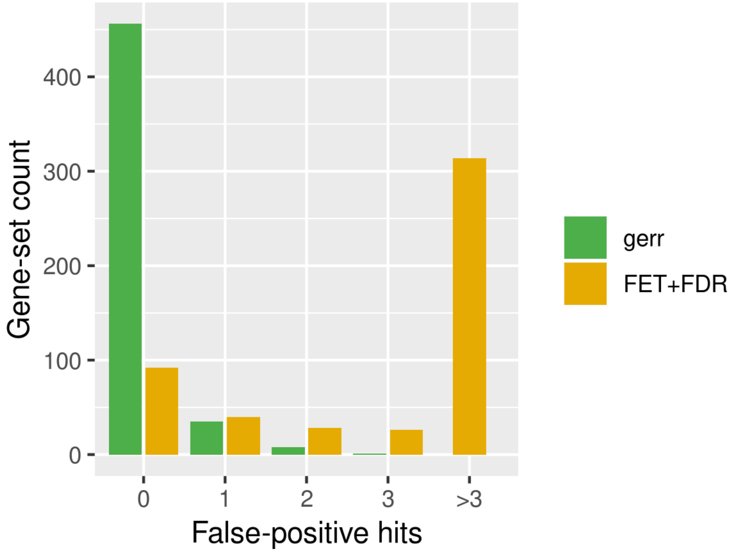

The gerr returned no false-positive hits in 91.2% cases, and otherwise only returned no more than two false-positive hits. In contrast, FET+FDR very frequently returned a large number of false-positive hits (median 7, mad 10.3782).

We believe that the false-positive hits of gerr are likely caused by redundancy in gene-sets. Therefore we show below the mean overlap coefficient between false-positive and true-positive hits returned by gerr. As a comparison, we show the value also for hits returned by FET+FDR and for randomly selected gene-sets.

```
densityDf <- function(x, from=0, to=1) {
  den <- density(x, from=from, to=to)
  res <- data.frame(x=den$x,
                    y=den$y/max(den$y))
  return(res)
}
verifMt1AvgOe <- sapply(which(verifSelCount>1), function(i) {
  inputGs <- names(simGenesets)[i]
  selGnames <- setdiff(names(verifKeyRes$selected_pathways_names[[i]]),
                       inputGs)
  poe <- listOverlapCoefficient(simGenesets[inputGs],
                                simGenesets[selGnames])
  mean(poe)
})
verifMt1AvgOeDenData <- densityDf(verifMt1AvgOe)
```

```
randomAvgOe <- sapply(which(verifSelCount>1), function(i) {
  selNo <- length(verifKeyRes$selected_pathways_names[[i]])
  randomGs <- sample(names(simGenesets), selNo, replace=TRUE)
  poe <- listOverlapCoefficient(simGenesets[randomGs])
  mean(poe[upper.tri(poe, diag=FALSE)])
})
randomAvgOeDenData <- densityDf(randomAvgOe)
fisherAvgOe <- na.omit(sapply(fisherVerif, function(x) x$meanPoe))
fisherAvgOeDenData <- densityDf(fisherAvgOe)
oeDen <- rbind(cbind(Class="gerr", verifMt1AvgOeDenData),
               cbind(Class="FET+FDR", fisherAvgOeDenData),
               cbind(Class="random", randomAvgOeDenData))
oeDen$Class <- factor(oeDen$Class,
                      levels=c("gerr", "FET+FDR", "random"))
oeDenPlot <- ggplot(oeDen,
       aes(x=x, y=y, color=Class)) +
  scale_colour_manual(values=c(gerrCol,
                               fetFdrCol,
                               randomCol),
                      name="") +
  geom_line(lwd=1.5)  +
  ylab("Normalised density") +
  xlab("Mean overlap coefficient")
print(oeDenPlot)
```

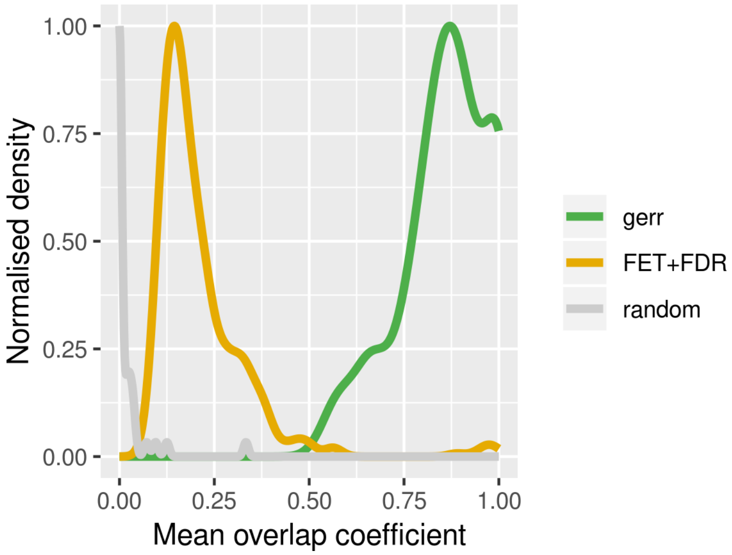

The average pairwise overlap coefficient between false-positive hits and the true positive hit is very high (median 0.8775, MAD 0.096369), and in comparison much higher than the mean overlap coefficient between false and true positive hits returned by FET+FDR and between randomly selected gene-sets. Such *false-positive hits* are selected by `gerr` because the elastic net algorithm wil give half of the weight to two identical features. While under certain circumstances they complicate results interpretation (but much less so compared with the results of the FET+FDR procedure), in other contexts gene-sets with similar but not identical sets of genes may be potentially interesting for end users since they may reflect relevant but not identical biological functions.

In short, with a very simple verification procedure, we verify that by using `gerr`, all gene-sets that were assigned as genes of interest were successfully recovered. In most cases, the only gene-set selected by `gerr` was the true positive hit. In other cases, more than one gene-set was selected by `gerr`. The co-selected gene-sets are highly redudant with the true positive hit.

### 4 Simulations based on a probabilistic model

Next we test the performance of `gerr` using a probabilistic framework. Specifically, we assume a generative model of GOI in the following form:

$$p_{g \in G} = \sum_{\omega} p_{g|\omega} p_{\omega} + p_{g_n}$$

.

The model specifies that the probability that a gene $g$ is a member of the set $G$ is modelled by the probability of the gene-set $\omega$ contributes to GOI, expressed as $p_\omega$, multiplied by the probability that $g$ is selected to contributed to GOI given that $\omega$ contributes to GOI, summed over all genet-sets, and then adding the term $p_{g_n}$ that models the probability that the gene $g$ contributes to GOI independent of its associations with gene-sets. The two parts on the right side of the equation can be observed as a gene-set dependent and a noise term (therefore the subscript $n$) respectively. Apparently, $p_{g \in G}$ should be located between $[0, 1]$.

In the following simulations, we assume that $p_\omega$ follows a binomial distribution that is specified by the parameter $p_{gs}$. For simplicity, we model $p_{g|\omega}$ as a binomial distribution specified by the parameter $p_g$ and assume that the same parameter applies to all selected gene-sets. Furthermore, we assume that $p_{g_n}$ is modelled by a small number $p_n$ that varies in a given range, e.g. from $10^{-4}$ to $10^{-1}$, and the value applies to all genes.

The simulation procedure can be described in following steps:

1. Randomly sample $k$ gene-sets from $\Omega$, the set of gene-sets, with the probability of $p_{gs}$.
2. Within each $k$ gene-sets, randomly select $m$ genes each with the probability of $p_g$.
3. Randomly sample genes from the background $B$ by a uniform distribution with the parameter $p_n$.
4. Merge genes selected in step (2) and (3) into a set of GOI.
5. Perform the `gerr` method with $G$ and $\Omega$ as input
6. Assess the specificity and sensitivity of the `gerr` method.

The codes below implement the logic.

```
probSim <- function(p_gs=0.01, p_g=0.5, p_n=1E-3, seed=1L) {
  set.seed(seed)
  ## step 1
  gsSel <- rbinom(n=length(simGenesets), size=1, prob=p_gs) == 1
  gsSelNames <- names(simGenesets)[gsSel]
  ## step 2
  gsGenes <- lapply(simGenesets[gsSel], function(genes) {
    selGenes <- rbinom(n=length(genes), size=1, prob=p_g)
    res <- genes[selGenes==1]
```

```
    return(res)
  })
  ## step 3
  isNoiseGenes <- rbinom(n=length(bgGenes), size=1, prob=p_n) == 1
  noiseGenes <- bgGenes[isNoiseGenes]
  ## step 4
  probGoi <- unique(c(unlist(gsGenes), noiseGenes))
  ## step 5
  probRes <- regression_selected_pathways(probGoi,
                                          gene_pathway_matrix = gsMatrix)
  ## step 6
  probSelGs <- names(probRes$selected_pathways_names)

  probTp <- intersect(probSelGs, gsSelNames)
  probFp <- setdiff(probSelGs, gsSelNames)
  probFn <- setdiff(gsSelNames, probSelGs)
  probTn <- setdiff(names(simGenesets),
                    c(probTp, probFp, probFn))
  ## True-positive rate = Sensitivity
  probTpr <- length(probTp)/length(gsSelNames)
  probFpr <- length(probFp)/(length(probFp) + length(probTn))
  probPpv <- length(probTp)/(length(probTp) + length(probFp))
  probF1 <- 2 * probPpv * probTpr / (probPpv + probTpr)
  probMeanOe <- listOverlapCoefficient(simGenesets[probFp],
                                       simGenesets[probTp])
  ## special cases where there are no positives
  nTP <- length(probTp)
  nFP <- length(probFp)
  if(nTP==0 || nFP==0) {
    medTpFpMaxOe <- NA
  } else if (nTP==1 || nFP==1) {
    medTpFpMaxOe <- mean(probMeanOe)
  } else   {
    medTpFpMaxOe <- median(apply(probMeanOe, 2, max))
  }

  ## compare with Fisher's exact test + FDR (ff)
  ff <- fetFdr(goi=probGoi, bgGenes=bgGenes)
  ffSelGs <- names(which(ff<=0.05))

  ffTp <- intersect(ffSelGs, gsSelNames)
  ffFp <- setdiff(ffSelGs, gsSelNames)
  ffFn <- setdiff(gsSelNames, ffSelGs)
  ffTn <- setdiff(names(simGenesets),
                  c(ffTp, ffFp, ffFn))
  ## True-positive rate = Sensitivity
  ffTpr <- length(ffTp)/length(gsSelNames)
  ffFpr <- length(ffFp)/(length(ffFp) + length(ffTn))
  ffPpv <- length(ffTp)/(length(ffTp) + length(ffFp))
  ffF1 <- 2 * ffPpv * ffTpr / (ffPpv + ffTpr)
```

```
  res <- c(p_gs=unname(p_gs), p_g=unname(p_g), p_n=unname(p_n),
           TPR.gerr=probTpr,
           FPR.gerr=probFpr,
           PPV.gerr=probPpv,
           F1.gerr=probF1,
           nTP.gerr=nTP,
           nFP.gerr=length(probFp),
           nTN.gerr=length(probTn),
           nFN.gerr=length(probFn),
           medTpFpMaxOe.gerr = medTpFpMaxOe,
           TPR.ff=ffTpr,
           FPR.ff=ffFpr,
           PPV.ff=ffPpv,
           F1.ff=ffF1,
           nTP.ff=nTP,
           nFP.ff=length(ffFp),
           nTN.ff=length(ffTn),
           nFN.ff=length(ffFn))
  return(res)
}
```

#### 4.1 A small-scale simulation with the probabilistic model

For instance, below we example the results of the particular simulation setting.

```
testSim <- probSim(p_gs=0.05, p_g=0.5, p_n=1E-3)
print(testSim[c("TPR.gerr", "TPR.ff",
                "FPR.gerr", "FPR.ff",
                "F1.gerr", "F1.ff",
                "medTpFpMaxOe.gerr")])
##          TPR.gerr           TPR.ff          FPR.gerr            FPR.ff
##        1.00000000       0.92000000        0.08631579        0.16000000
##           F1.gerr            F1.ff medTpFpMaxOe.gerr
##        0.54945055       0.37096774        0.18181818
```

The results can be interpreted as follows.

- The true-positive rate (TPR) of the `gerr` method is 1.0, namely all gene-sets that were selected to contribute to GOI are successfully recovered. By comparison, TPR of one-sided Fisher's exact test and Benjamini-Hochberg FDR correction (FET+FDR) is 0.92.
- The false-positive rate (FPR) of the `gerr` method is 0.086, namely among all gene-sets that were *not* selected to contribute to GOI, about 9% of them are selected by `gerr`. In contrast, the FPR of FET+FDR is about 0.16.
- $F_1$ score, the harmonic mean of precision and sensitivity, of the `gerr` method is higher than that of FET+FDR.
- The value `medTpFpMaxOe` deserves explanation. We know that if two gene-sets share common genes, they may be both selected by the regularized regression procedure as relevant features, even if only one of the two gene-sets was selected to contribute to GOI. That means the false-positive hits may make still sense. To quantify this aspect,

we calculated pairwise overlap coefficients between true and false positive hits, and report the median value of maximum overlap coefficient of each false positive hit. The larger the number is, the stronger is the evidence that the redundancy between gene-sets may cause false-positive hits.

#### 4.2 A full scale simulation with the probabilistic model and results interpretation

Below we perform the full-scale simulation with the probabilistic model, using a variety of combinations of the parameters $p_{gs}$, $p_g$, and $p_n$. For each parameter set, five independent simulations are run.

```
p_gs_cand <- c(0.002, 0.005, 0.01,
               seq(0.02, 0.1, by=0.01))
p_g_cand <- c(0.05, seq(0.1, 1, by=0.1))
p_n_cand <- c(0, 1E-4, 1E-3, 1E-2, 5E-2, 1E-1)
probSimParamsOneRep <- expand.grid(p_gs_cand, p_g_cand, p_n_cand)
## five replicates per condition
probSimParams <- do.call(rbind, list(probSimParamsOneRep,
                                     probSimParamsOneRep,
                                     probSimParamsOneRep,
                                     probSimParamsOneRep,
                                     probSimParamsOneRep))
probSimParams <- cbind(probSimParams, seed=1:nrow(probSimParams))
probSimRdata <- file.path(gerrRootDir,
                          "data/modelProbSimulation.RData")
if(!file.exists(probSimRdata) || params$overwriteCache) {
  system.time(fullProbSimRes <- apply(probSimParams, 1, function(x) {
    probSim(p_gs=x[1], p_g=x[2], p_n=x[3], seed=x[4])
  }))
  fullProbSimResDf <- as.data.frame(t(fullProbSimRes))
  save(fullProbSimResDf, file=probSimRdata)
} else {
  load(probSimRdata)
}
```

Due to the stochastic nature of sampling, in some runs no gene-set is selected at all to contribute to GOI, especially when $p_{gs}$ is set small. These cases are excluded from the analysis below, because they will distort the results. Including them however do not change the conclusions.

The median values of the five simulation runs are reporteed for each measure (true-positive rate, false-positive rate, *etc.*).

```
posFullProbSimResDf <- with(fullProbSimResDf,
                            fullProbSimResDf[nTP.gerr+nFN.gerr!=0,])
avgFullProbSimRes <- aggregate(posFullProbSimResDf,
                               list(posFullProbSimResDf$p_gs,
                                    posFullProbSimResDf$p_g,
                                    posFullProbSimResDf$p_n), median)
```

### 5 Interpretation of the simulation results

We investigate the simulation results by visualizing true positive rate, false positive rate, and $F_1$ scores of the `gerr` and FET+FDR procedure.

#### 5.1 Sensitivity, or true positive rate (TPR)

The plot below visualizes how sensitivity, or true positive rate (TPR) varies by the probability that each gene-set contributes to GOI ($p_{gs}$) and the probability of genes in each gene-set contribute to GOI ($p_g$), conditional on the noise probability $p_n$. Five independent simulations were performed for each parameter set, and the median value is used for visualization.

```
lowCol <- "#004495"
midCol <- "#CCCCCC"
highCol <- "#AA3555"
theme_update(axis.text.x = element_text(angle=45, hjust=1),
        axis.text = element_text(size=11),
        axis.title = element_text(size=14),
        strip.text = element_text(size=11),
        legend.text = element_text(size=11))
tprGerr <- ggplot(avgFullProbSimRes,
                  aes(x=factor(p_gs), y=factor(p_g), fill=TPR.gerr)) +
  facet_wrap(~p_n, labeller = label_both) +
  geom_tile() +
  scale_fill_gradient2(low=lowCol, mid=midCol, high=highCol,
                       midpoint=0.5,
                       name="TPR") +
  ggtitle("True positive rate of gerr")+
  xlab(expression(p[gs])) + ylab(expression(p[g]))
print(tprGerr)
```

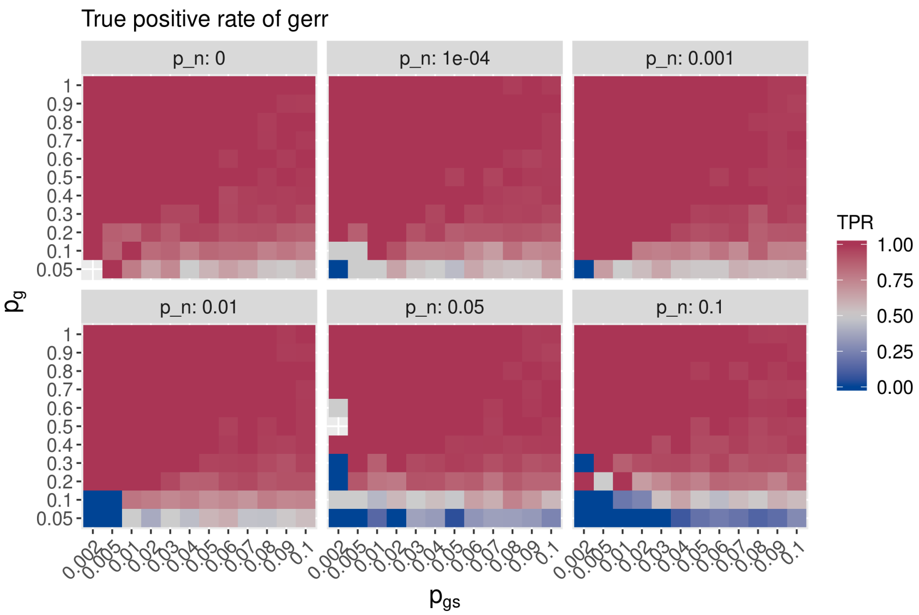

It seems that `gerr` in general has high sensitivity, even when the noise probabiliy is as high as 0.1 (namely each gene has the probability of 0.1 to be selected as a gene of interest, independent whether it is associated with any gene-set or not). The sensitivity is only low when very few genes in the gene-set contribute to GOI (say less than 10%), which makes sense intuitively.

The cell $p_{gs} = 0.002, p_g = 0.5, and p_n = 0.05$ is missing because in five runs of simulation, the sampling procedure did not pick any gene-set to contribute to contribute to GOI.

The plot below visualizes the pattern of TPR in case of the FET+FDR procedure.

```
tprFF <- ggplot(avgFullProbSimRes,
                aes(x=factor(p_gs), y=factor(p_g), fill=TPR.ff)) +
  facet_wrap(~p_n, labeller = label_both) +
  geom_tile() +
  scale_fill_gradient2(low=lowCol, mid=midCol, high=highCol,  midpoint=0.5,
                       name="TPR") +
  theme(axis.text.x = element_text(angle=45, hjust=1)) +
  ggtitle("True positive rate of FET+FDR") +
  xlab(expression(p[gs])) + ylab(expression(p[g]))
print(tprFF)
```

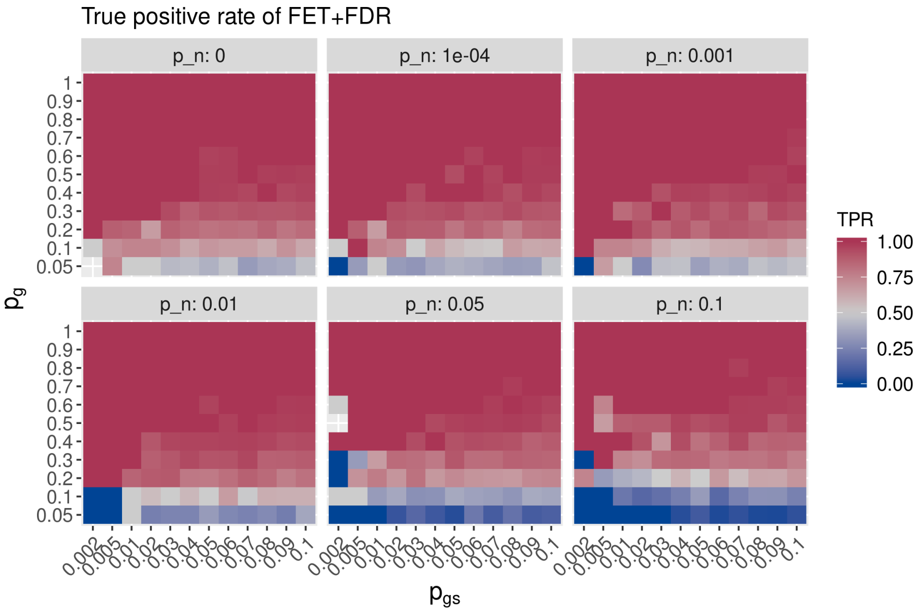

The difference of TPR between `gerr` and `FET+FDR` of matching parameter set is visualized by the plot below.

```
diffLowCol <- "#E6AB02"
diffMidCol <- "#CCCCCC"
diffHighCol <- "#4DAF4A"
tprDiff <- ggplot(avgFullProbSimRes, aes(x=factor(p_gs), y=factor(p_g), fill=TPR.gerr-TPR.ff)) +
  facet_wrap(~p_n, labeller = label_both) +
  geom_tile() +
  scale_fill_gradient2(low=diffLowCol, mid=diffMidCol, high=diffHighCol, midpoint=0,
                       limits=c(-0.5, 0.5), oob=scales::squish,
                       name=expression(Delta~TPR)) +
```

```
  theme(axis.text.x = element_text(angle=45, hjust=1)) +
  ggtitle("True positive rate difference (gerr - FET+FDR)") +
  xlab(expression(p[gs])) + ylab(expression(p[g]))
print(tprDiff)
```

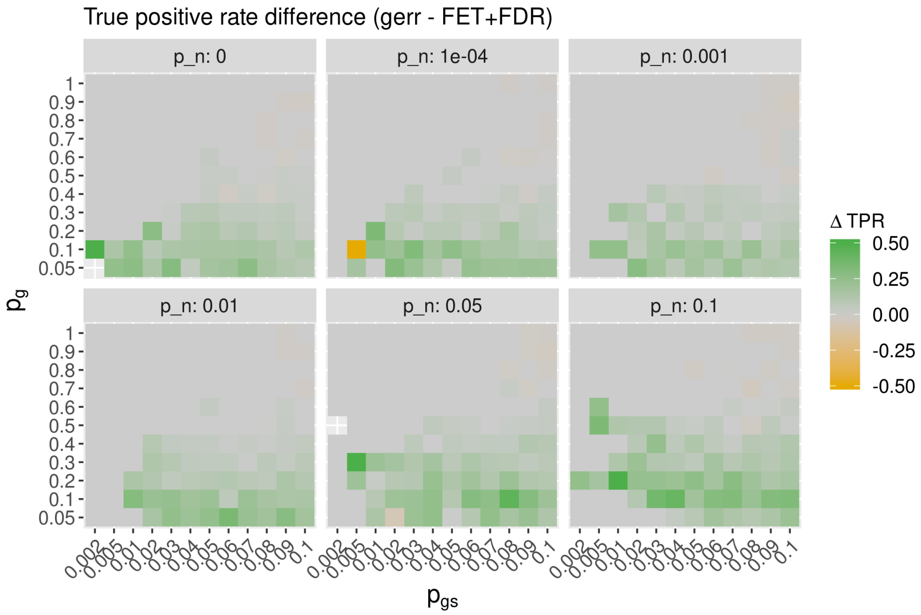

It seems that when $p_g$, the probability of genes in a gene-set to contribute to GOI, is low, and when the noise $(p_n)$ is strong, `gerr` has higher sensitivity (true-positive rate) than FET+FDR. Otherwise, when $pg$ is high, both `gerr` and FET+FDR has high sensitivity that is near 1.

#### 5.2 Specificity, or 1-false positive rate (FPR)

Next we examine specificity, which equals 1-false-positive rate First, we visualize the variation precision of `gerr` by varying parameters.

```
fprGerr <- ggplot(avgFullProbSimRes,
                  aes(x=factor(p_gs), y=factor(p_g), fill=1-FPR.gerr)) +
  facet_wrap(~p_n, labeller = label_both) +
  geom_tile() +
  scale_fill_gradient2(low=lowCol, mid=midCol, high=highCol,
                       name="1-FPR",
                       midpoint = 0.5,
                       limits=c(0,1), oob=scales::squish) +
  theme(axis.text.x = element_text(angle=45, hjust=1)) +
  ggtitle("False positive rate of gerr") +
  xlab(expression(p[gs])) + ylab(expression(p[g]))
print(fprGerr)
```

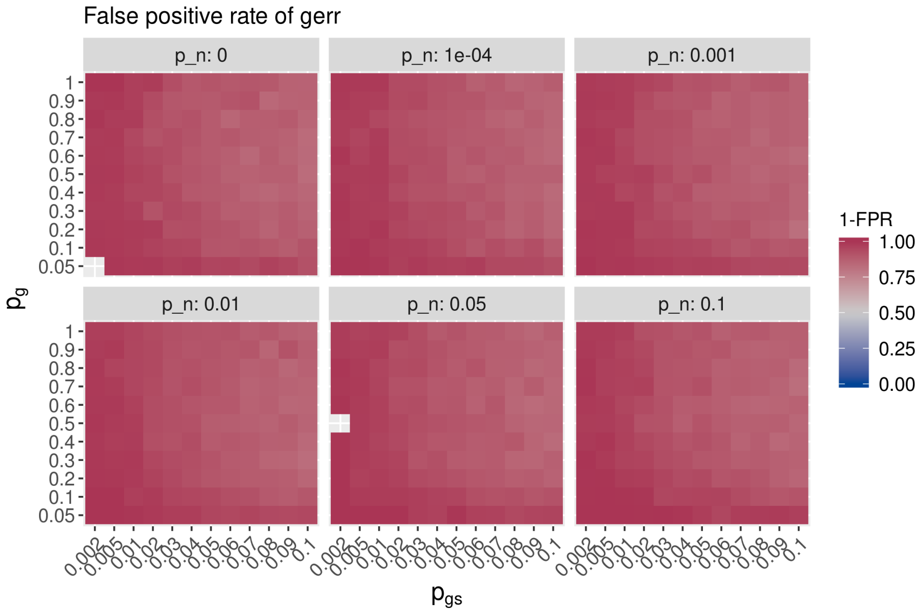

The specificity of `gerr` seems quite robust against the choice of $p_n$. It decreases when $p_{gs}$ is high, namely when many gene-sets contribute to GOI. Likely it is because of calling other gene-sets that are partially redundant as false-positive hits, as shown in the previous verification step.

Next we reveal how FPR change in the case of FET+FDR.

```
fprFF <- ggplot(avgFullProbSimRes,
                aes(x=factor(p_gs), y=factor(p_g), fill=1-FPR.ff)) +
 facet_wrap(~p_n, labeller = label_both) +
  geom_tile() +
  scale_fill_gradient2(low=lowCol, mid=midCol, high=highCol,
                       name="1-FPR",
                       midpoint = 0.5,
                       limits=c(0,1), oob=scales::squish) +
  theme(axis.text.x = element_text(angle=45, hjust=1)) +
  ggtitle("False positive rate of FET+FDR") +
  xlab(expression(p[gs])) + ylab(expression(p[g]))
print(fprFF)
```

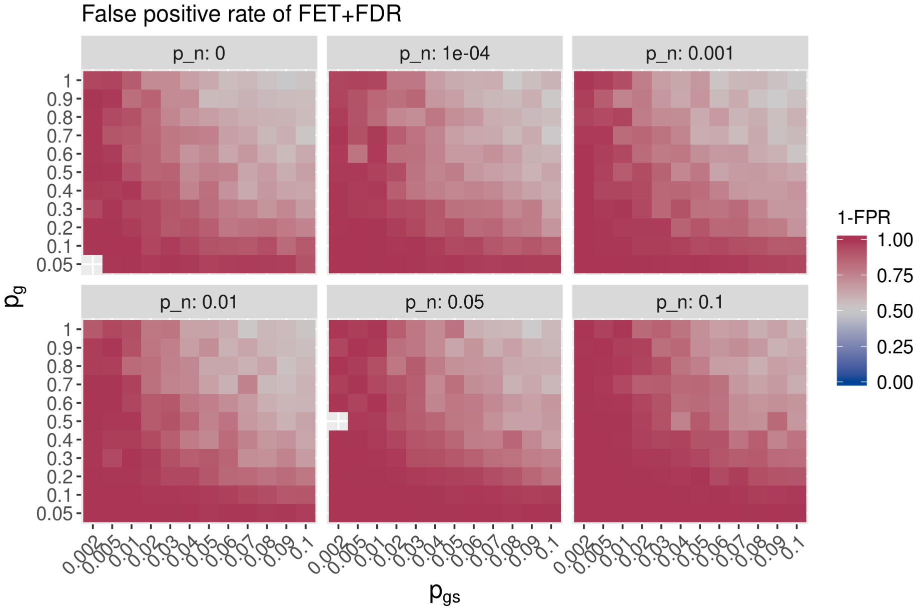

The plot below visualizes the difference of FPR.

```
fprDiff <- ggplot(avgFullProbSimRes,
                  aes(x=factor(p_gs), y=factor(p_g), fill=(1-FPR.gerr) - (1-FPR.ff))) +
  facet_wrap(~p_n, labeller = label_both) +
  geom_tile() +
  scale_fill_gradient2(low=diffLowCol, mid=diffMidCol, high=diffHighCol,
                       midpoint = 0,
                       limits=c(-0.5, 0.5), oob=scales::squish,
                       name=expression(Delta~"(1-FPR)")) +
  theme(axis.text.x = element_text(angle=45, hjust=1)) +
  ggtitle("False positive ratedifference: gerr - FET+FDR") +
  xlab(expression(p[gs])) + ylab(expression(p[g]))
print(fprDiff)
```

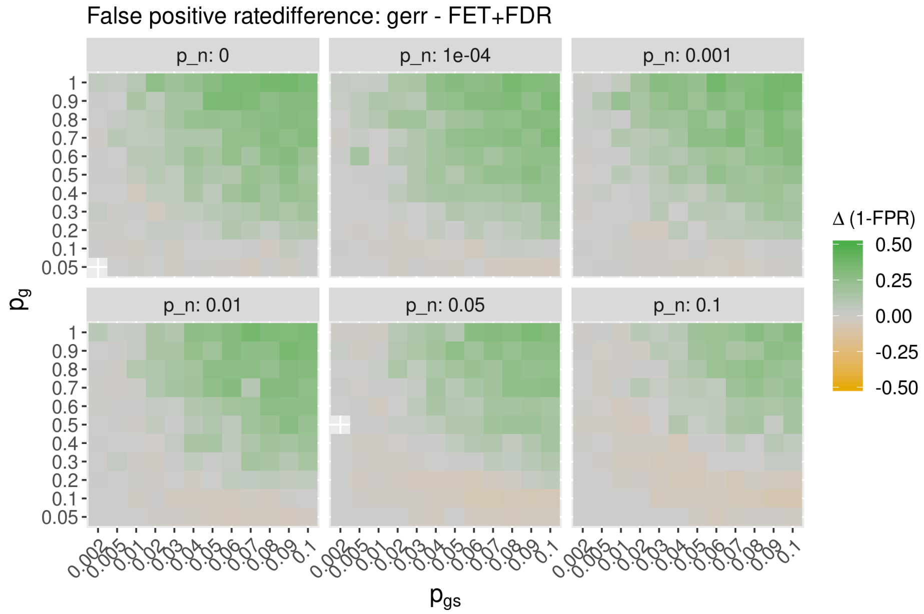

We observe that when $p_g$ is low, FET+FDR has slightly lower FPR than `gerr` (on the cost of having slightly lower sensitivty). Otherwise, gerr has much lower false-positive rate than the FET+FDR procedure. The higher $p_{gs}$ and $p_g$ values are, the stronger are the difference in favor of `gerr`.

#### 5.3 $F_1$ score

$F_1$ score is the harmonic mean of precision and sensitivity and therefore a good measure of balanced performance. Below we visualize the $F_1$ score of `gerr` results.

```
f1Gerr <- ggplot(avgFullProbSimRes,
                 aes(x=factor(p_gs), y=factor(p_g), fill=F1.gerr)) +
  facet_wrap(~p_n, labeller = label_both) +
  geom_tile() +
  scale_fill_gradient2(low=lowCol, mid=midCol, high=highCol,
                       name=expression(F[1]~score),
                       limits=c(0,1), midpoint = 0.5,
                       oob=scales::squish) +
  theme(axis.text.x = element_text(angle=45, hjust=1)) +
  ggtitle("F1 score of gerr") +
  xlab(expression(p[gs])) + ylab(expression(p[g]))
print(f1Gerr)
```

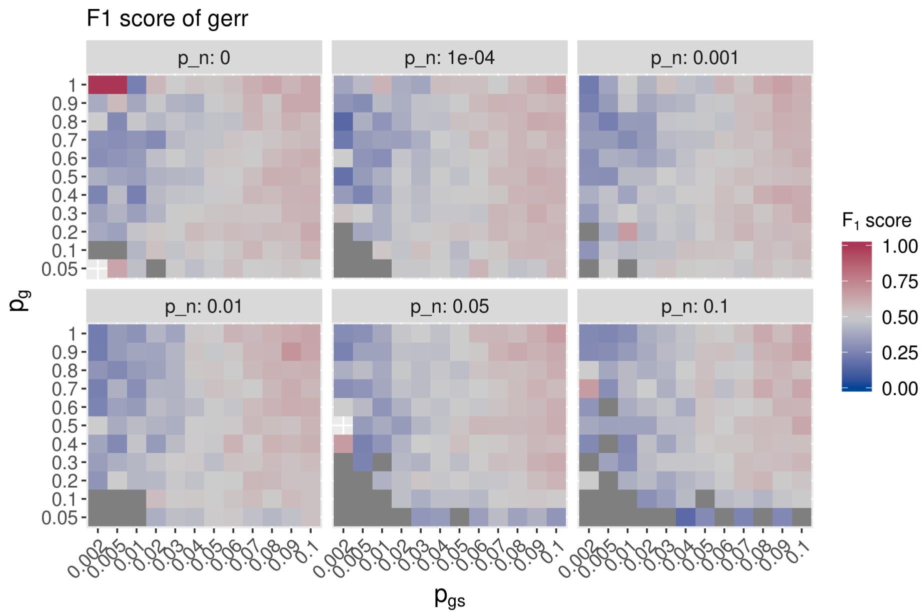

It seems that $F_1$ score of `gerr` in quite robust against $p_n$, as long as $p_{gs}$ and $p_g$ is reasonably high.

Next we visualize the $F_1$ scores of the FET+FDR procedure.

```
f1FF <- ggplot(avgFullProbSimRes,
               aes(x=factor(p_gs), y=factor(p_g), fill=F1.ff)) +
  facet_wrap(~p_n, labeller = label_both) +
  geom_tile() +
  scale_fill_gradient2(low=lowCol, mid=midCol, high=highCol,
                       name=expression(F[1]~score),
                       limits=c(0,1), midpoint=0.5,
                       oob=scales::squish) +
  theme(axis.text.x = element_text(angle=45, hjust=1)) +
  ggtitle("F1 score of FET+FDR") +
  xlab(expression(p[gs])) + ylab(expression(p[g]))
print(f1FF)
```

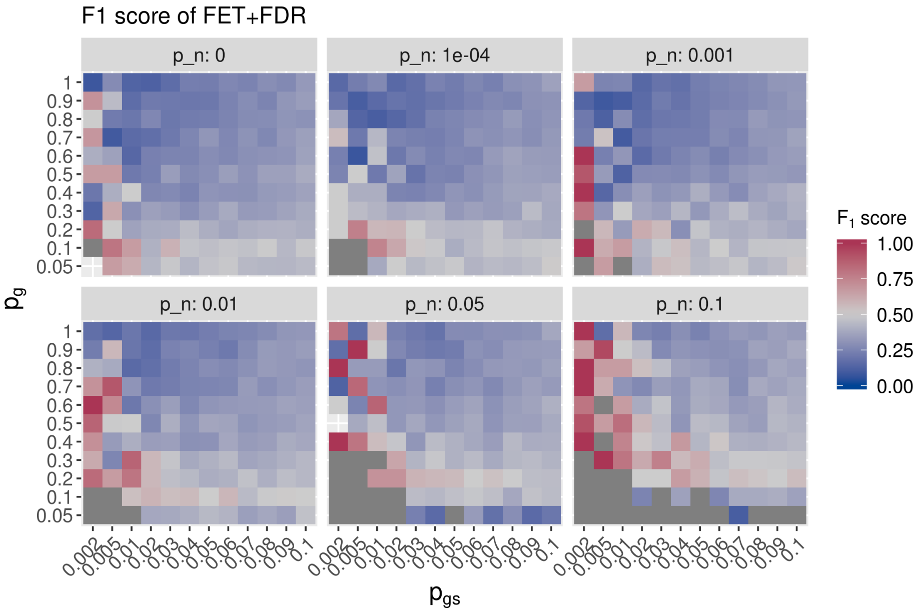

Next we calculate the difference of $F_1$ scores.

```
f1Diff <- ggplot(avgFullProbSimRes,
                 aes(x=factor(p_gs), y=factor(p_g), fill=F1.gerr - F1.ff)) +
  facet_wrap(~p_n, labeller = label_both) +
  geom_tile() +
  scale_fill_gradient2(low=diffLowCol, mid=diffMidCol, high=diffHighCol,
                       midpoint = 0,
                       name=expression(Delta~F[1]~score),
                       limits=c(-0.5, 0.5), oob=scales::squish) +
  theme(axis.text.x = element_text(angle=45, hjust=1)) +
  ggtitle("F1 score difference: gerr - FET+FDR") +
  xlab(expression(p[gs])) + ylab(expression(p[g]))
print(f1Diff)
```

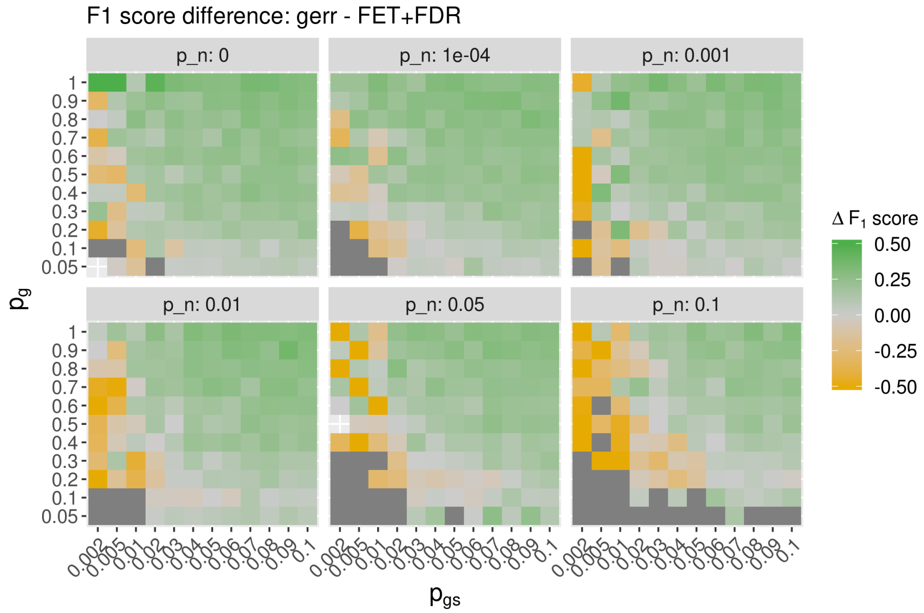

We observe that $F_1$ score, which combines precision $(TP/(TP+FP))$ and sensitivity (TPR), of FET+FDR is higher when few gene-sets are selected to construct GOI; otherwise, the score of `gerr` is higher. This is because when only few gene-sets are selected, `gerr` tend to find other gene-sets that are partially overlapping with the true positive hits, therefore its precision can be lower. Though as discussed before, this does not necessarily mean that the results of `gerr` is inferior, since the partially redundant gene-sets may be of interest under circumstances. Otherwise, when $p_{gs}$ is reasonably high (above 0.01), $F_1$ score of `gerr` is generally higher than FET+FDR. Namely when not few gene-sets contribute to GOI, and when many genes of selected gene-sets are chosen to construct GOI, `gerr` outperforms FET+FDR by $F_1$ score.

### 6 Results of model verification and performance evaluation in one figure

Finally, we merge key results of the simulation studies into one figure. We show the particular case of $p_g = 0.01$. The conclusions however are valid for other settings of $p_g$.

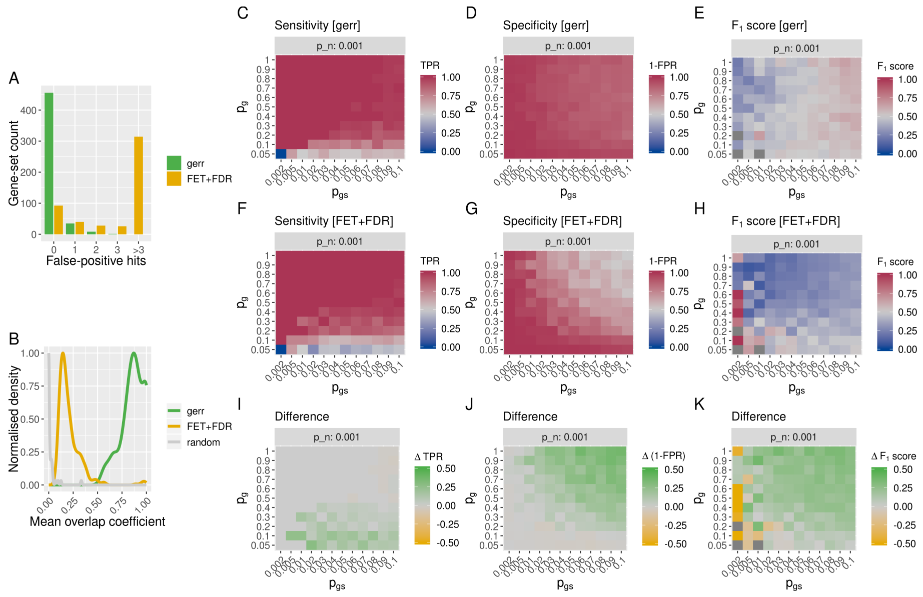

### 7 Conclusions

In this document, we first characterized the gene-sets (a subset of MSigDB C2 category gene-sets) that we used for the simulation study. Next, we verificated that our model performs as expected by using gene-sets directly as genes of interest and recovering them. Last but not least, we used a generative probabilistic model to generate genes of interest by sampling gene-sets and genes within selected gene-sets, and by adding a noise term.

Model verification and simulations with the probabistic model show that gene-set enrichment with regularized regression (`gerr`) is a method with high sensitivity and specificity, and is robust with regard to noise in the set of genes of interest. Within the parameter space of our simulation, `gerr` outperforms Fisher's exact test and FDR correction in situations where the genes of interest are contributed by many gene-sets and/or where genes within a gene-set have a large probability to contribute to GOI.

### 8 R session info

```
sessionInfo()
## R version 3.6.0 (2019-04-26)
## Platform: x86_64-pc-linux-gnu (64-bit)
## Running under: Linux Mint 18.3
##
## Matrix products: default
## BLAS:   /usr/lib/openblas-base/libblas.so.3
## LAPACK: /usr/lib/libopenblasp-r0.2.18.so
##
## locale:
##  [1] LC_CTYPE=en_GB.UTF-8       LC_NUMERIC=C
```

```
##  [3] LC_TIME=en_GB.UTF-8        LC_COLLATE=en_GB.UTF-8
##  [5] LC_MONETARY=de_CH.UTF-8    LC_MESSAGES=en_GB.UTF-8
##  [7] LC_PAPER=de_CH.UTF-8       LC_NAME=C
##  [9] LC_ADDRESS=C               LC_TELEPHONE=C
## [11] LC_MEASUREMENT=de_CH.UTF-8 LC_IDENTIFICATION=C
##
## attached base packages:
## [1] grid      stats     graphics  grDevices utils     datasets  methods
## [8] base
##
## other attached packages:
##  [1] gridExtra_2.3    dplyr_0.8.1      scales_1.0.0     ggplot2_3.1.1
##  [5] MASS_7.3-51.4    glmnet_2.0-18    foreach_1.4.4    Matrix_1.2-17
##  [9] gerr_0.99.19     BiocStyle_2.13.0
##
## loaded via a namespace (and not attached):
##  [1] Rcpp_1.0.1         compiler_3.6.0     pillar_1.4.0
##  [4] BiocManager_1.30.4 plyr_1.8.4         iterators_1.0.10
##  [7] tools_3.6.0        digest_0.6.18      evaluate_0.13
## [10] tibble_2.1.1       gtable_0.3.0       lattice_0.20-38
## [13] pkgconfig_2.0.2    rlang_0.3.4        igraph_1.2.4.1
## [16] yaml_2.2.0         xfun_0.7           withr_2.1.2
## [19] stringr_1.4.0      knitr_1.22         tidyselect_0.2.5
## [22] glue_1.3.1         R6_2.4.0           rmarkdown_1.12
## [25] bookdown_0.10      purrr_0.3.2        magrittr_1.5
## [28] codetools_0.2-16   htmltools_0.3.6    assertthat_0.2.1
## [31] colorspace_1.4-1   labeling_0.3       stringi_1.4.3
## [34] lazyeval_0.2.2     munsell_0.5.0      crayon_1.3.4
```
