## Supplementary document 3 for "Gene-set Enrichment with Regularized Regression"

### Gene-set enrichment with regularized regression: simulation studies with Gene Ontology

*July 08, 2019*

**Conclusions** **44**

**R session info** **44**

#### Background

In the manuscript *Gene-set Enrichment with Regularized Regression* (`gerr` in short), we propose using regularized regression to model the relationship between $Y$, a dichotomous dependent variable indicating membership of genes in a set of genes of interest (GOI hereafter), and $\Omega$, a matrix of dichotomous variables indicating membership of genes in gene-sets that are potentially overlapping or even identical with each other.

While `gerr` makes no explicit assumptions about the structure of the gene-set collection, some gene-sets are derived from tree-like structures. A prominent example is Gene Ontology (GO). Methods, for instance the `topGO` package in Bioconductor, has been developed to take advantage of the tree-like structure to decorrelate gene-sets. It is not known how a method `gerr`, which is agonistic about gene-set structure, performes in comparison with methods that take tree-like structures into account.

In this document, we perform simulation studies to demonstrate the sensitivity and specificity of the `gerr` method, using the software package of the same name that we published along with the manuscript and the elastic net implemented in the R software package `glmnet`. We compare results produced by `gerr` with the results of the `topGO` package.

Throughout the analysis, we will use the default parameters of `gerr`, using the elastic net of Gaussian-family linear regression, with $\alpha = 0.5$.

For methods implemented in `topGO` we will use Benjamini & Hochberg (1995) $p$-value correction with the FDR threshold of 0.05.

#### Gene-sets used for simulations

We wish to use real-world gene-sets that are commonly used by the community for the simulation study for `gerr`, because synthesized gene-sets may have distributions of sizes, defined by the number of unique genes and overlapping patterns, that depart from real-world gene-sets. In this simulation we use a randomly selected subset of Gene Ontology Biological process annotation.

When performing GO enrichment, we need to account for the fact that GO-term of a parent node is also associated with all the child nodes. In the code below we select a random subset of GO-categories and include all their parents into the list of nodes. We also ensure that all the parent terms are explicitly encoded in the G matrix for genes assoicated with their children.

```
# extract GO graph
go_graph <- makeGOGraph('bp')

# extract human gene sets
geneSets <- annFUN.org("BP", mapping = "org.Hs.eg.db", ID = "symbol") %>%
  keep(~length(.) > 10)
```

```
## Loading required package: org.Hs.eg.db

##
```

```
# choose ranom gene sets
set.seed(1)
randGeneSets <- sample(geneSets, 50)

# create a subset of GO graph including all the genes of interest
# note that the number of nodes in this graph will not only include
# randomGeneSets, but also all their parents
g_induced <- inducedGraph(go_graph, names(randGeneSets))

# extract levels
g_lev <- buildLevels(g_induced)

gs_induced <- list()

# propagate terms form children to parents
for (lev in getNoOfLevels(g_lev):3) {
  for (child in g_lev$level2nodes[[as.character(lev)]]) {
    for (parent in edges(g_induced)[[child]]) {
      gs_induced[[child]] <- c(
        gs_induced[[child]],
        geneSets[[child]]
      ) %>%
        unique()

      gs_induced[[parent]] <- c(
        gs_induced[[child]],
        gs_induced[[parent]],
        geneSets[[parent]]
      ) %>%
```

```
        unique()

    }
  }
}

stopifnot(g_lev$level2nodes[['1']]=='all')
# root will include all the genes
# we do not want to test for the root,
# as it does not make sence
# The root term GO:0008150 "Biological process"
root <- g_lev$level2nodes[['2']]
gs_induced[[root]] <- NULL

length(gs_induced)
```

```
## [1] 536
```

```
# this a list of gene sets including propagated terms
simGenesets <- gs_induced

# background, aka universe
bgGenes <- flatten_chr(simGenesets) %>%
  unique()

# create binary matrix for `gerr`
gsMatrix <- map(simGenesets, ~ bgGenes %in% .) %>%
  bind_cols() %>%
  as.matrix() %>%
  `*`(1) %>%
  `rownames<-`(bgGenes)

simLen <- map_int(simGenesets, length)
```

For the purpose of simulation, we randomly sample 50 gene-sets from all gene-sets and their parents. In total this makes 536 gene-sets (GO-terms).

#### Distribution of gene-set sizes

The histogram below shows the distribution of gene-set size in the sub-sampled set of gene-sets.

```
{
  hist(simLen, xlab="Number of unique genes",
       breaks=50, freq = FALSE,
       col="lightblue", main="Gene-set size distribution")
  lines(density(simLen, from=0), col="#004495", lwd=2)
}
```

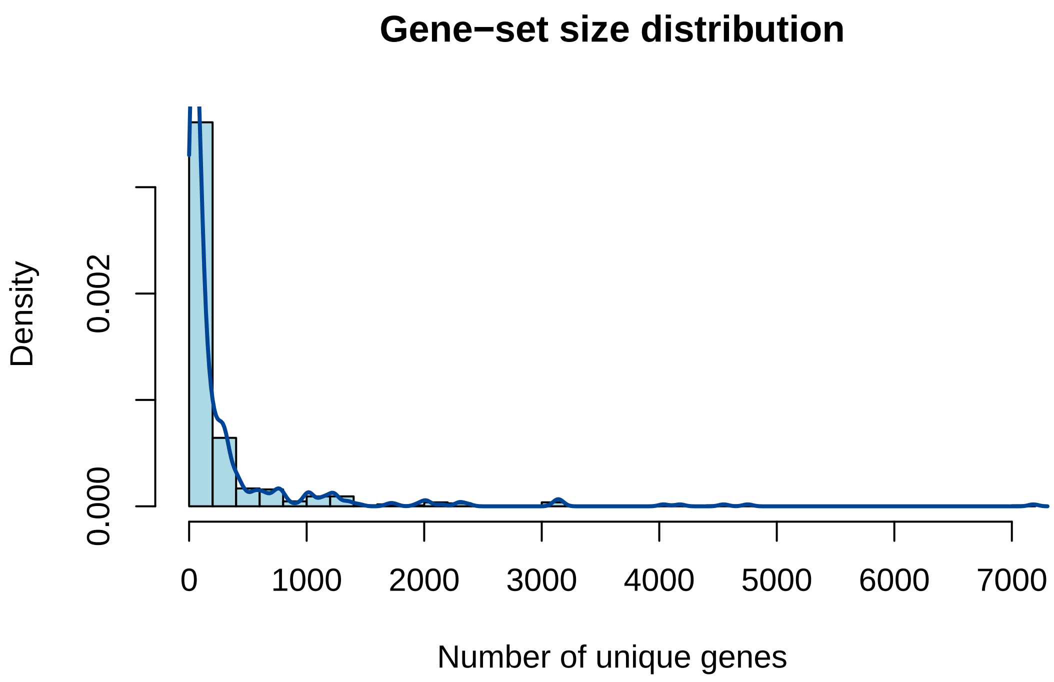

The median gene-set size is 83.5, with heavy tail on the right side.

##### Distribution of pairwise overlap coefficients between gene-sets

Next, we investigate the degree of redundancy among these gene-sets. We use the overlap coefficient, defined by $|A \cap B|/min(|A|, |B|)$ between two sets $A$ and $B$, to measure this.

When the gene-sets are represented as a dichotomous matrix, pairwise overlap coefficients can be calculated efficiently by using matrix operations. The functions below are extracted from the ribiosUtils package, which is currently being submitted to the CRAN repository.

```
uniqueNonNA <- function(x) {
  x <- x[!is.na(x)]
  res <- unique(x)
  return(res)
}
columnOverlapCoefficient <- function(x, y=NULL) {
  if(!is.matrix(x)) x <- as.matrix(x)
  if(is.null(y)) y <- x
  if(is.matrix(y)) y <- as.matrix(y)

  stopifnot(nrow(x)==nrow(y))
  storage.mode(x) <- storage.mode(y) <- "integer"

  tmatProd <- t(x) %*% y
  xCount <- apply(x, 2, function(xx) sum(xx!=0))
  yCount <- apply(y, 2, function(yy) sum(yy!=0))
  tmatPmin <- outer(xCount, yCount, pmin)
  res <- tmatProd/tmatPmin
  if(is.null(y)) {
    diag(res) <- 1L
  }
```

```
  dimnames(res) <- list(colnames(x), colnames(y))
  return(res)
}
listOverlapCoefficient <- function(x, y=NULL, checkUniqueNonNA=TRUE) {
  if(checkUniqueNonNA) {
    x <- lapply(x, uniqueNonNA)
    if(!is.null(y)) {
      y <- lapply(y, uniqueNonNA)
    }
  }

  if(is.null(y)) {
    elements <- unique(unlist(x))
    mat <- sapply(x, function(xx) as.integer(elements %in% xx))
    res <- columnOverlapCoefficient(mat)
  } else {
    elements <- unique(c(unlist(x), unlist(y)))
    mat1 <- sapply(x, function(xx) as.integer(elements %in% xx))
    mat2 <- sapply(y, function(xx) as.integer(elements %in% xx))
    res <- columnOverlapCoefficient(mat1, mat2)
  }
  return(res)
}
```

With these functions, we can calculate pairwise overlap coefficients between gene-sets.

```
simPairwiseOverlap <- columnOverlapCoefficient(gsMatrix) %>%
  as.dist() %>%
  broom::tidy(diagonal=FALSE, upper=TRUE) %>%
  rename(gsSim=item1, gsTest=item2, overlap=distance)

{
  hist(simPairwiseOverlap$overlap, xlab="Pairwise overlapping cofficient between gene-sets",
     breaks=50, freq = FALSE,
     col="orange",
     main="Overlapping coefficient distribution")
  lines(density(simPairwiseOverlap$overlap, from=0), col="red", lwd=2)
}
```

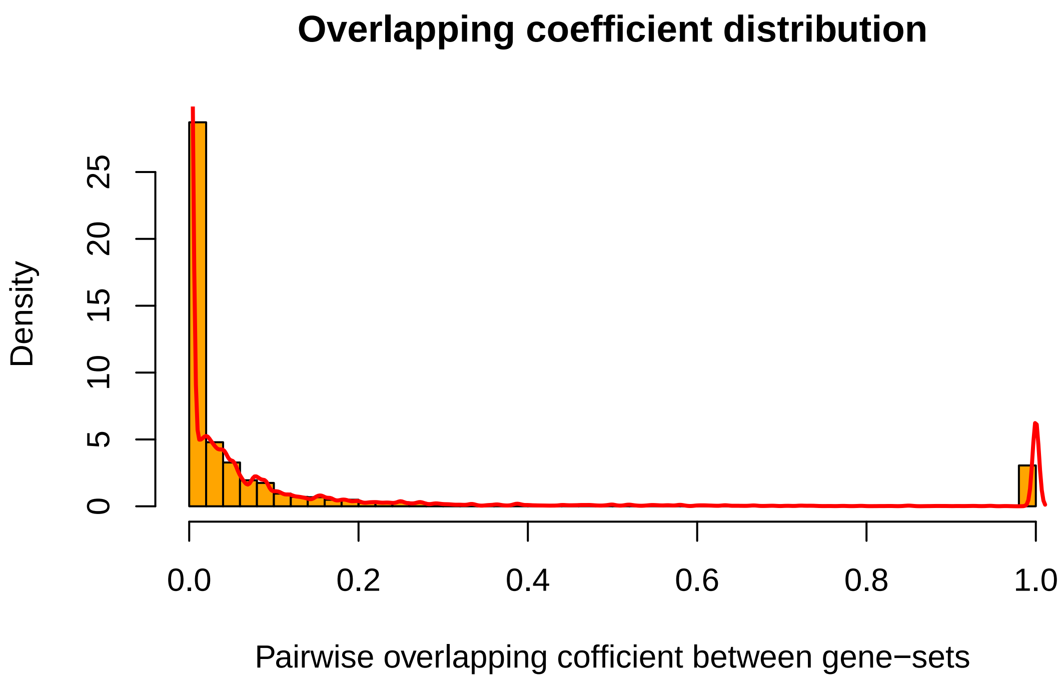

There is substantial overlap between genes. The peak on the right is most probably related to child-parent relationships. Note that because of high overlap it is unlikely that any methods will reliably distinguish with such GO-terms.

#### Model verification

We first verify that gene-set enrichment with regularized regression (`gerr`) performs as expected using the simplest simulation that is possible: we assign genes of one gene-set as GOI, and test whether we can recover the gene-set using `gerr`.

##### A small-scale verification with few cases

In the code chunk below, we verify that the model works in the sense that the gene-set that is used as source of GOI is correctly recovered, using randomly selected 10 gene-sets.

Since the analysis is computationally intensive, we use `clustermq` package and the `Q` function to parallelize computations on a cluster.

```
set.seed(2)
selInd <- sample(seq(along=simGenesets), 10)
selGsName <- names(simGenesets)[selInd]
selGs <- simGenesets[selInd]

selRes <- Q(function(i, goi, gsMatrix)
  regression_selected_pathways(
    gene_input=goi,
    gene_pathway_matrix=gsMatrix),
  i=seq_along(selGs),
  goi=selGs,
  gsMatrix=list(gsMatrix),
```

```
  n_jobs=10,
  export=list(regression_selected_pathways=regression_selected_pathways)
)
```

```
foundCounts <- sapply(seq(along=selInd),
                      function(i) length(selRes[[i]][[1]]))
isRecovered <- sapply(seq(along=selInd),
                      function(i) selGsName[i] %in% names(selRes[[i]][[1]]))

all(isRecovered)
```

```
## [1] TRUE
```

```
table(foundCounts)
```

```
## foundCounts
##  3  4  5  6  8 24
##  3  2  1  1  2  1
```

The `foundCounts` variable indicates the number of gene-sets whose coefficient is positive. In this particular example, we have multiple cases where more than one gene-set is selected. However in all cases the input gene-set is all recovered (`isRecovered` is all true).

#### The full-scale verification with all gene-sets

Below we run the simulation for all gene-sets.

```
estimate_gerr <- function(gs, gsMatrix) {
  tm <- system.time(
    res <- regression_selected_pathways(
      gene_input=gs,
      gene_pathway_matrix = gsMatrix,
      verbose=FALSE)
    )
    tibble(
      method='gerr',
      time=tm['elapsed'],
      res=list(tibble(
        gsTest=colnames(gsMatrix),
        detected=colnames(gsMatrix) %in% names(res$selected_pathways_names)
      )))
}

runGerr <- function(genesets, gsMatrix, index) {
  stopifnot(index %in% seq(along=genesets))
  selGsName <- names(genesets)[index]
  selGs <- genesets[index]
  imap_dfr(selGs, ~ estimate_gerr(., gsMatrix), .id='gsSim')
}
```

```
verifGerr <- Q(runGerr,
  index=seq_along(simGenesets),
  genesets=list(simGenesets),
  gsMatrix=list(gsMatrix),
  export=list(
```

```
    estimate_gerr=estimate_gerr
    ),
  pkgs=c('gerr', 'tidyverse'),
  n_jobs=100) %>%
  bind_rows()
```

We first confirm that each gene-set used as a GOI was successfully rediscovered by `gerr`.

```
verifGerr %>%
  unnest(res) %>%
  filter(gsTest==gsSim) %>%
  summarize(detected=mean(detected))
```

```
## # A tibble: 1 x 1
##   detected
##      <dbl>
## 1        1
```

Next we query the frequency of cases where `gerr` returned more than one gene-set.

```
verifGerr %>%
  unnest(res) %>%
  group_by(gsSim) %>%
  summarize(n_fp=sum(detected & gsSim!=gsTest)) %>%
  count(n_fp)
```

```
## # A tibble: 19 x 2
##     n_fp     n
##    <int> <int>
##  1     0     7
##  2     1    45
##  3     2    73
##  4     3    96
##  5     4    71
##  6     5    59
##  7     6    51
##  8     7    47
##  9     8    15
## 10     9    25
## 11    10     5
## 12    11     2
## 13    12     1
## 14    13     8
## 15    14     6
## 16    16    19
## 17    18     2
## 18    23     2
## 19    26     2
```

Now we implement the FET+FDR strategy.

```
fetFdr <- function(goi, bgGenes, goi_name=NULL) {
  goi <- uniqueNonNA(goi)
  bgGenes <- uniqueNonNA(bgGenes)
  bgDiffGoi <- setdiff(bgGenes, goi)
  tm <- system.time(
    fetRes <- imap_dfr(simGenesets, function(gs, gs_name) {
```

```
      selHits <- length(intersect(gs, goi))
      nonselHits <- length(goi) - selHits
      selNonhits <- length(gs) - selHits
      nonselNonhits <- length(bgDiffGoi) - selNonhits
      mat <- matrix(c(selHits, nonselHits, selNonhits, nonselNonhits),2,2)
      pval <- stats::fisher.test(mat, alternative = "greater")$p.value
      list(gsTest=gs_name,
           p_value=pval)
  }) %>%
    mutate(p_adj=p.adjust(p_value, "BH"),
           detected=p_adj<fdr_thr)
  )
  tibble(
    gsSim=goi_name,
    time=tm['elapsed'],
    method='FET',
    res=list(fetRes)
  )
}
```

Next we run the verification step using FET+FDR.

```
fisherVerif <- Q(fetFdr,
                 goi=simGenesets,
                 goi_name=names(simGenesets),
                 bgGenes=list(bgGenes),
                 pkgs=c('tidyverse'),
                 export=list(uniqueNonNA=uniqueNonNA,
                             simGenesets=simGenesets,
                             fdr_thr=fdr_thr),
                 n_jobs=10
) %>%
    bind_rows()
```

Next, we perform the same analysis with topGO. Note that topGO implements several strategies to reduce the graph redundancy. We try all the methods implemented. The `classic` method is exactly identical to FET+FDR after propagation of terms along the GO-graph.

```
# we create this object and reuse it later
tgData <- new(
  'topGOdata',
  description = 'simluation',
  ontology = 'BP',
  nodeSize=1,
  allGenes = bgGenes %in% simGenesets[[1]] %>%
  as.integer %>%
  set_names(bgGenes) %>%
  as.factor(),
  annot = annFUN.GO2genes,
  GO2genes=simGenesets
)

topgo_methods <- c('classic', 'elim', 'weight', 'weight01', 'lea', 'parentchild')
```

```
estimate_topgo <- function(gs, tgData, method) {
  data <- updateGenes(
    tgData,
    bgGenes %in% gs %>%
      as.integer %>%
      set_names(bgGenes) %>%
      factor(levels=c(0, 1)))
  tm <- system.time(
    test_res <- runTest(data,
                        algorithm=method,
                        statistic="fisher")
  )
  p_value <- score(test_res)
  tibble(
    method=method,
    time=tm['elapsed'],
    res=list(tibble(
      gsTest=names(p_value),
      p_value=p_value
      ) %>%
        filter(gsTest != root) %>%
        mutate(p_adj=p.adjust(p_value, method='BH'),
       detected=p_adj < fdr_thr)
    )
  )
}
```

```
topgo_results <- map_dfr(topgo_methods, function(method)
  Q(estimate_topgo,
    gs=simGenesets,
    tgData=list(tgData),
    method=method,
    pkgs=c('tidyverse', 'topGO'),
    export=list(bgGenes=bgGenes,
                root=root,
                fdr_thr=fdr_thr),
    n_jobs=150
  ) %>%
    set_names(names(simGenesets)) %>%
    bind_rows(.id='gsSim')
)
```

Next, we summarize the running time of different methods.

```
all_verif <- topgo_results %>%
  bind_rows(verifGerr) %>%
  bind_rows(fisherVerif)

all_verif %>%
  group_by(method) %>%
  summarize(time=sum(time))
```

```
## # A tibble: 8 x 2
##   method       time
##   <chr>       <dbl>
```

```
## 1 classic      472.
## 2 elim         569.
## 3 FET          105.
## 4 gerr        2698.
## 5 lea          549.
## 6 parentchild 1379.
## 7 weight      1845.
## 8 weight01    1024.
```

The fastest method is FET+FDR. Probably because we preprocessed the GO-graph once prior to analysis. `gerr` is the slowest method in our comparison.

Below we summarize false-positive rate (FPR) and true-positive rate (TPR).

```
all_verif %>%
  unnest(res) %>%
  group_by(method) %>%
  summarize(
    fpr=sum(detected & (gsSim!=gsTest))/sum(gsSim!=gsTest),
    tpr=sum(detected & (gsSim==gsTest))/sum(gsSim==gsTest)
  )
```

```
## # A tibble: 8 x 3
##   method          fpr   tpr
##   <chr>         <dbl> <dbl>
## 1 classic     0.134   1
## 2 elim        0.0171  0.424
## 3 FET         0.134   1
## 4 gerr        0.00958 1
## 5 lea         0.0272  0.541
## 6 parentchild 0.0809  0.920
## 7 weight      0.0117  0.494
## 8 weight01    0.0168  0.422
```

Notice that `gerr` has the lowest FPR, while having TPR of 1. This is the best performance among methods.

We compare results of `topGO` in the `classic` mode with FET+FDR. The results should be identical, as this is the same method.

```
all_verif %>%
  filter(method %in% c('classic', 'FET')) %>%
  unnest(res) %>%
  select(gsSim, gsTest, method, p_adj) %>%
  spread(method, p_adj) %>%
  mutate(diff=abs(classic-FET)) %>%
  pull(diff) %>%
  summary()
```

```
##    Min. 1st Qu.  Median    Mean 3rd Qu.    Max.
##       0       0       0       0       0       0
```

Now we first compare the number of false-positive hits in all the methods.

```
all_verif %>%
  unnest(res) %>%
  group_by(method, gsSim) %>%
  summarize(fp=sum(gsSim!=gsTest & detected)) %>%
  ggplot(aes(fp, stat(density), col=method)) +
```

```
  geom_freqpoly(size=1) +
  xlab("False-positive hits") +
  ylab("Gene-set count") +
  theme(axis.text.x=element_text(angle = 0)) +
  scale_color_brewer(palette = "Set1") +
  #facet_wrap(~method, ncol=1) +
  theme(strip.background = element_blank(),
        strip.text.x = element_blank())
```

```
## `stat_bin()` using `bins = 30`. Pick better value with `binwidth`.
```

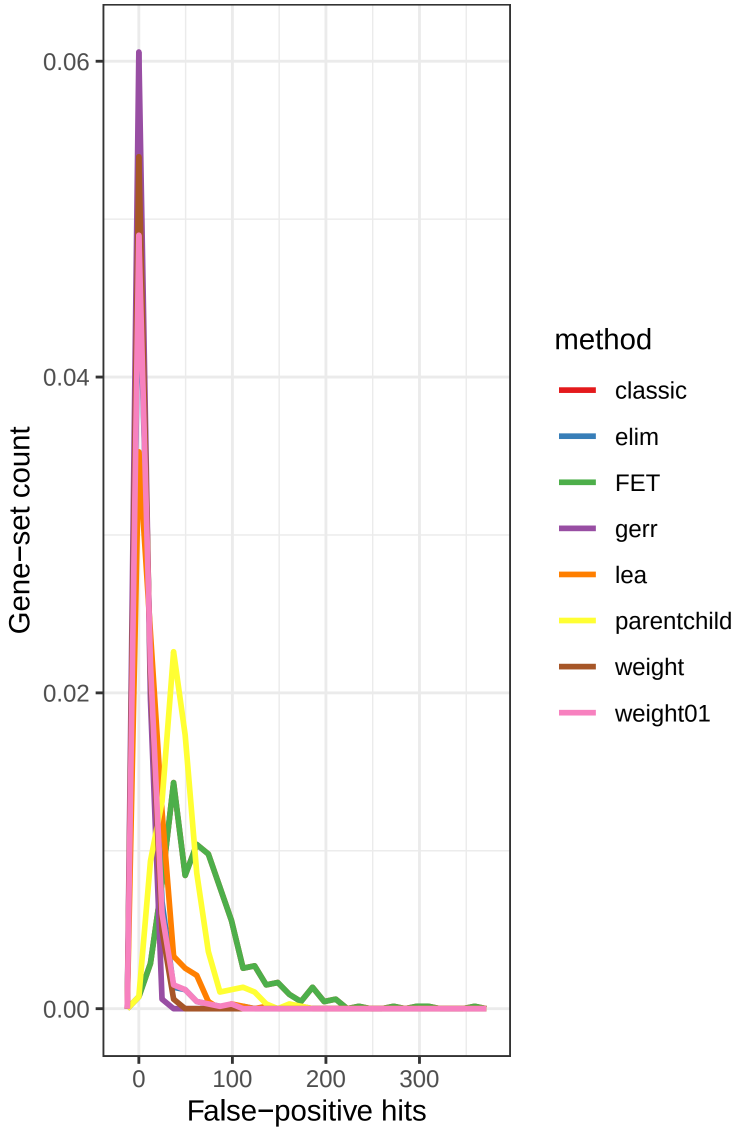

We see that number of false positives is high in the following methods: `classic` (aka `FET`) and `parentchild`. We remove those and focus on the rest.

```
all_verif %>%
  unnest(res) %>%
  group_by(method, gsSim) %>%
  filter(method %in% c('elim', 'gerr', 'lea', 'weight', 'weight01')) %>%
```

```
  summarize(fp=sum(gsSim!=gsTest & detected)) %>%
  ggplot(aes(fp, stat(density), col=method)) +
  geom_freqpoly(binwidth=2, size=0.8) +
  xlab("False-positive hits") +
  ylab("Gene-set count") +
  theme(axis.text.x=element_text(angle = 0)) +
  scale_color_brewer(palette = "Set1")
```

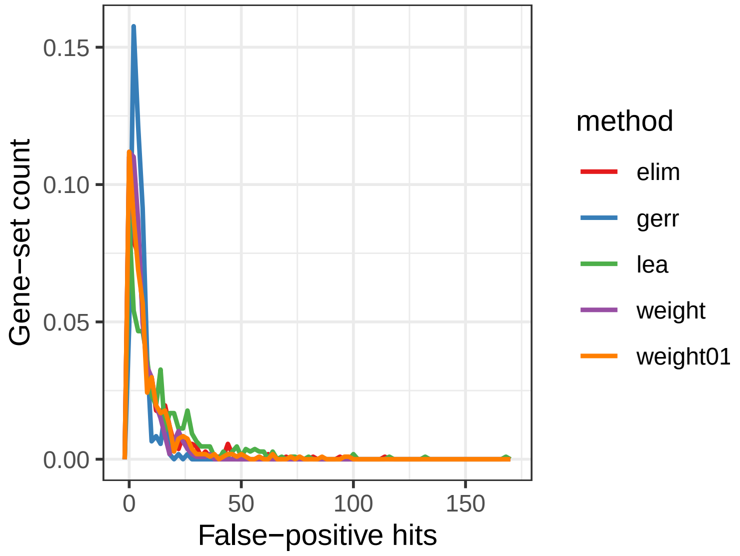

We can also compute difference in frequencies.

```
max_fp <- all_verif %>%
  unnest(res) %>%
  group_by(method, gsSim) %>%
  summarize(fp=sum(gsSim!=gsTest & detected)) %>%
  ungroup() %>%
  summarize(max_fp=max(fp)) %>%
  pull(max_fp)

dens_df <- all_verif %>%
  unnest(res) %>%
  group_by(method, gsSim) %>%
  filter(method %in% c('elim', 'gerr', 'lea', 'weight', 'weight01')) %>%
  summarize(fp=sum(gsSim!=gsTest & detected)) %>%
  group_by(method) %>%
  summarize(hist=list(hist(fp, plot=FALSE, breaks=seq(0, max_fp+1, 5)))) %>%
  rowwise() %>%
  mutate(hist=list(tibble(mids=hist$mids, density=hist$density))) %>%
  unnest(hist)

filter(dens_df, method=='gerr') %>%
  select(-method) %>%
  rename(density_gerr=density) %>%
  left_join(filter(dens_df, method!='gerr')) %>%
  mutate(method=str_c('frequency(gerr) - frequency(', method, ')')) %>%
  mutate(diff_density=density_gerr-density) %>%
  ggplot(aes(mids, diff_density, col=method)) +
```

```r
  geom_line(size=0.5) +
  labs(x='Difference in frequency', y='Number of false positives') +
  scale_color_brewer(palette = "Set1") +
  theme(legend.position="bottom",
        legend.direction="vertical")
```

```
## Joining, by = "mids"
```

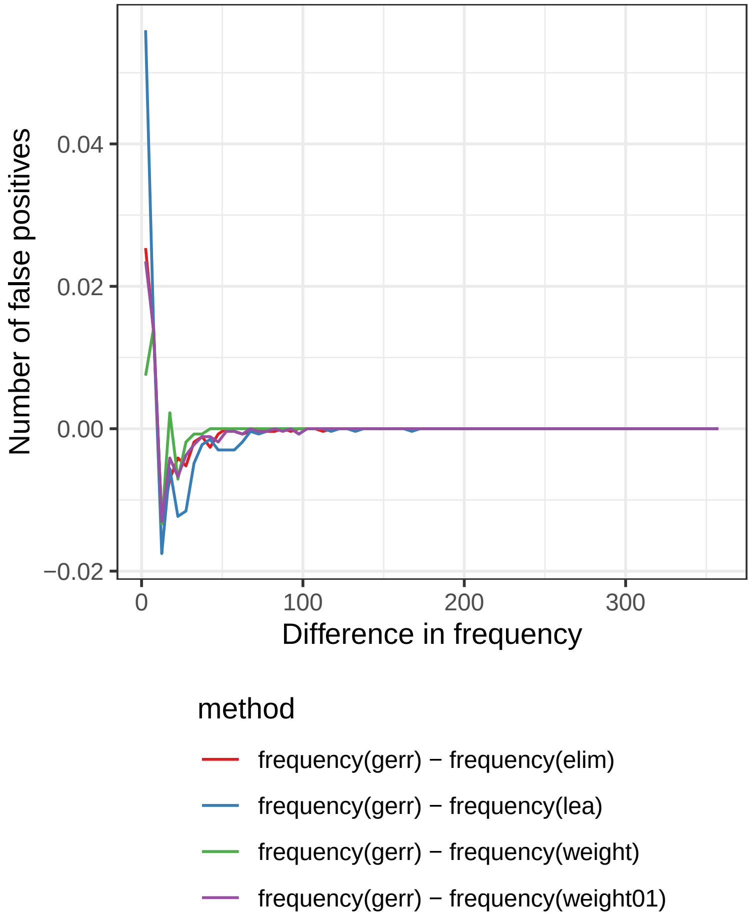

The `gerr` has the largest peak on the left, confirming low false positive rate observed before.

We believe that the false-positive hits of `gerr` are likely caused by redundancy in gene-sets. To show that, we illustrate below the mean overlap coefficient between false-positive and true-positive hits returned by `gerr`. As a comparison, we show the value also for hits returned by FET+FDR and for all gene-sets.

```r
all_verif %>%
  unnest(res) %>%
  filter(method=='classic') %>%
  mutate(detected=TRUE, method='all') %>%
  bind_rows(all_verif %>% unnest(res)) %>%
  left_join(simPairwiseOverlap) %>%
  filter((gsSim != gsTest) & detected) %>%
  ggplot(aes(overlap, col=method)) +
  geom_freqpoly(size=1) +
  scale_color_brewer(palette = "Set1") +
  facet_wrap(~method, ncol=1, scales='free_y') +
  theme(strip.background = element_blank(),
```

```
        strip.text.x = element_blank())
```

```
## Joining, by = c("gsSim", "gsTest")

## Warning: Column `gsSim` joining character vector and factor, coercing into
## character vector

## Warning: Column `gsTest` joining character vector and factor, coercing into
## character vector

## `stat_bin()` using `bins = 30`. Pick better value with `binwidth`.
```

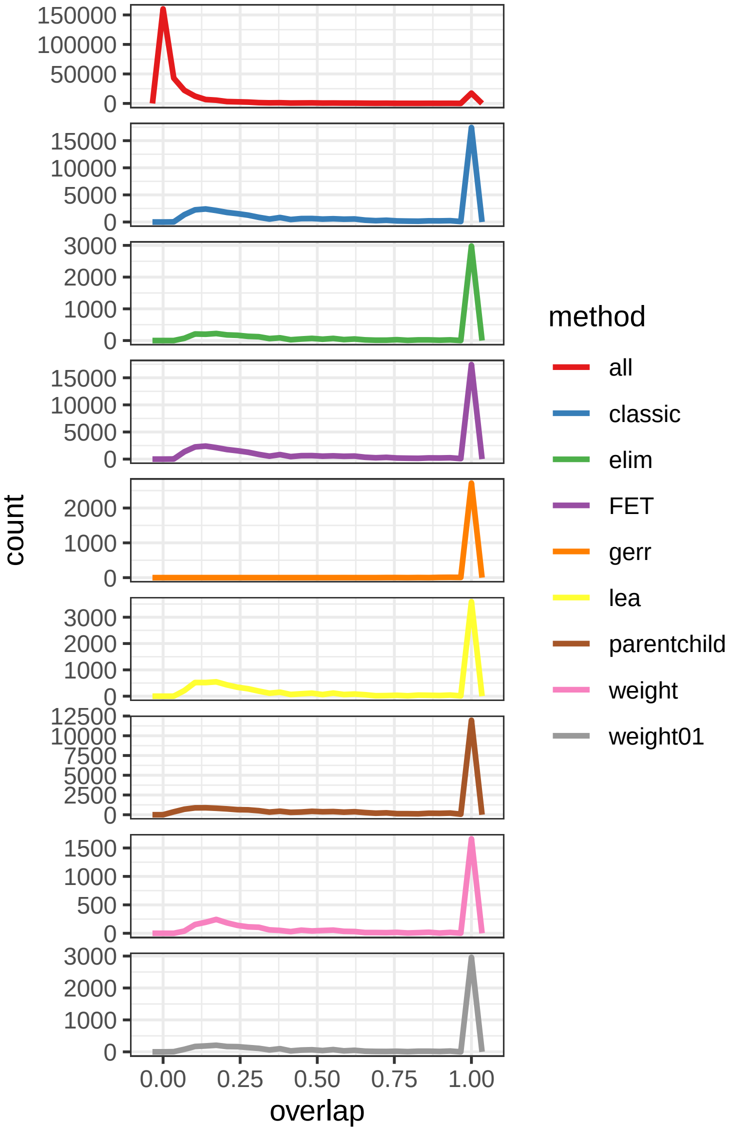

In short, with a very simple verification procedure, we verify that by using `gerr`, all gene-sets that were assigned as genes of interest were successfully recovered. In most cases, the only gene-set selected by `gerr` was the true positive hit. In other cases, more than one gene-set was selected by `gerr`. The co-selected gene-sets are highly redundant with the true positive hit.

#### Simulations based on a probabilistic model

Next we test the performance of `gerr` using a probabilistic framework. Specifically, we assume a generative model of GOI in the following form:

$$p_{g \in G} = \sum_{\omega} p_{g|\omega} p_{\omega} + p_{g_n}$$

.

The model specifies that the probability that a gene $g$ is a member of the set $G$ is modeled by the probability of the gene-set $\omega$ contributes to GOI, expressed as $p_{\omega}$, multiplied by the probability that $g$ is selected to contributed to GOI given that $\omega$ contributes to GOI, summed over all gene-sets, and then adding the term $p_{g_n}$ that models the probability that the gene $g$ contributes to GOI independent of its associations with gene-sets. The two parts on the right side of the equation can be observed as a gene-set dependent and a noise term (therefore the subscript $n$) respectively. Apparently, $p_{g \in G}$ should be located between $[0, 1]$.

In the following simulations, we assume that $p_{\omega}$ follows a binomial distribution that is specified by the parameter $p_{gs}$. For simplicity, we model $p_{g|\omega}$ as a binomial distribution specified by the parameter $p_g$ and assume that the same parameter applies to all selected gene-sets. Furthermore, we assume that $p_{g_n}$ is modeled by a small number $p_n$ that varies in a given range, e.g. from $10^{-4}$ to $10^{-1}$, and the value applies to all genes.

The simulation procedure can be described in following steps:

1. Randomly sample $k$ gene-sets from $\Omega$, the set of gene-sets, with the probability of $p_{gs}$.
2. Within each $k$ gene-sets, randomly select $m$ genes each with the probability of $p_g$.
3. Randomly sample genes from the background $B$ by a uniform distribution with the parameter $p_n$.
4. Merge genes selected in step (2) and (3) into a set of GOI.
5. Perform the `gerr` and `topGO` analysis with $G$ and $\Omega$ as input
6. Assess the specificity and sensitivity of all the methods.

The codes below implement the logic.

```
probSim <- function(p_gs=0.01, p_g=0.5, p_n=1E-3, seed=1L) {
  set.seed(seed)
  ## step 1
  gsSel <- rbinom(n=length(simGenesets), size=1, prob=p_gs) == 1
  gsSelNames <- names(simGenesets)[gsSel]
  ## step 2
  gsGenes <- lapply(simGenesets[gsSel], function(genes) {
    selGenes <- rbinom(n=length(genes), size=1, prob=p_g)
    res <- genes[selGenes==1]
    return(res)
  })
  ## step 3
  isNoiseGenes <- rbinom(n=length(bgGenes), size=1, prob=p_n) == 1
  noiseGenes <- bgGenes[isNoiseGenes]
  ## step 4
  tibble(
    goi=list(unique(c(unlist(gsGenes), noiseGenes))),
    gs=list(gsSelNames)
  )
}

estimate <- function(goi) {
  ## step 5
  map_dfr(topgo_methods,
          ~estimate_topgo(goi, tgData, .x)) %>%
```

```
    bind_rows(
      estimate_gerr(goi, gsMatrix)
    )
}

compute_performance <- function(gs, res) {
  ## step 6
  res <- res %>%
    mutate(sim=gsTest %in% gs)

  perf <- res %>%
    summarize(
      tpr=sum(sim&detected)/sum(sim),
      fpr=sum(detected&!sim)/sum(!sim),
      ppv=sum(sim&detected)/sum(detected),
      f1=2*ppv*tpr/(ppv+tpr)
  )

  fp <- res %>%
    filter(!sim & detected) %>%
    pull(gsTest)

  overlap <- simPairwiseOverlap %>%
    filter(gsSim %in% gs & gsTest %in% fp) %>%
    summarize(min_overlap=min(overlap), median_overlap=median(overlap))

  perf$min_overlap <- overlap$min_overlap
  perf$median_overlap <- overlap$median_overlap

  perf
}

estimate_and_performance <- function(df) {
  df %>%
    rowwise() %>%
    mutate(est=list(estimate(goi))) %>%
    unnest(est, .drop=FALSE) %>%
    rowwise() %>%
    mutate(perf=list(compute_performance(
      gs,
      res))) %>%
  unnest(perf)
}
```

#### A small-scale simulation with the probabilistic model

For instance, below we example the results of the particular simulation setting.

```
testSim <- probSim(p_gs=0.05, p_g=0.5, p_n=1E-3) %>%
  estimate_and_performance()
```

```r
testSim %>%
  select(method, time, tpr, fpr, ppv, f1, median_overlap, min_overlap)
```

```
## # A tibble: 7 x 8
##   method       time   tpr    fpr    ppv    f1 median_overlap min_overlap
##   <chr>       <dbl> <dbl>  <dbl>  <dbl> <dbl>          <dbl>       <dbl>
## 1 classic      1.25 1     0.685  0.0745 0.139        0                 0
## 2 elim         1.85 0.429 0.157  0.130  0.2          0                 0
## 3 weight       5.20 0.321 0.0531 0.25   0.281        0.0130            0
## 4 weight01     3.22 0.429 0.152  0.135  0.205        0                 0
## 5 lea          1.66 0.571 0.236  0.118  0.195        0.00990           0
## 6 parentchild  3.41 0.857 0.344  0.121  0.211        0.0150            0
## 7 gerr        18.8  0.821 0.177  0.204  0.326        0                 0
```

#### A full scale simulation with the probabilistic model and results interpretation

Below we perform the full-scale simulation with the probabilistic model, using a variety of combinations of the parameters $p_{gs}$, $p_g$, and $p_n$. For each parameter set, five independent simulations are run.

```r
p_gs_cand <- c(0.002, 0.005, 0.01,
               seq(0.02, 0.1, by=0.01))
p_g_cand <- c(0.05, seq(0.1, 1, by=0.1))
p_n_cand <- c(0, 1E-4, 1E-3, 1E-2, 5E-2, 1E-1)

probSimParamsOneRep <- expand.grid(p_gs=p_gs_cand,
  p_g=p_g_cand,
  p_n=p_n_cand) %>%
  ## five replicates per condition
  slice(rep(1:n(), each = 5)) %>%
  mutate(seed=1:n())
```

Due to the stochastic nature of sampling, in some runs no gene-set is selected at all to contribute to GOI, especially when $p_{gs}$ is set small. These cases are excluded from the analysis below, because they will distort the results. Including them however do not change the conclusions.

Below we run the full simulation.

```r
full_sim <- probSimParamsOneRep %>%
  rowwise() %>%
  mutate(
    sim=list(probSim(p_gs=p_gs, p_g=p_g, p_n=p_n, seed=seed))
    ) %>%
  unnest(sim) %>%
  rowwise() %>%
  filter(length(goi) > 3)
```

Next we estimate gene sets. Warning: this is *very* slow.

```r
# this is quite slow
full_sim_res <- Q(
  function(i) estimate_and_performance(full_sim[i,]),
  i=1:nrow(full_sim),
  pkgs=c('tidyverse', 'gerr', 'topGO'),
  export=list(
    bgGenes=bgGenes,
```

```
    tgData=tgData,
    gsMatrix=gsMatrix,
    full_sim=full_sim,
    simPairwiseOverlap=simPairwiseOverlap,
    topgo_methods=topgo_methods,
    estimate_and_performance=estimate_and_performance,
    compute_performance=compute_performance,
    estimate=estimate,
    estimate_gerr=estimate_gerr,
    estimate_topgo=estimate_topgo,
    root=root,
    fdr_thr=fdr_thr
  ),
  n_jobs=500
) %>%
  bind_rows()
```

```
## Warning in qsys$cleanup(quiet = !verbose): 58/277 workers did not shut down
## properly
```

Summary of the simulations is shown below.

```
full_sim_res %>%
  select(method, time, tpr, fpr, ppv, f1, median_overlap) %>%
  group_by(method) %>%
  summarize_all(median, na.rm=TRUE)
```

```
## # A tibble: 7 x 7
##   method       time   tpr    fpr    ppv    f1 median_overlap
##   <chr>       <dbl> <dbl>  <dbl>  <dbl> <dbl>          <dbl>
## 1 classic      1.32 1      0.608  0.0736 0.136        0.0115
## 2 elim         1.97 0.324 0.106  0.124  0.179        0.00698
## 3 gerr        15.2  0.72  0.143  0.182  0.284        0
## 4 lea          1.83 0.489 0.182  0.117  0.188        0.0154
## 5 parentchild  3.76 0.667 0.289  0.0987 0.168        0.0222
## 6 weight       6.34 0.25  0.0459 0.211  0.229        0.0179
## 7 weight01     3.67 0.289 0.0955 0.123  0.172        0.00714
```

The median values of the five simulation runs are reported for each measure (true-positive rate, false-positive rate, *etc.*).

```
full_sim_res_agg <- full_sim_res %>%
  filter(!is.na(f1)) %>%
  select(p_gs, p_g, p_n, method, tpr, fpr, ppv, f1) %>%
  group_by(p_gs, p_g, p_n, method) %>%
  summarise_all(median) %>%
  mutate(m_fpr=1-fpr)

write_tsv(full_sim_res_agg,
          'full_sim_results_aggregated.tsv.gz')
```

#### Interpretation of the simulation results

We investigate the simulation results by visualizing true positive rate, false positive rate, and $F_1$ scores of the `gerr` and classic (aka FET+FDR) procedure.

##### Sensitivity, or true positive rate (TPR)

The plot below visualizes how sensitivity, or true positive rate (TPR) varies by the probability that each gene-set contributes to GOI ($p_{gs}$) and the probability of genes in each gene-set contribute to GOI ($p_g$), conditional on the noise probability $p_n$. Five independent simulations were performed for each parameter set, and the median value is used for visualization.

```
lowCol <- "#004495"
midCol <- "#CCCCCC"
highCol <- "#AA3555"
theme_update(axis.text.x = element_text(angle=45, hjust=1),
        axis.text = element_text(size=11),
        axis.title = element_text(size=14),
        strip.text = element_text(size=11),
        legend.text = element_text(size=11))

plot_stat <- function(res, m, stat, label, name) res %>%
  filter(method==m) %>%
  ggplot(aes(x=factor(p_gs), y=factor(p_g), fill=!!sym(stat))) +
  facet_wrap(~p_n, labeller = label_both) +
  geom_tile() +
  scale_fill_gradient2(low=lowCol, mid=midCol, high=highCol,
                       midpoint=0.5,
                       limits=c(0,1),
                       oob=scales::squish,
                       name=label) +
  ggtitle(str_c(name, " of ", m))+
  xlab(expression(p[gs])) + ylab(expression(p[g]))

plot_stat(full_sim_res_agg, 'gerr', 'tpr', 'TPR', 'True positive rate')
```

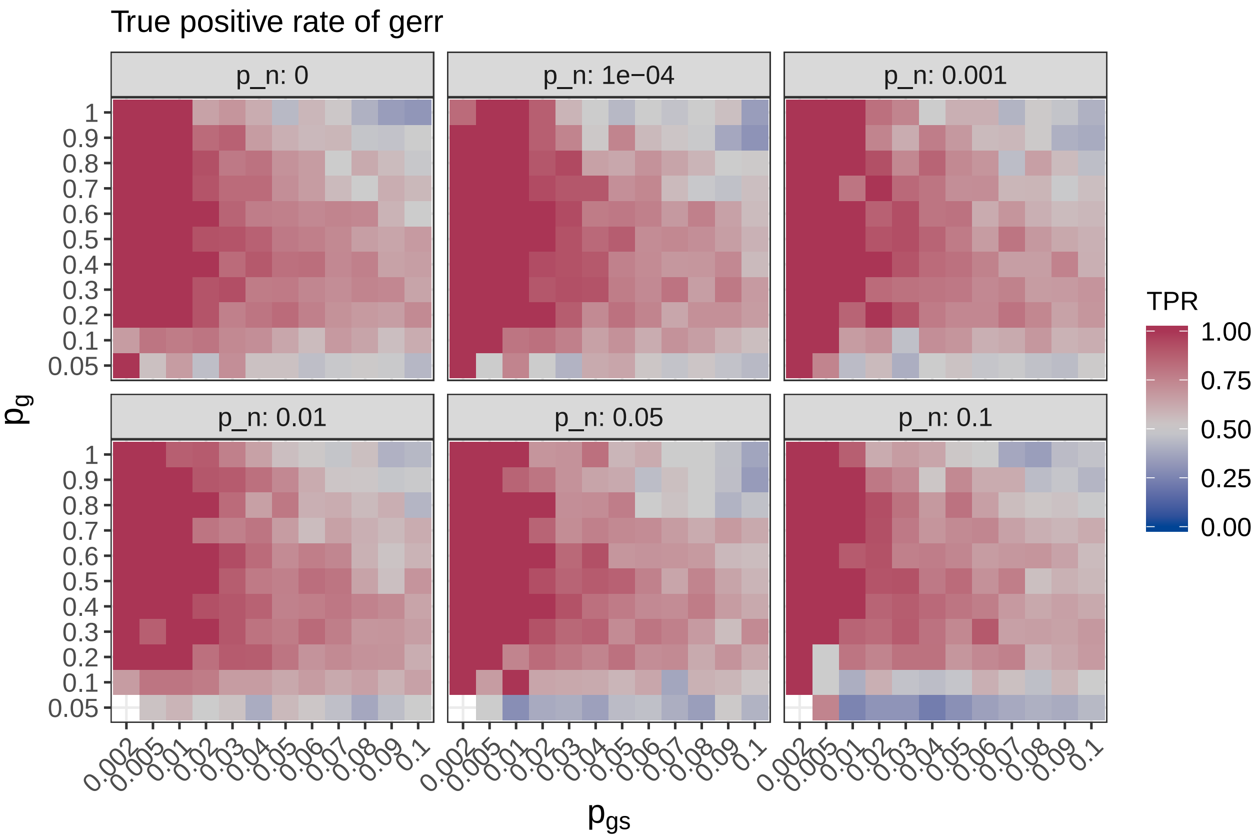

It seems that **gerr** in general has high sensitivity, even when the noise probability is as high as 0.1 (namely each gene has the probability of 0.1 to be selected as a gene of interest, independent whether it is associated with any gene-set or not). The sensitivity is only low when very few genes in the gene-set contribute to GOI (say less than 10%), which makes sense intuitively.

Some cells are missing because in five runs of simulation, the sampling procedure did not pick any gene-set to contribute to contribute to GOI.

The plot below visualizes the pattern of TPR for other procedures.

```
walk(topgo_methods, ~print(plot_stat(
  full_sim_res_agg,
  .,
  'tpr',
  'TPR',
  'True positive rate')))
```

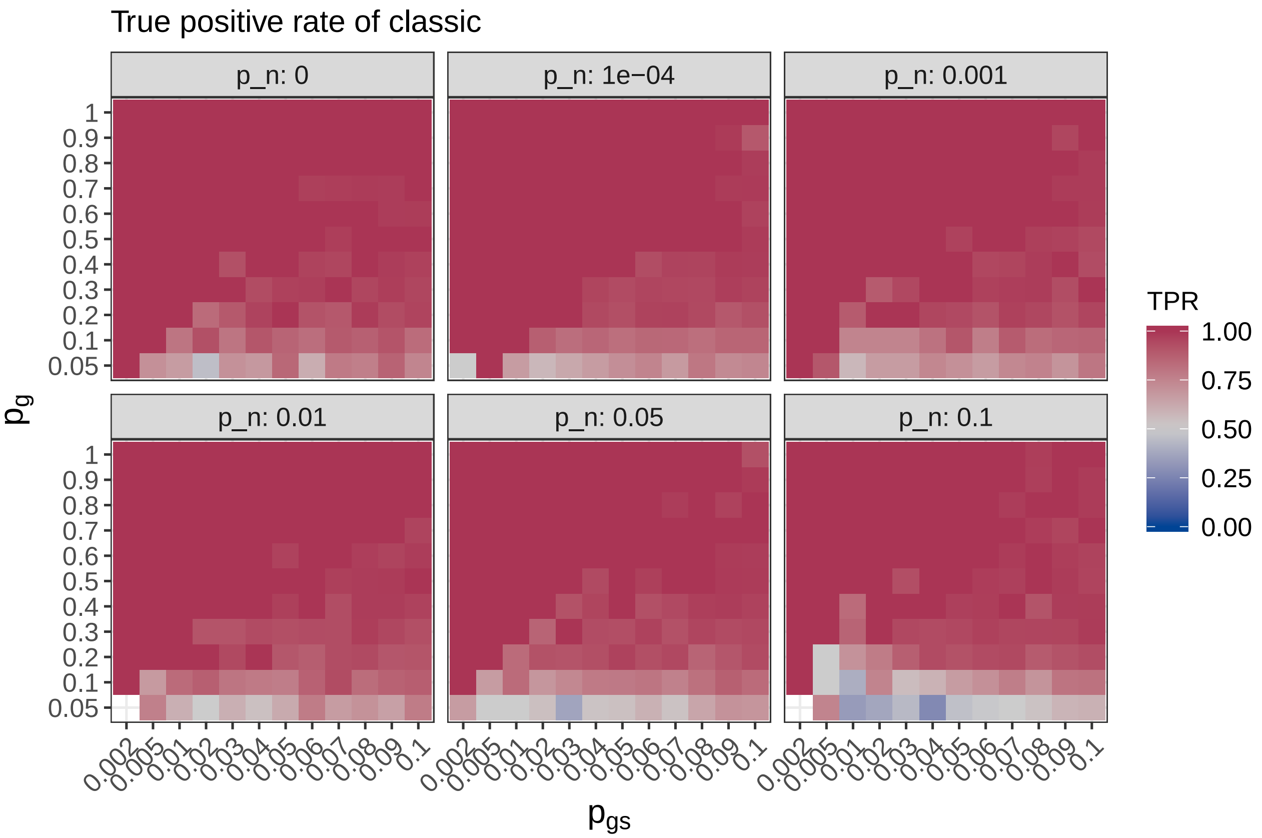
True positive rate of classic
p_n: 0
p_n: 1e−04
p_n: 0.001
p_n: 0.01
p_n: 0.05
p_n: 0.1
TPR
1.00
0.75
0.50
0.25
0.00
1
0.9
0.8
0.7
0.6
0.5
0.4
0.3
0.2
0.1
0.05
0.002
0.005
0.01
0.02
0.03
0.04
0.05
0.06
0.07
0.08
0.09
0.1
$p_g$
$p_{gs}$

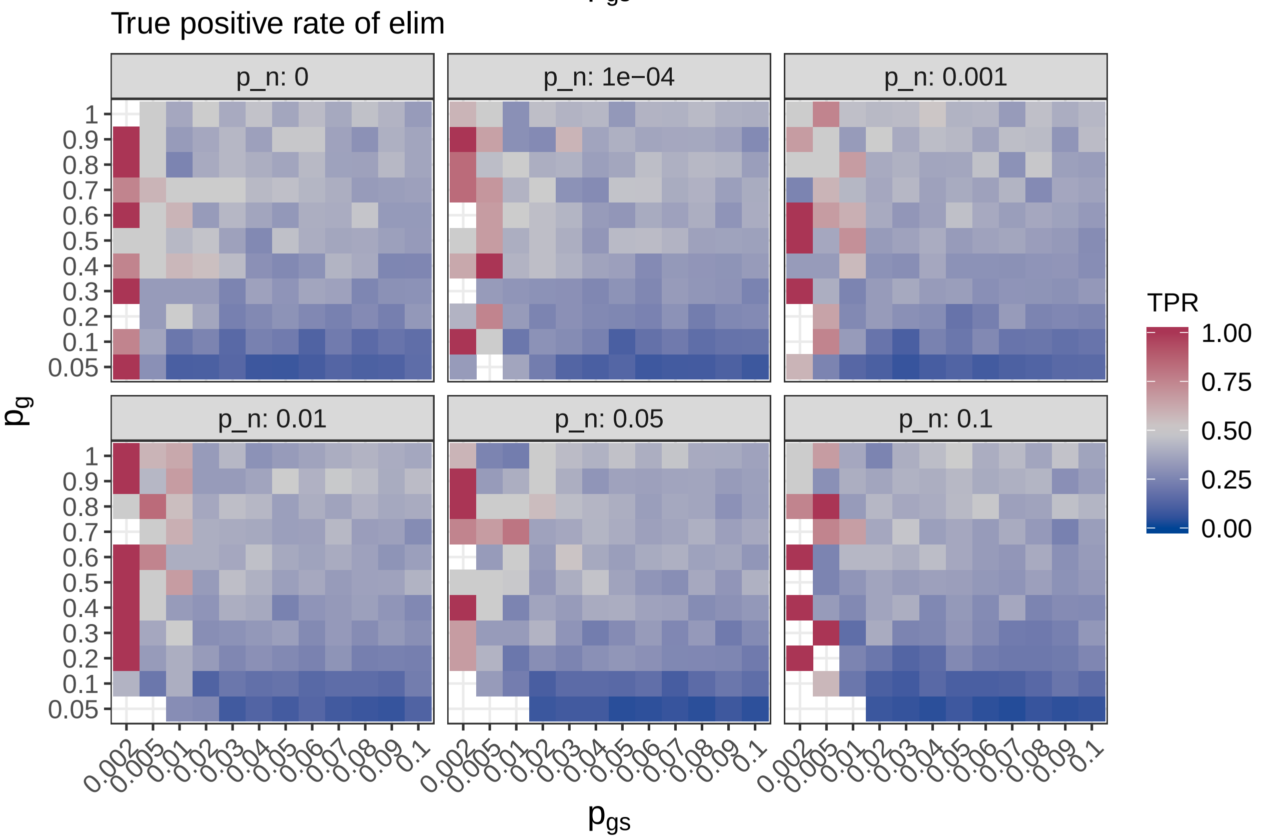
True positive rate of elim
p_n: 0
p_n: 1e−04
p_n: 0.001
p_n: 0.01
p_n: 0.05
p_n: 0.1
TPR
1.00
0.75
0.50
0.25
0.00
1
0.9
0.8
0.7
0.6
0.5
0.4
0.3
0.2
0.1
0.05
0.002
0.005
0.01
0.02
0.03
0.04
0.05
0.06
0.07
0.08
0.09
0.1
$p_g$
$p_{gs}$

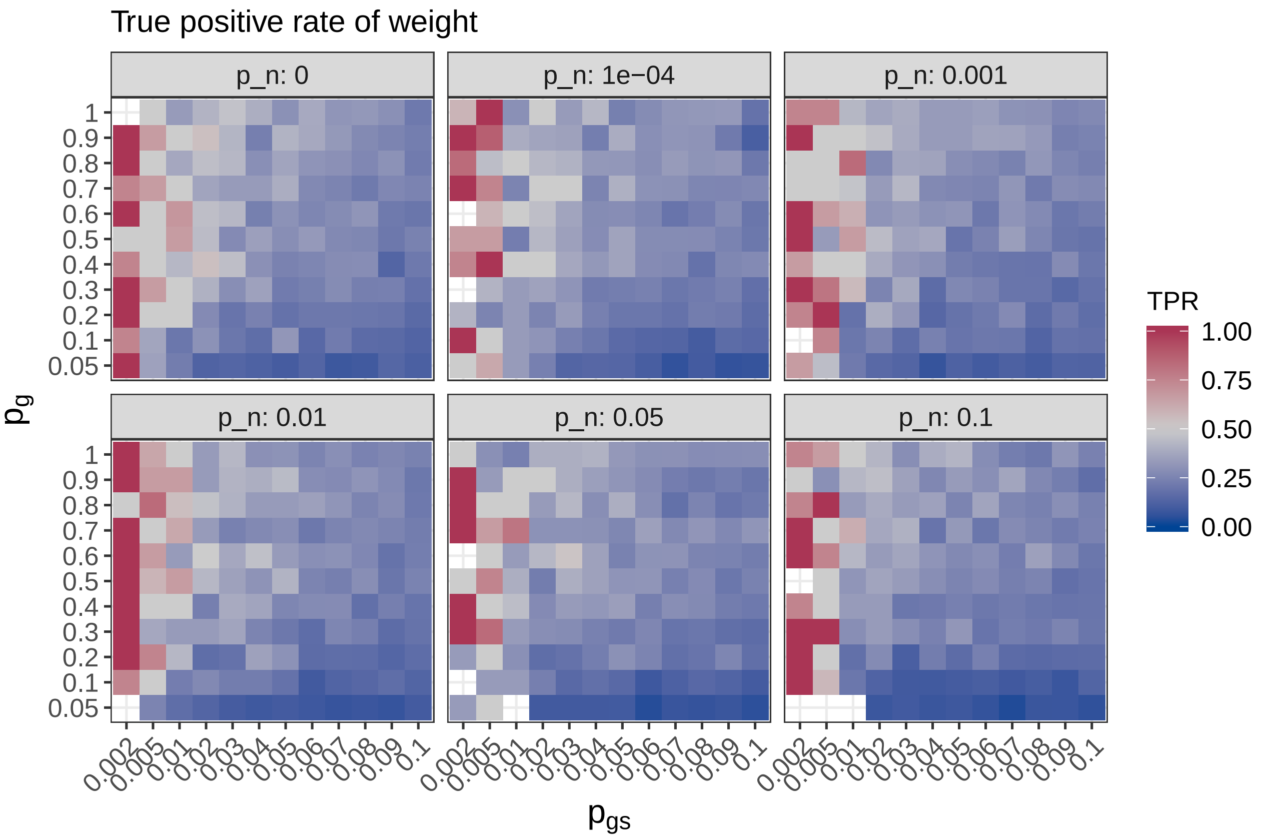
True positive rate of weight
p_n: 0
p_n: 1e−04
p_n: 0.001
p_n: 0.01
p_n: 0.05
p_n: 0.1
1
0.9
0.8
0.7
0.6
0.5
0.4
0.3
0.2
0.1
0.05
0.002
0.005
0.01
0.02
0.03
0.04
0.05
0.06
0.07
0.08
0.09
0.1
pg
pgs
TPR
1.00
0.75
0.50
0.25
0.00

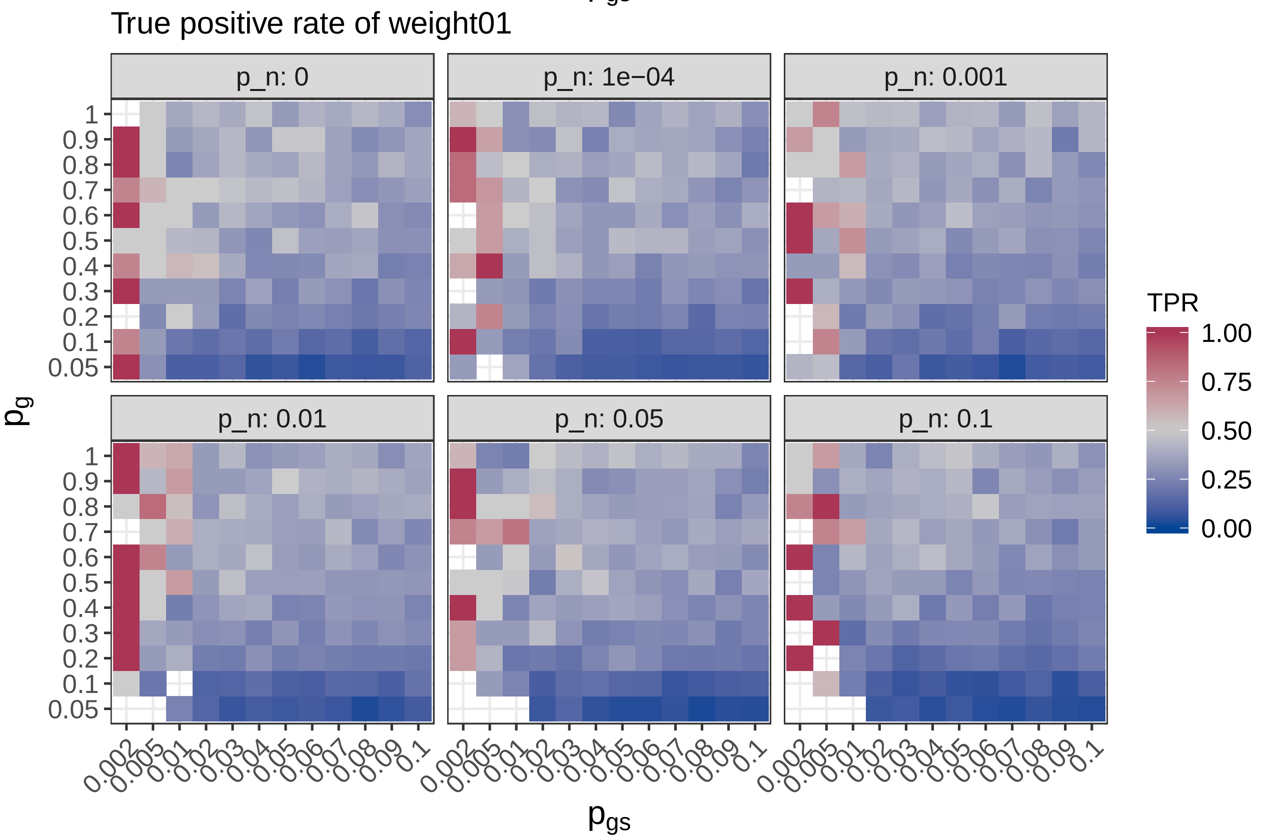
True positive rate of weight01
p_n: 0
p_n: 1e−04
p_n: 0.001
p_n: 0.01
p_n: 0.05
p_n: 0.1
1
0.9
0.8
0.7
0.6
0.5
0.4
0.3
0.2
0.1
0.05
0.002
0.005
0.01
0.02
0.03
0.04
0.05
0.06
0.07
0.08
0.09
0.1
pg
pgs
TPR
1.00
0.75
0.50
0.25
0.00

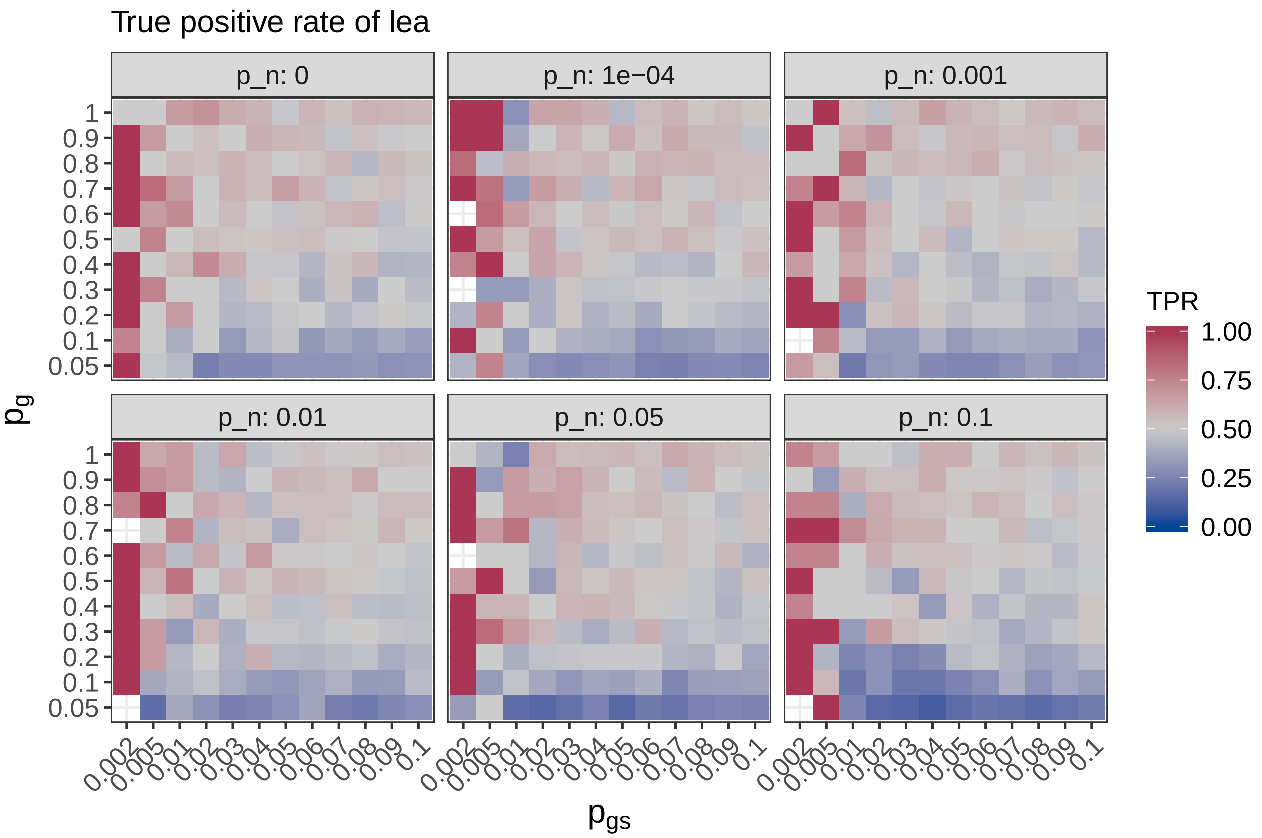

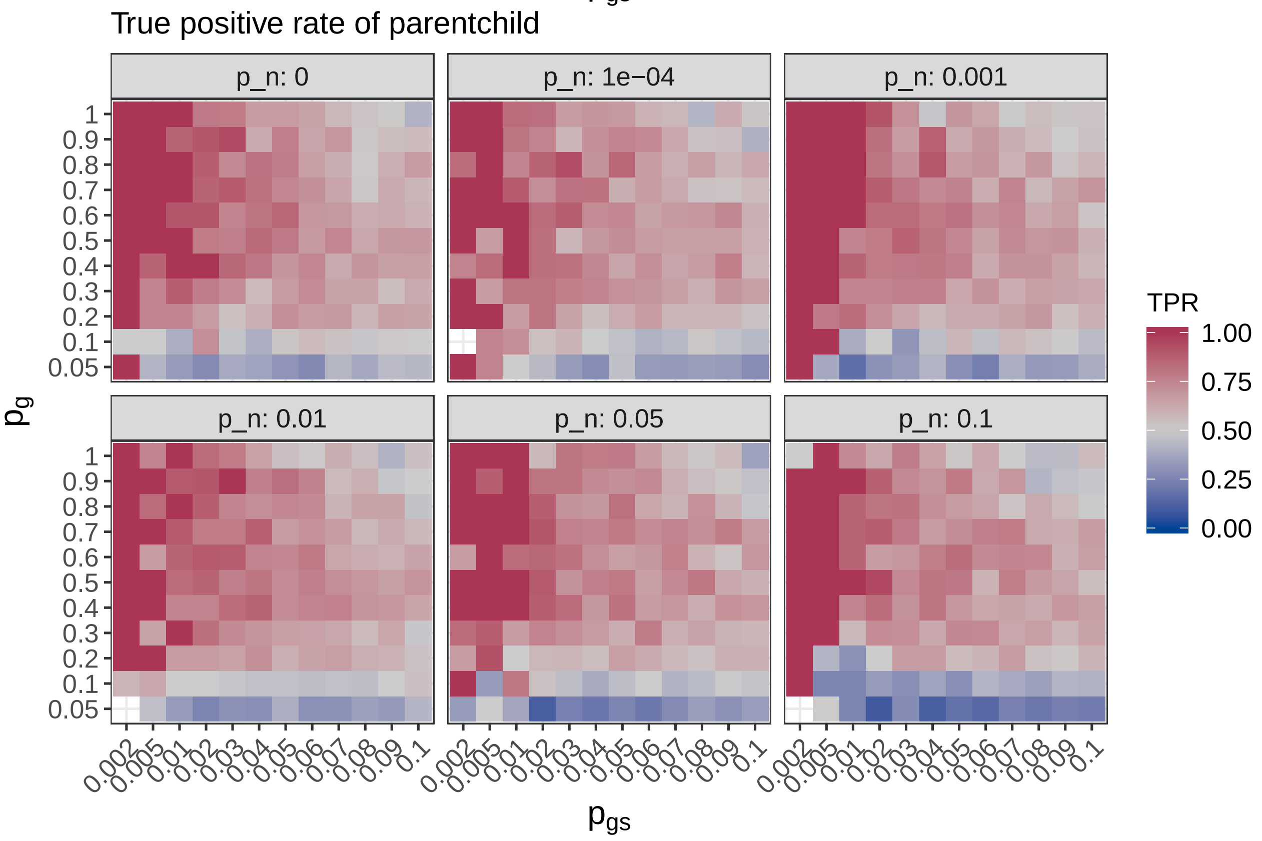

Only the `classic` and `parentchild` methods have a high TPR.

Below we visualize difference in TPR between `gerr` and other methods.

```
diffLowCol <- "#E6AB02"
diffMidCol <- "#CCCCCC"
diffHighCol <- "#4DAF4A"

plot_diff <- function(res, m, stat, label, name) res %>%
  filter(method %in% c(m, 'gerr')) %>%
  select(p_gs, p_g, p_n, method, !!sym(stat)) %>%
  spread(method, !!sym(stat)) %>%
  mutate(diff=gerr-!!sym(m)) %>%
  ggplot(aes(x=factor(p_gs), y=factor(p_g), fill=diff)) +
  facet_wrap(~p_n, labeller = label_both) +
  geom_tile() +
  scale_fill_gradient2(low=diffLowCol, mid=diffMidCol, high=diffHighCol, midpoint=0,
                       limits=c(-0.5, 0.5), oob=scales::squish,
                       name=bquote(Delta~.(label))) +
  theme(axis.text.x = element_text(angle=45, hjust=1)) +
  ggtitle(str_glue("{name} difference (gerr - {m})", name=name, m=m)) +
  xlab(expression(p[gs])) + ylab(expression(p[g]))

walk(topgo_methods, ~print(plot_diff(
  full_sim_res_agg,
  .,
  'tpr',
  'TPR',
  'True positive rate')))
```

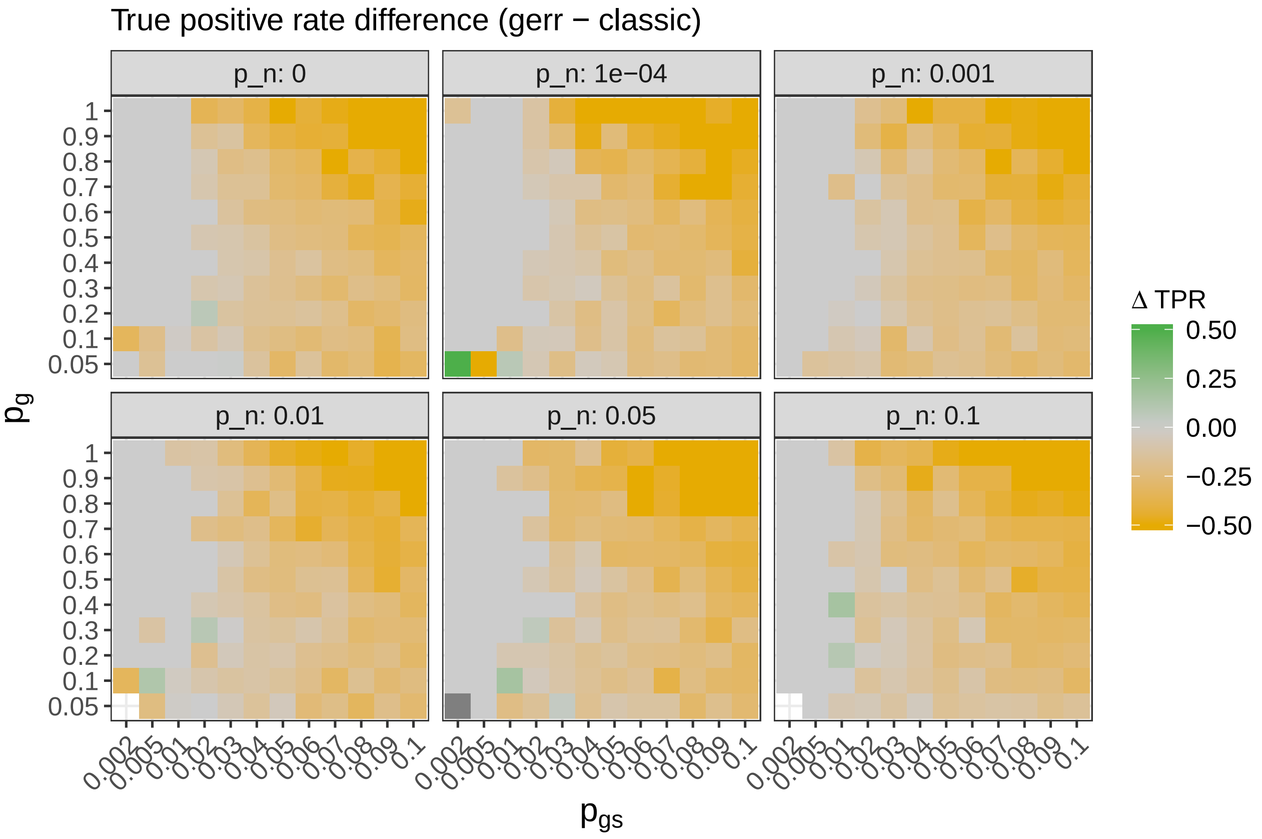

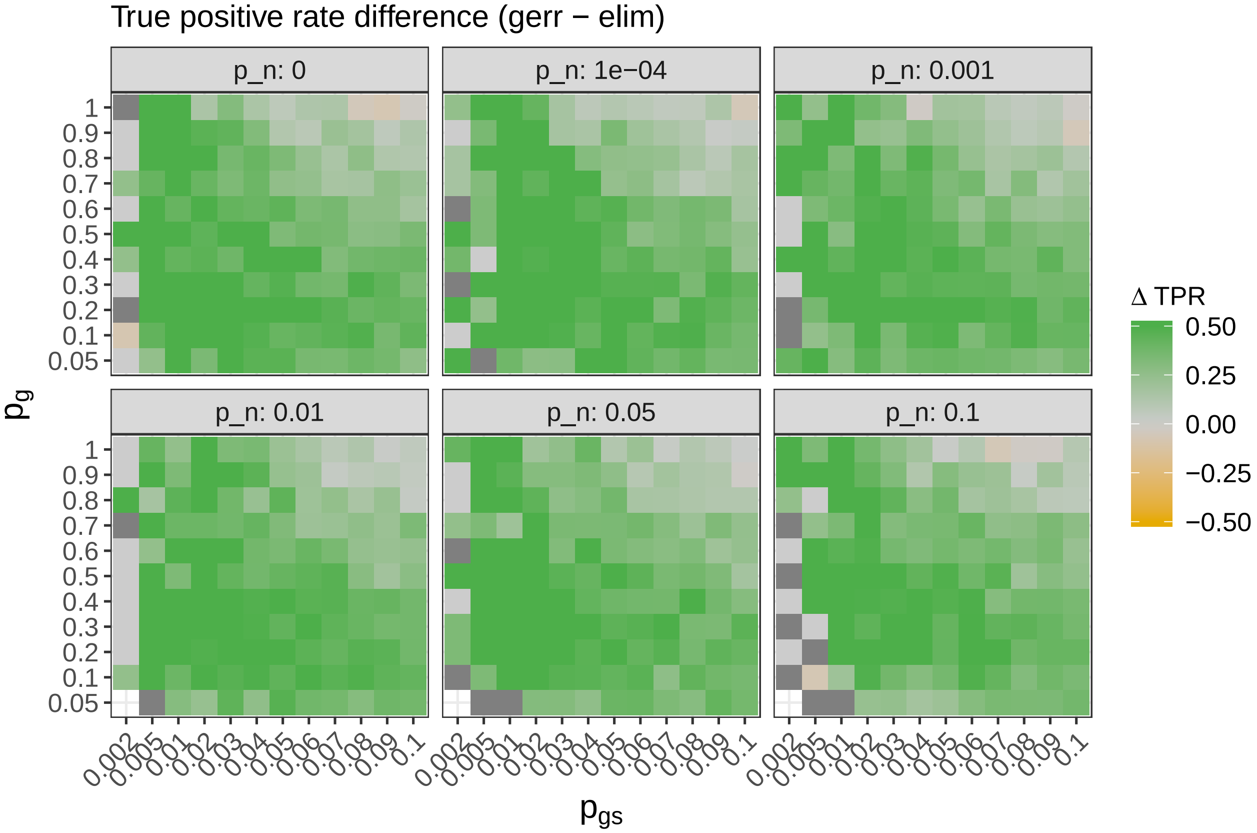
True positive rate difference (gerr − elim)
p_n: 0
p_n: 1e−04
p_n: 0.001
p_n: 0.01
p_n: 0.05
p_n: 0.1
$p_g$
$p_{gs}$
1 0.9 0.8 0.7 0.6 0.5 0.4 0.3 0.2 0.1 0.05
0.002 0.005 0.01 0.02 0.03 0.04 0.05 0.06 0.07 0.08 0.09 0.1
Δ TPR
0.50
0.25
0.00
−0.25
−0.50

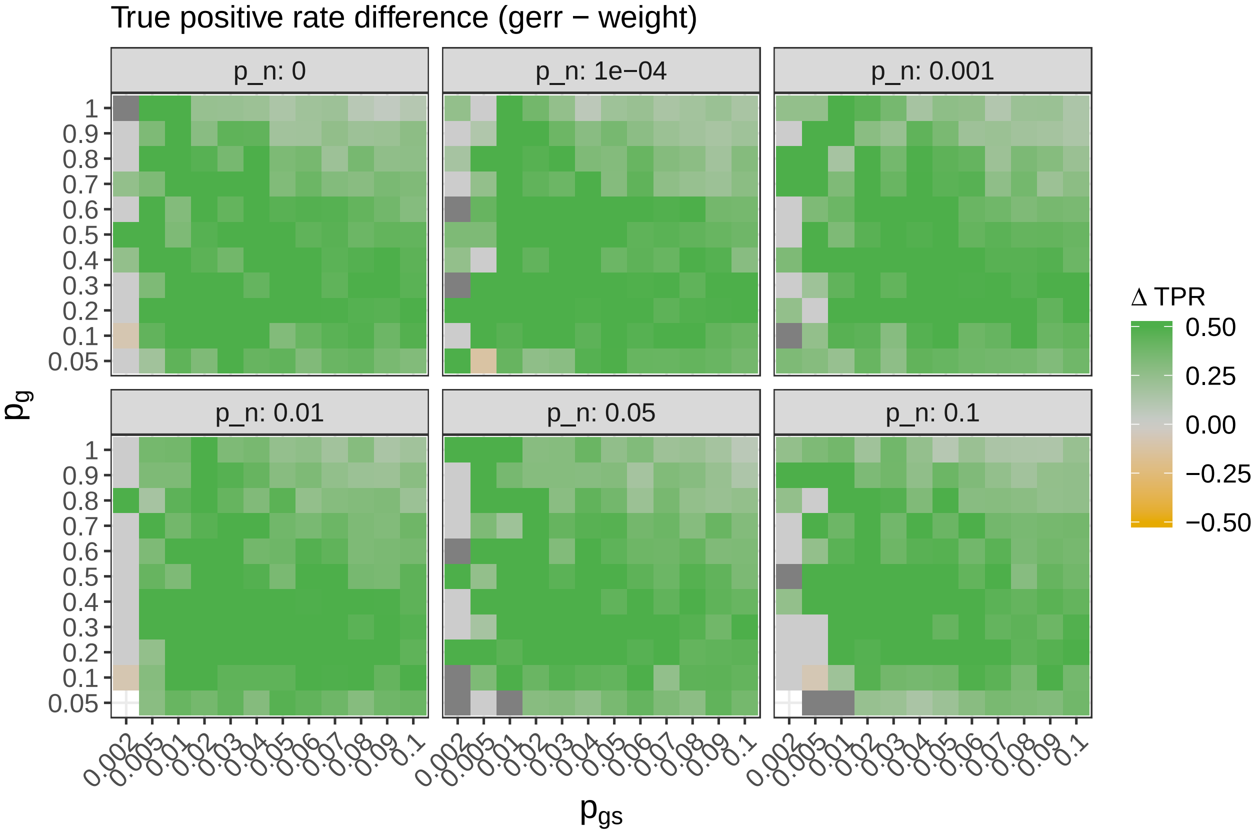
True positive rate difference (gerr − weight)
p_n: 0
p_n: 1e−04
p_n: 0.001
p_n: 0.01
p_n: 0.05
p_n: 0.1
$p_g$
$p_{gs}$
1 0.9 0.8 0.7 0.6 0.5 0.4 0.3 0.2 0.1 0.05
0.002 0.005 0.01 0.02 0.03 0.04 0.05 0.06 0.07 0.08 0.09 0.1
Δ TPR
0.50
0.25
0.00
−0.25
−0.50

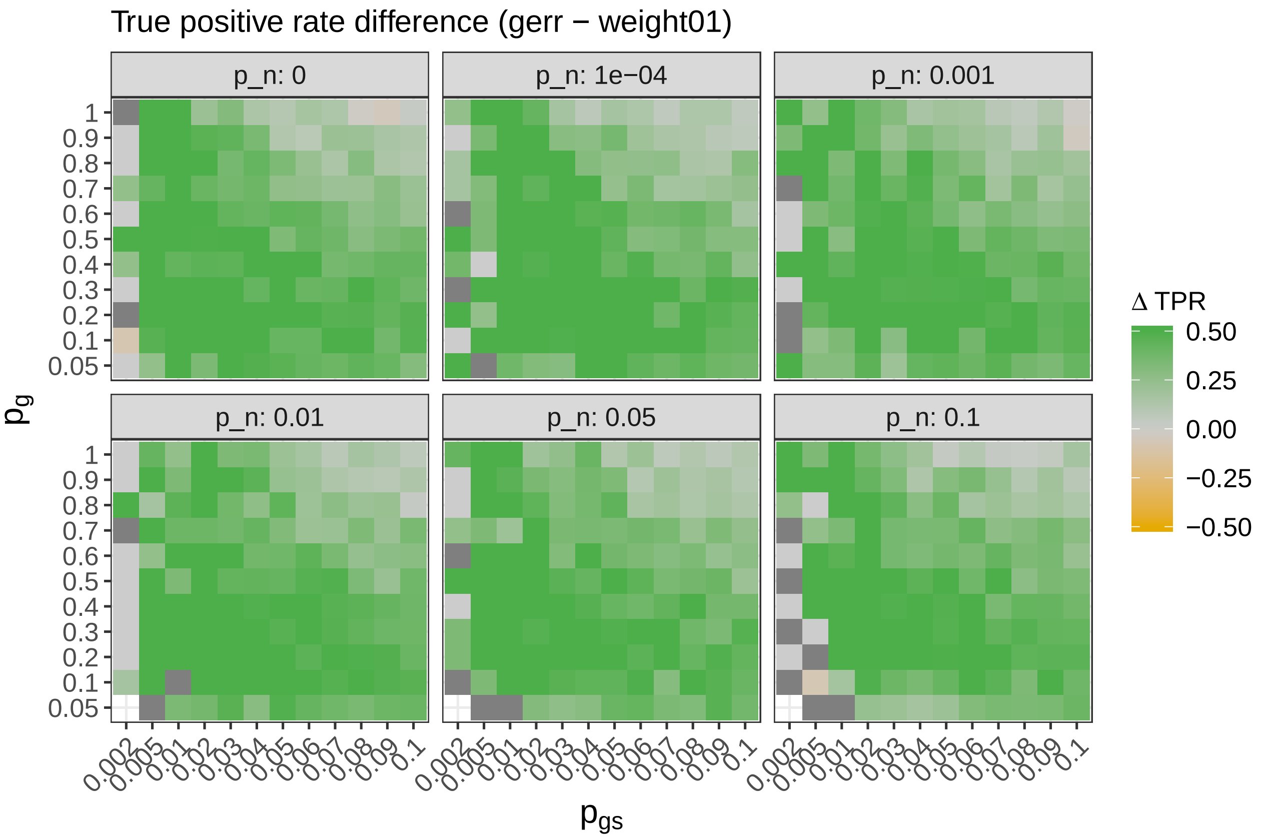
True positive rate difference (gerr − weight01)
p_n: 0
p_n: 1e−04
p_n: 0.001
p_n: 0.01
p_n: 0.05
p_n: 0.1
Δ TPR
0.50
0.25
0.00
−0.25
−0.50
$p_g$
$p_{gs}$
1 0.9 0.8 0.7 0.6 0.5 0.4 0.3 0.2 0.1 0.05
0.002 0.005 0.01 0.02 0.03 0.04 0.05 0.06 0.07 0.08 0.09 0.1

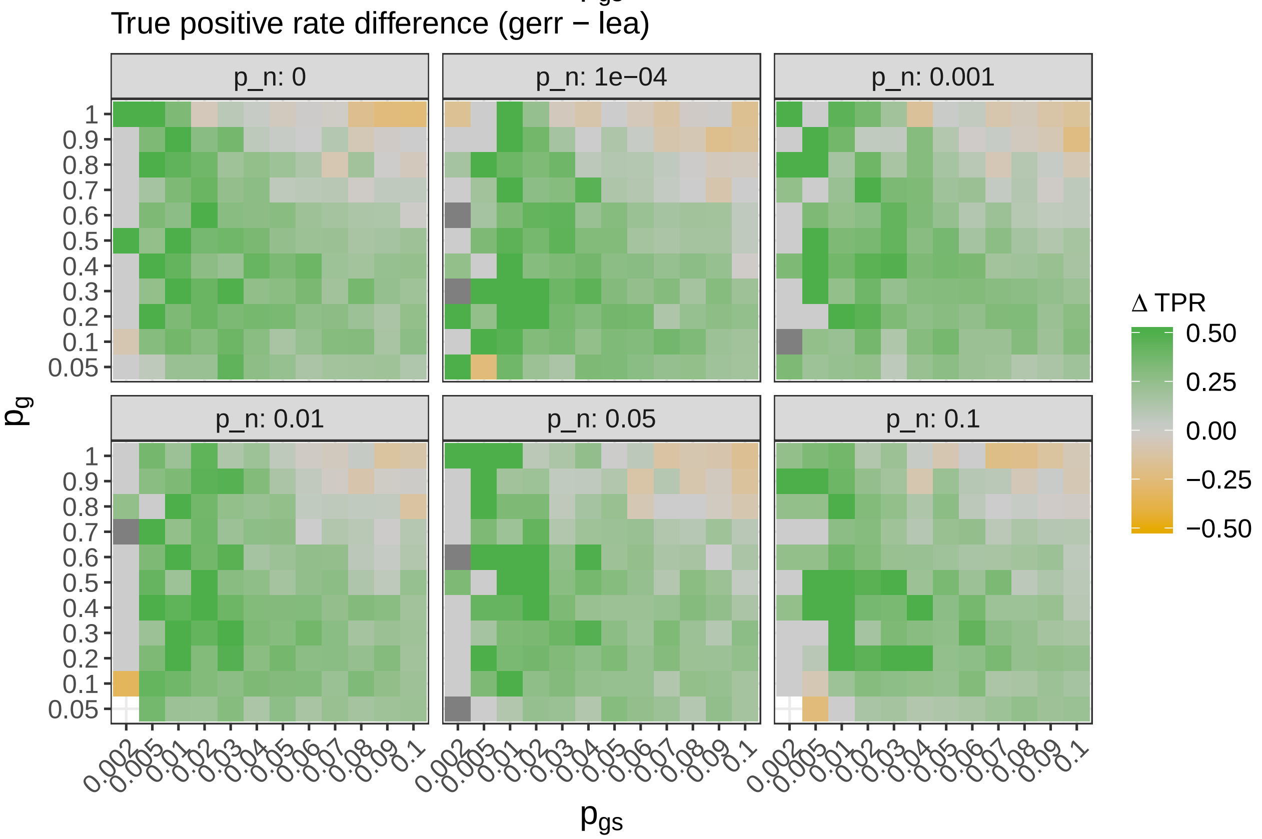
True positive rate difference (gerr − lea)
p_n: 0
p_n: 1e−04
p_n: 0.001
p_n: 0.01
p_n: 0.05
p_n: 0.1
Δ TPR
0.50
0.25
0.00
−0.25
−0.50
$p_g$
$p_{gs}$
1 0.9 0.8 0.7 0.6 0.5 0.4 0.3 0.2 0.1 0.05
0.002 0.005 0.01 0.02 0.03 0.04 0.05 0.06 0.07 0.08 0.09 0.1

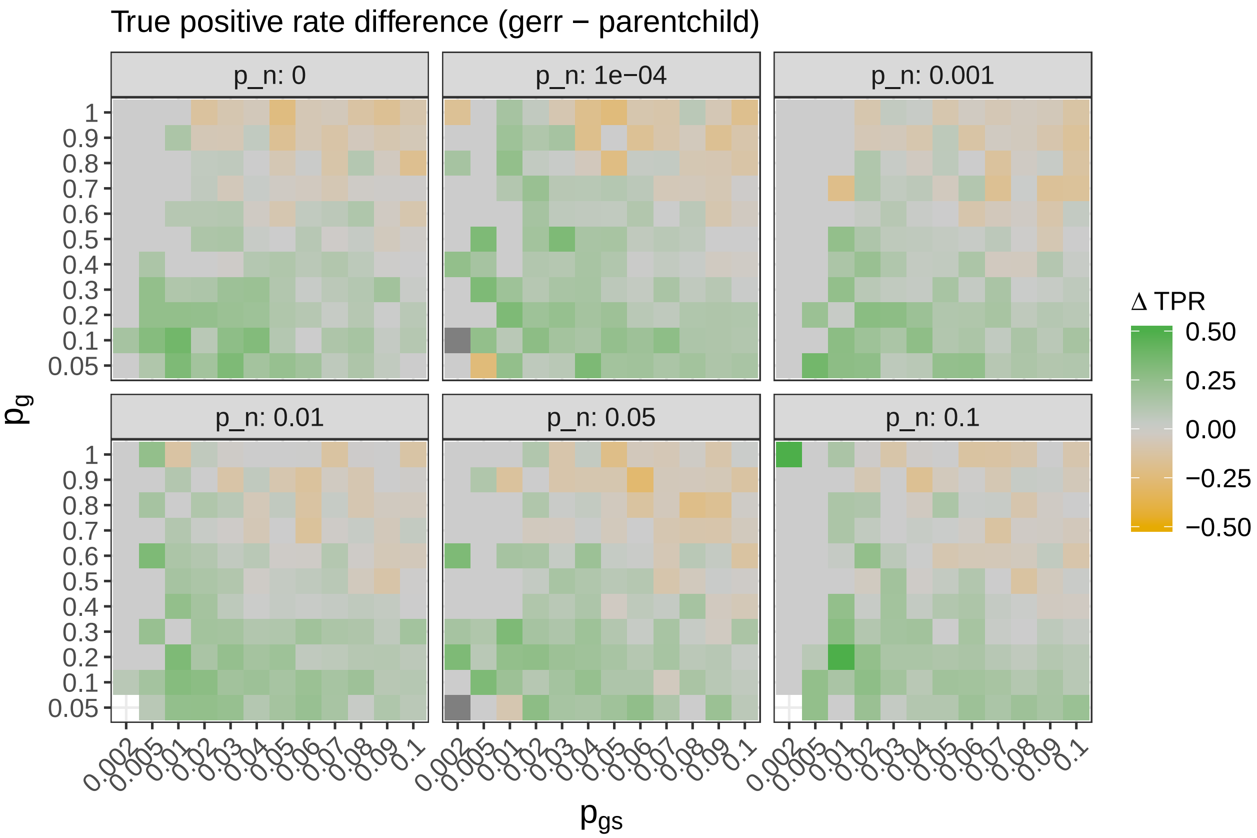

Only the `classic` and has the higher true positive rate. The `parentchild` method have a comparable TPR.

#### Specificity, or 1-false positive rate (FPR)

Next we examine specificity, which equals 1-false-positive rate. First, we visualize the variation precision of `gerr` by varying parameters.

```
plot_stat(full_sim_res_agg, 'gerr', 'm_fpr', '1-FPR',
          'False positive rate')
```

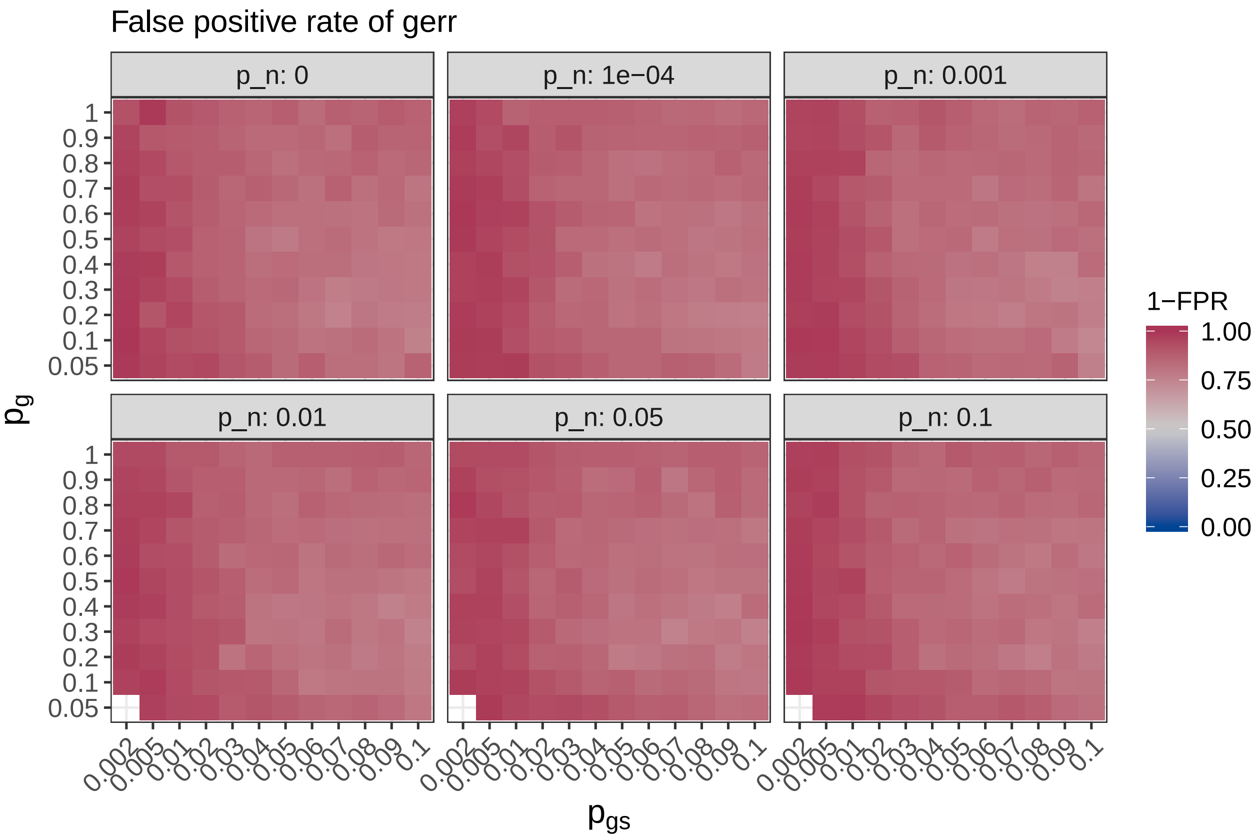

The specificity of `gerr` seems quite robust against the choice of $p_n$. It decreases when $p_{gs}$ is high, namely when many gene-sets contribute to GOI. Likely it is because of calling other gene-sets that are partially redundant as false-positive hits, as shown in the previous verification step.

Next we reveal how FPR changes for other methods.

```
walk(topgo_methods, ~print(plot_stat(
  full_sim_res_agg,
  .,
  'm_fpr',
  '1-FPR',
  'False positive rate')))
```

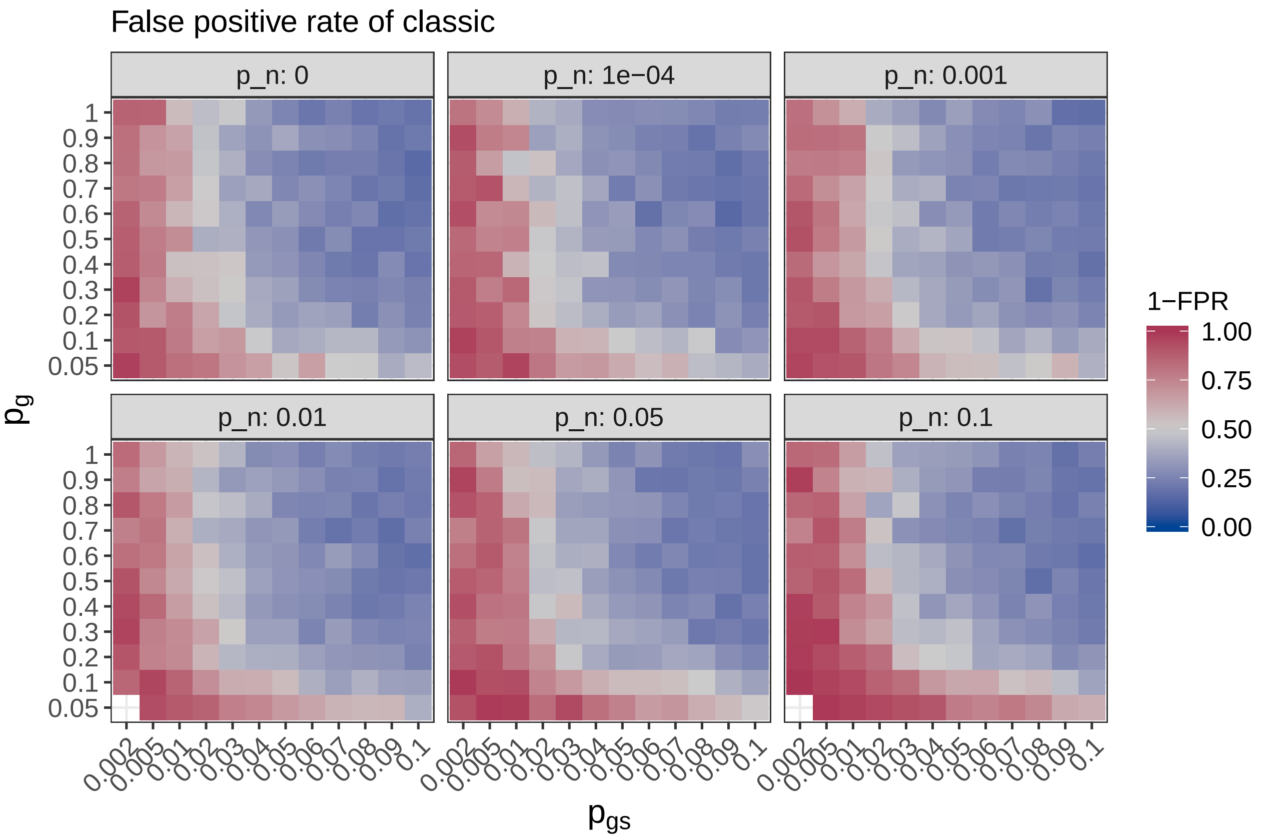
False positive rate of classic
p_n: 0
p_n: 1e−04
p_n: 0.001
p_n: 0.01
p_n: 0.05
p_n: 0.1
$p_g$
$p_{gs}$
1−FPR
1.00
0.75
0.50
0.25
0.00

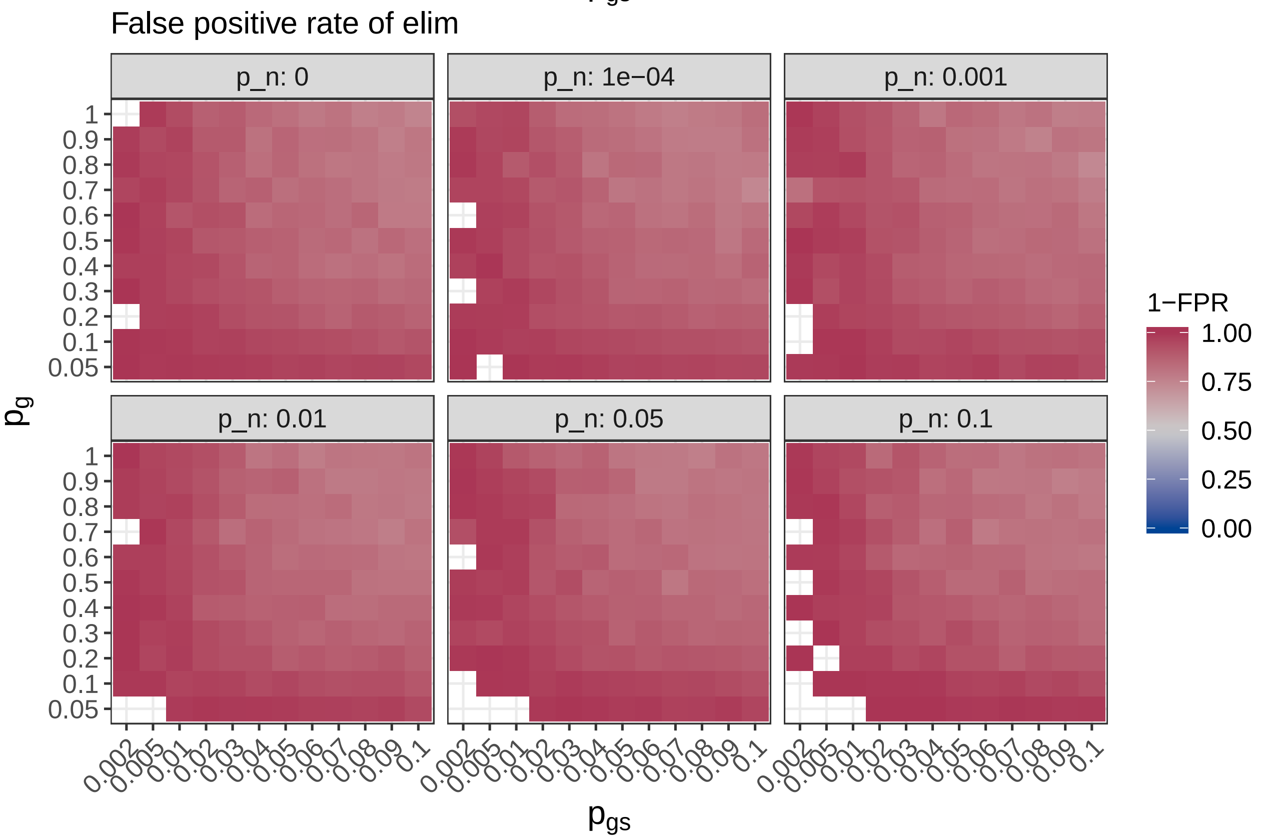
False positive rate of elim
p_n: 0
p_n: 1e−04
p_n: 0.001
p_n: 0.01
p_n: 0.05
p_n: 0.1
$p_g$
$p_{gs}$
1−FPR
1.00
0.75
0.50
0.25
0.00

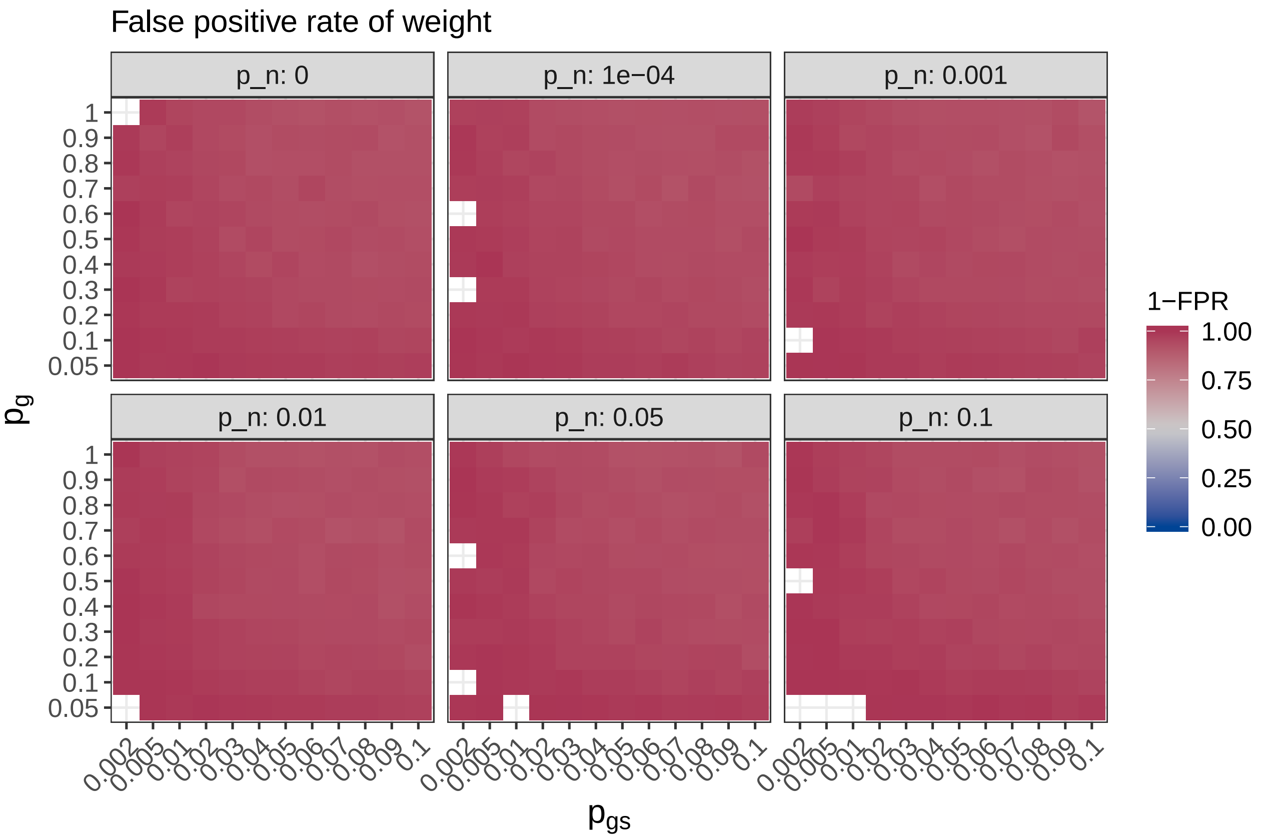
False positive rate of weight
p_n: 0
p_n: 1e−04
p_n: 0.001
p_n: 0.01
p_n: 0.05
p_n: 0.1
$p_g$
$p_{gs}$
1−FPR
1.00
0.75
0.50
0.25
0.00

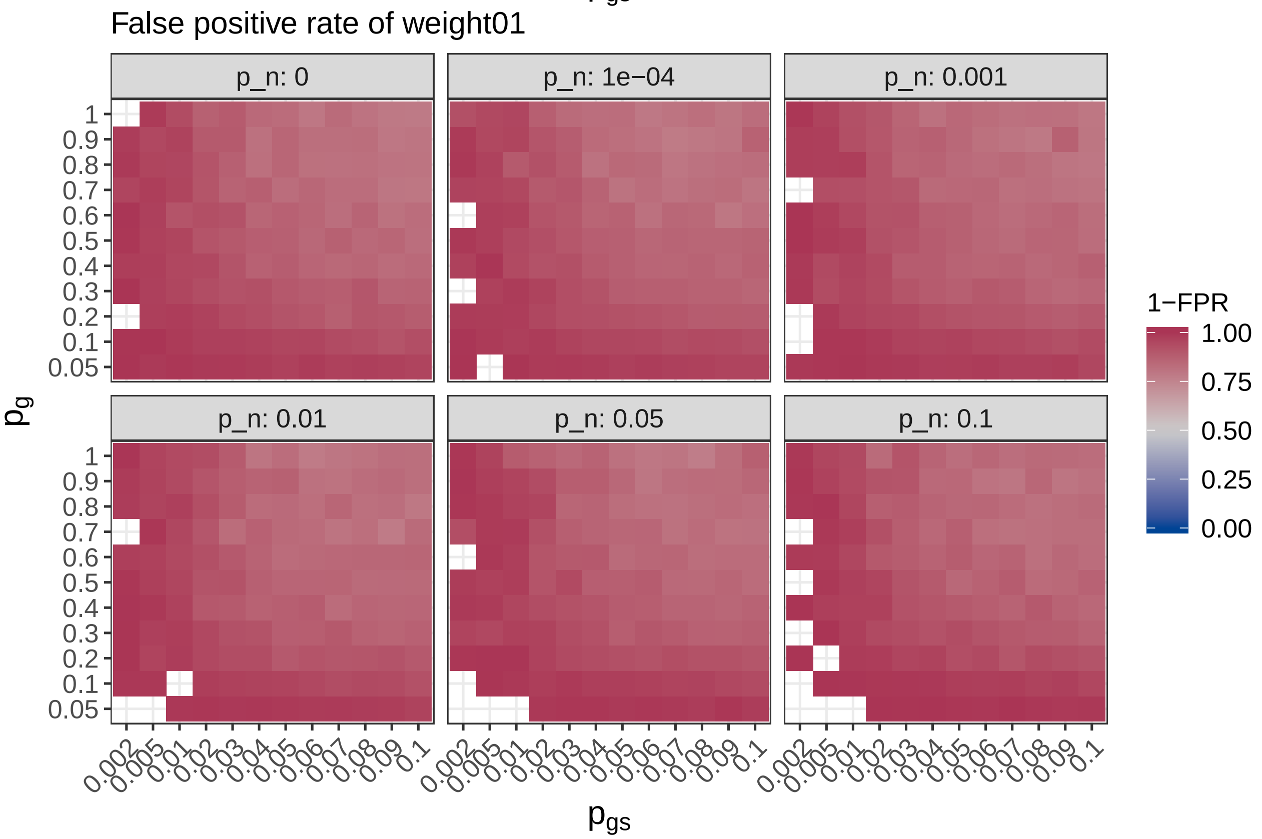
False positive rate of weight01
p_n: 0
p_n: 1e−04
p_n: 0.001
p_n: 0.01
p_n: 0.05
p_n: 0.1
$p_g$
$p_{gs}$
1−FPR
1.00
0.75
0.50
0.25
0.00

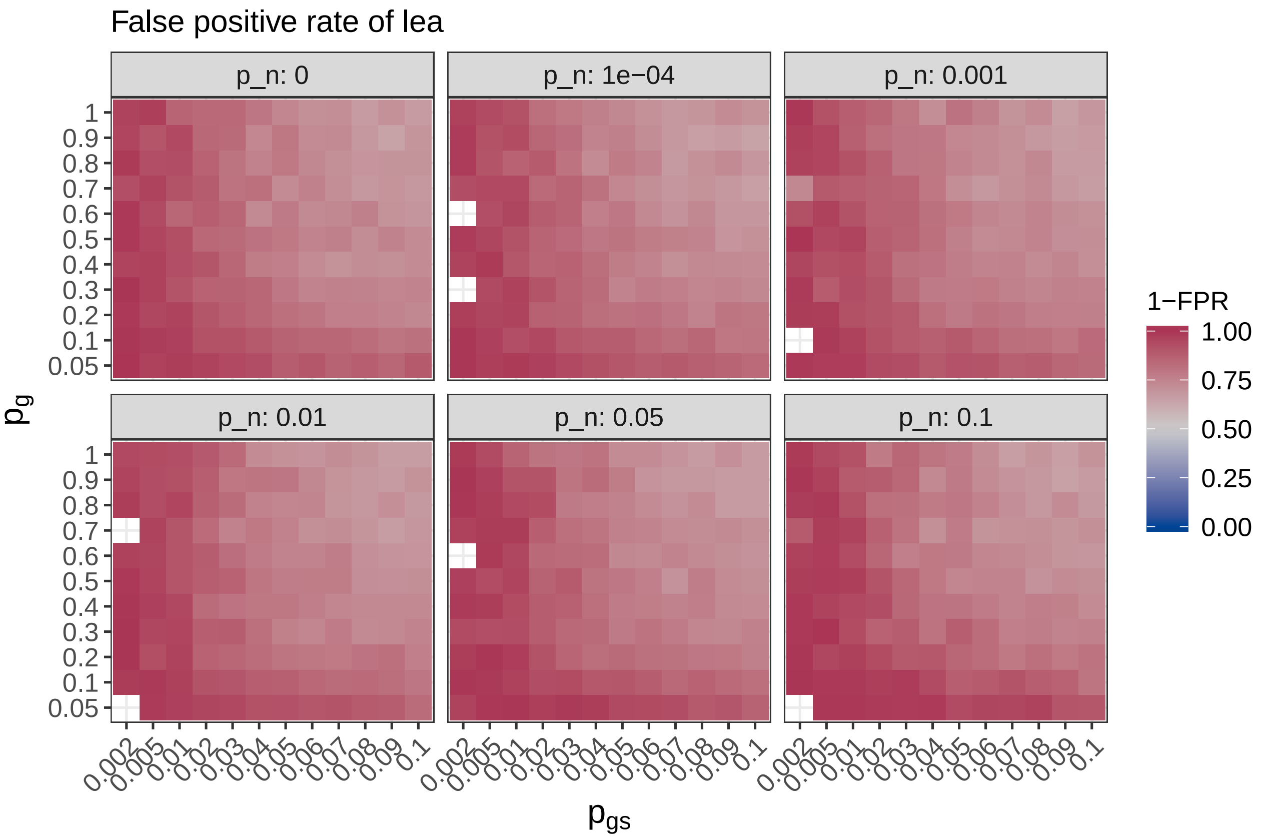

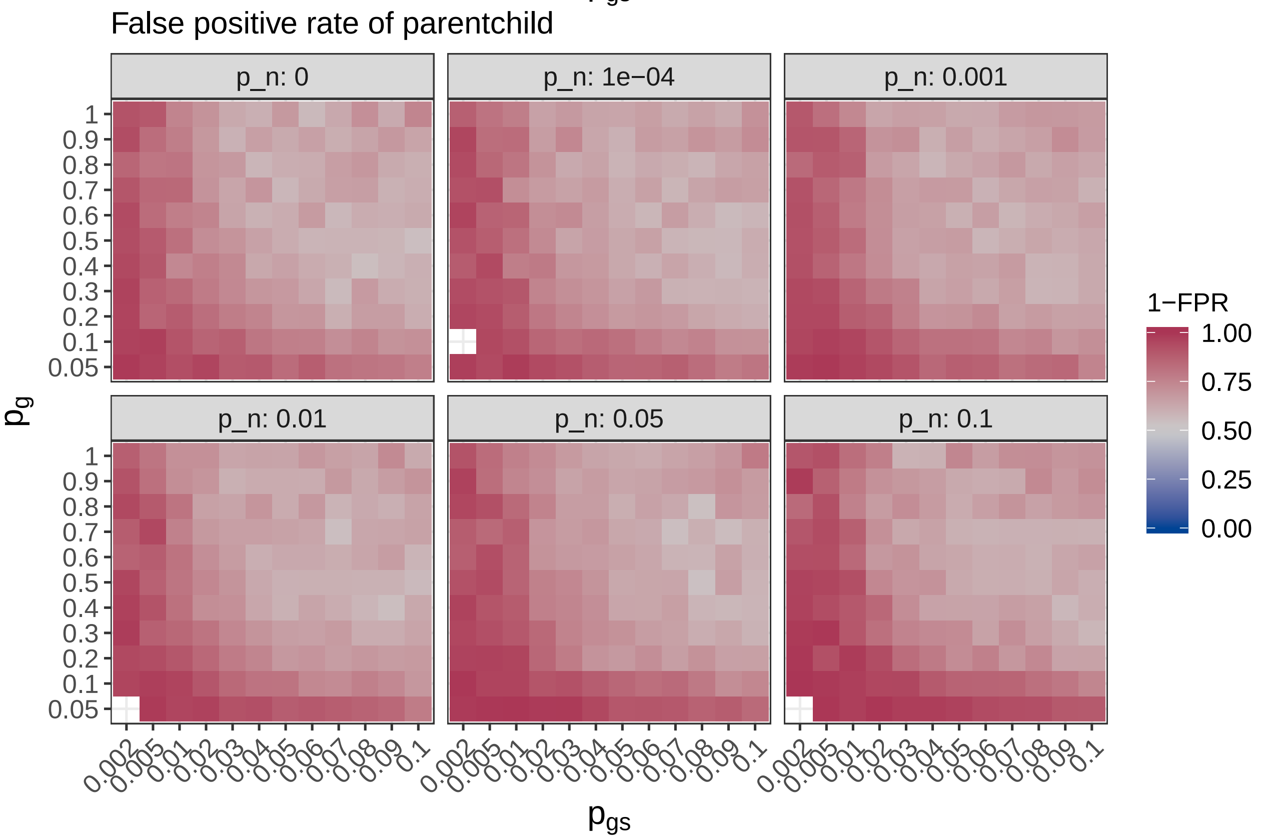

The `classic` method (aka FET+FDR) has the worst problems with the false positive rates, as it does not account for graph structure and gene set redundancy at all. Other methods have a much better FPR.

Now we look at the difference in specificity between `gerr` and other methods.

```
walk(topgo_methods, ~print(plot_diff(
  full_sim_res_agg,
  .,
  'm_fpr',
  '1-FPR',
  'False positive rate')))
```

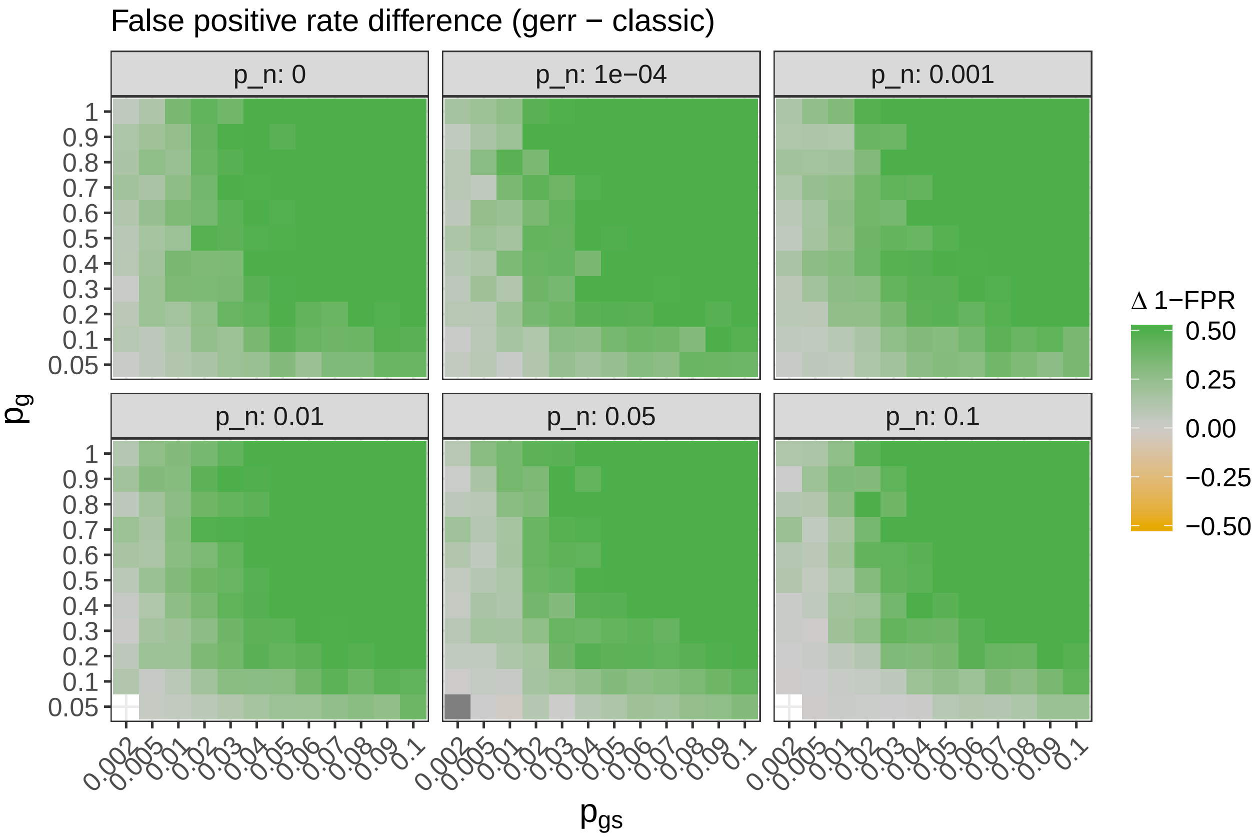

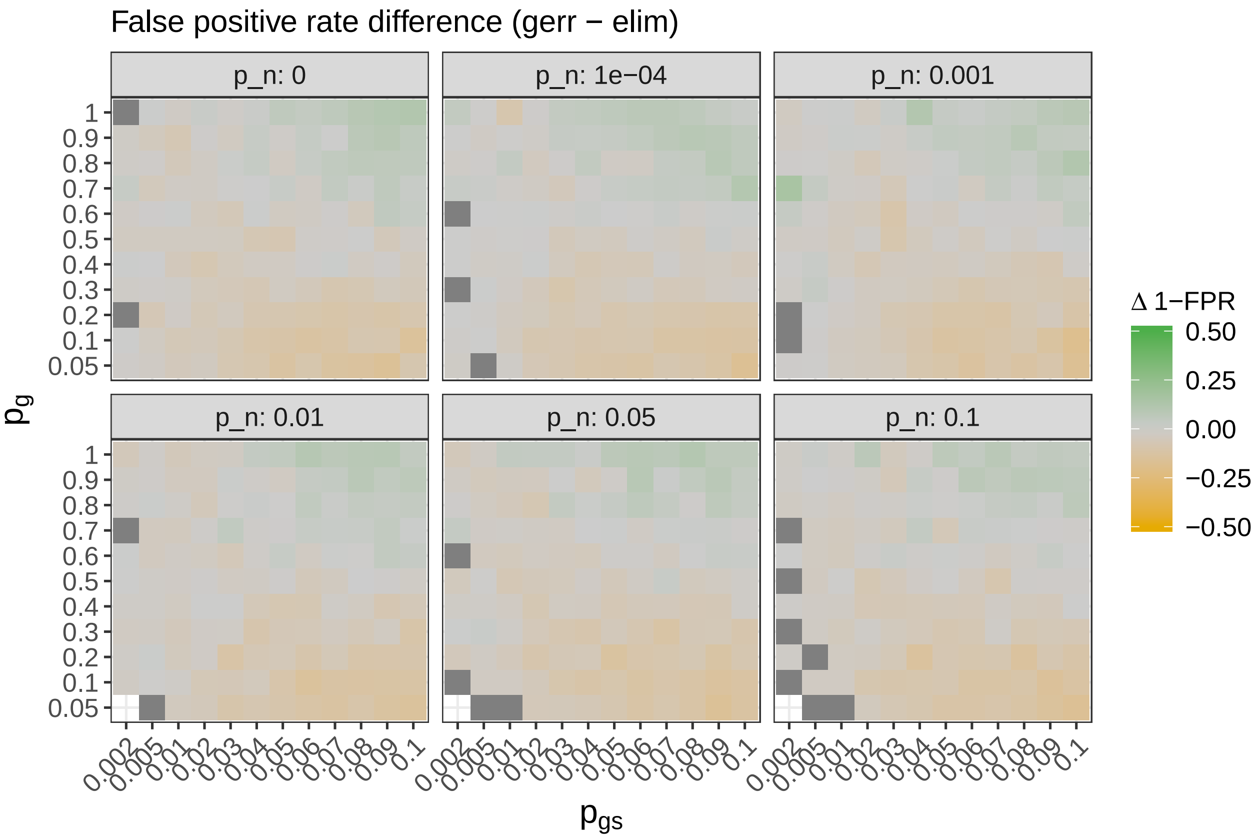
False positive rate difference (gerr − elim)
p_n: 0
p_n: 1e−04
p_n: 0.001
p_n: 0.01
p_n: 0.05
p_n: 0.1
1
0.9
0.8
0.7
0.6
0.5
0.4
0.3
0.2
0.1
0.05
0.002
0.005
0.01
0.02
0.03
0.04
0.05
0.06
0.07
0.08
0.09
0.1
p_g
p_gs
Δ 1−FPR
0.50
0.25
0.00
−0.25
−0.50

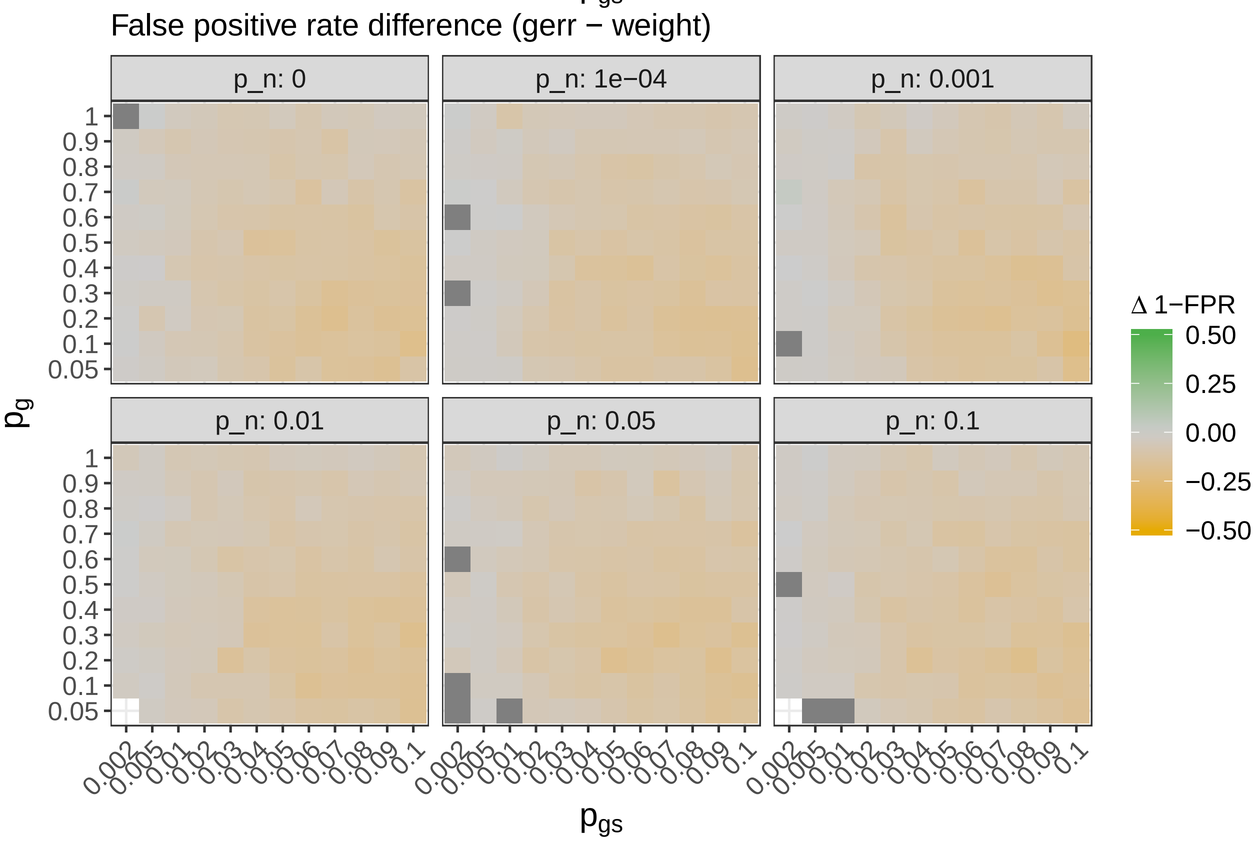
False positive rate difference (gerr − weight)
p_n: 0
p_n: 1e−04
p_n: 0.001
p_n: 0.01
p_n: 0.05
p_n: 0.1
1
0.9
0.8
0.7
0.6
0.5
0.4
0.3
0.2
0.1
0.05
0.002
0.005
0.01
0.02
0.03
0.04
0.05
0.06
0.07
0.08
0.09
0.1
p_g
p_gs
Δ 1−FPR
0.50
0.25
0.00
−0.25
−0.50

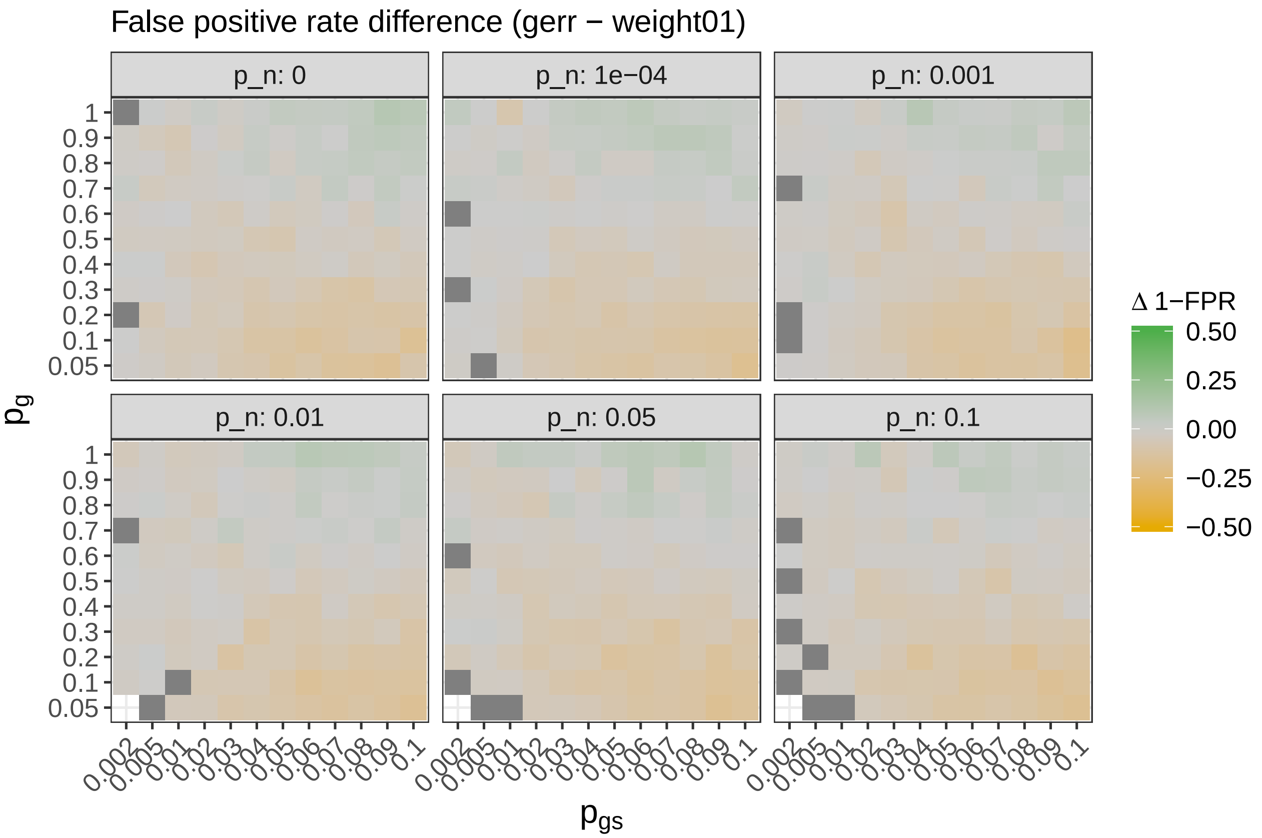
False positive rate difference (gerr − weight01)
p_n: 0
p_n: 1e−04
p_n: 0.001
p_n: 0.01
p_n: 0.05
p_n: 0.1
1
0.9
0.8
0.7
0.6
0.5
0.4
0.3
0.2
0.1
0.05
0.002 0.005 0.01 0.02 0.03 0.04 0.05 0.06 0.07 0.08 0.09 0.1
p_g
p_gs
Δ 1−FPR
0.50
0.25
0.00
−0.25
−0.50

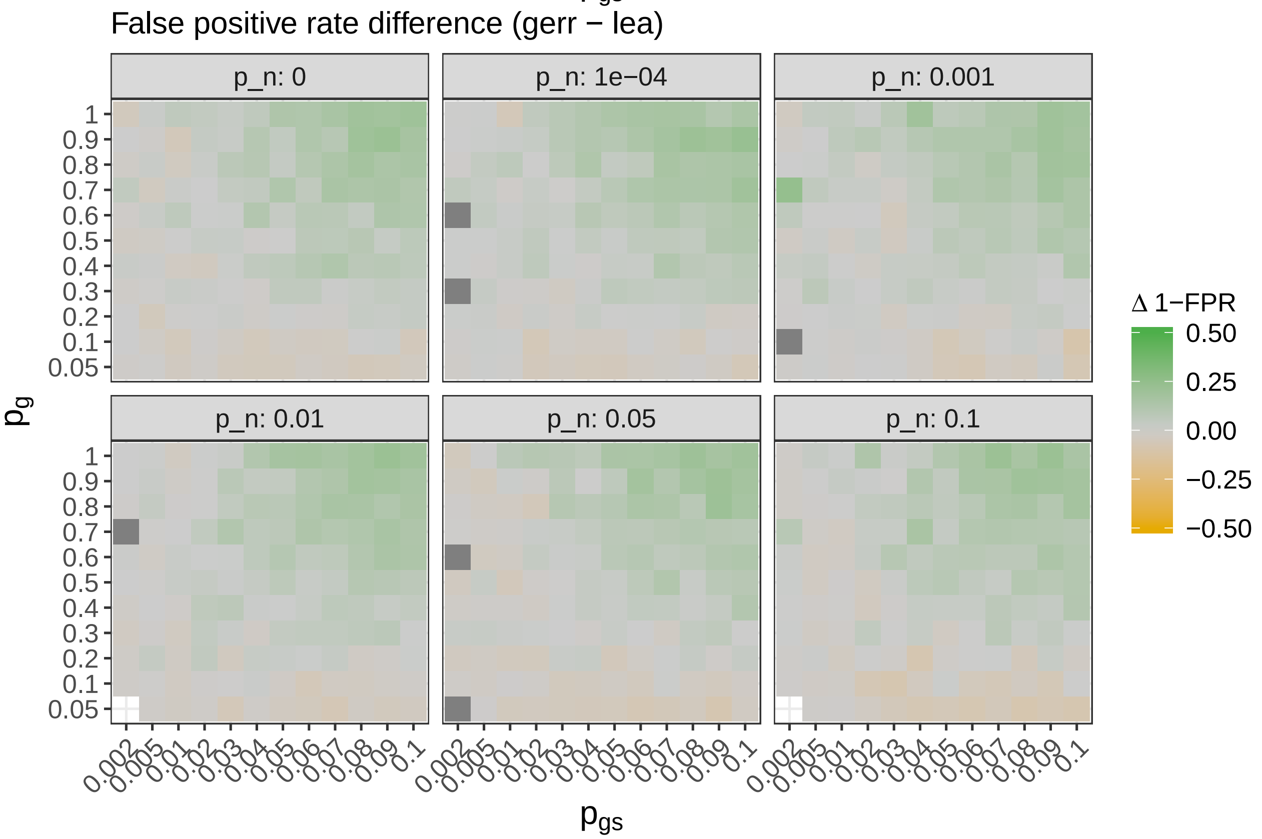
False positive rate difference (gerr − lea)
p_n: 0
p_n: 1e−04
p_n: 0.001
p_n: 0.01
p_n: 0.05
p_n: 0.1
1
0.9
0.8
0.7
0.6
0.5
0.4
0.3
0.2
0.1
0.05
0.002 0.005 0.01 0.02 0.03 0.04 0.05 0.06 0.07 0.08 0.09 0.1
p_g
p_gs
Δ 1−FPR
0.50
0.25
0.00
−0.25
−0.50

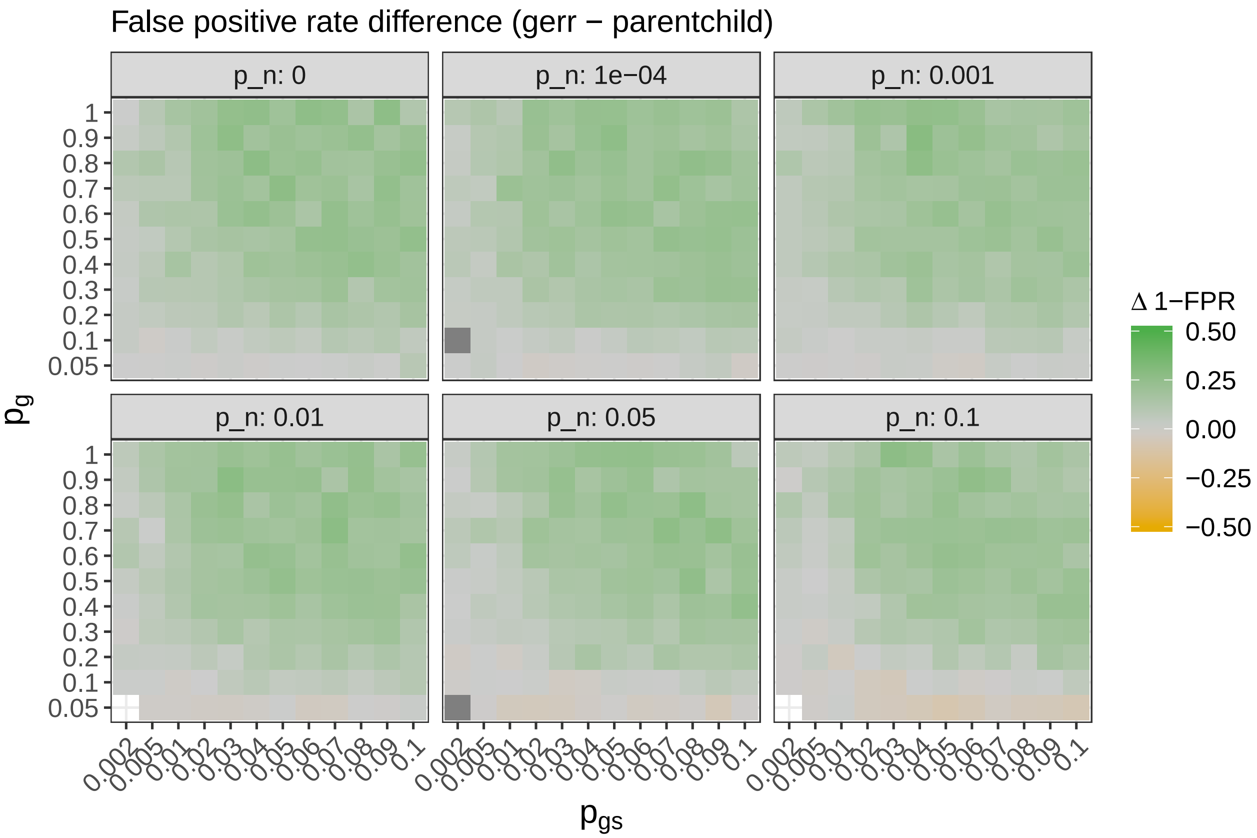

The method implemented in `gerr` shows higher specificity compared to `classic` and `parentchild`. For other methods the results are comparable.

#### $F_1$ score

$F_1$ score is the harmonic mean of precision and sensitivity and therefore a good measure of balanced performance. Below we visualize the $F_1$ score of `gerr` results.

```
plot_stat(full_sim_res_agg, 'gerr', 'f1', 'F1', 'F1 score')
```

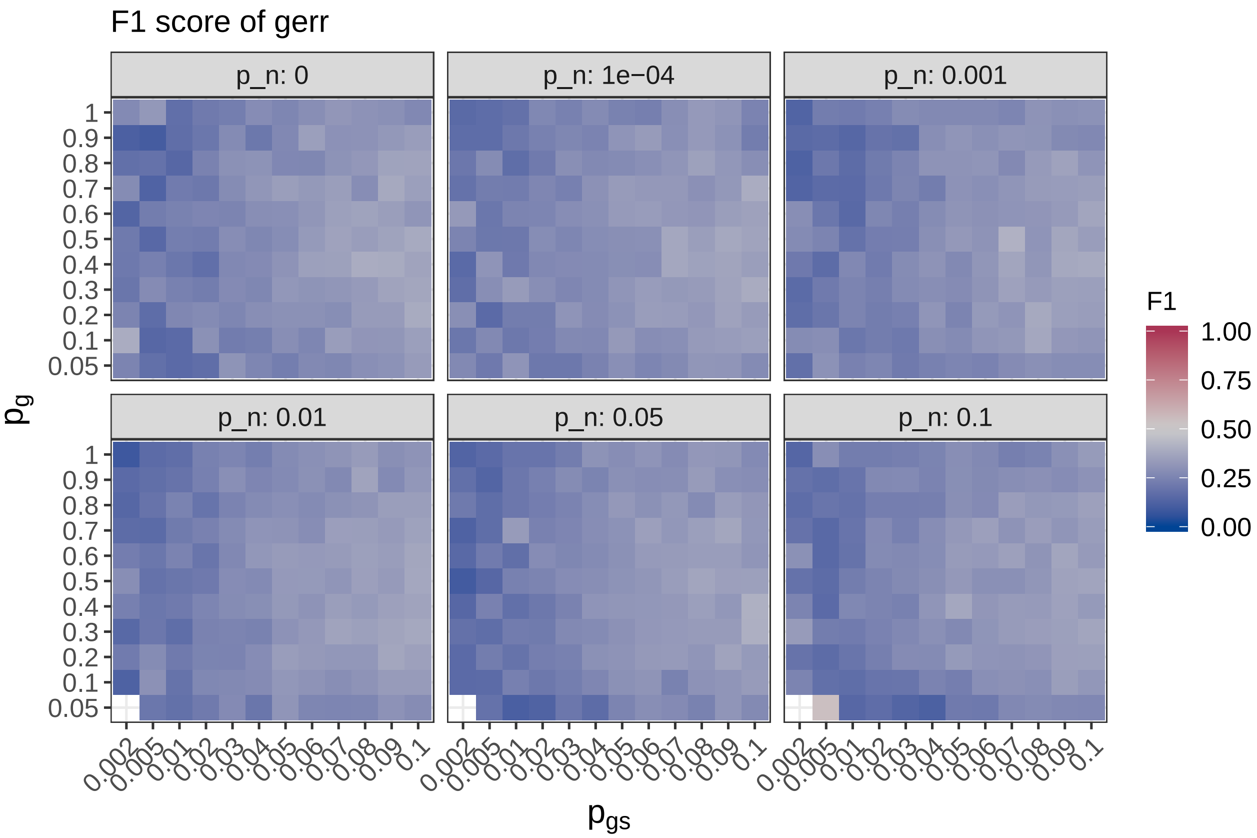

It seems that $F_1$ score of `gerr` is much lower than in simulations from MSigDB. This is probably because of higher redundancy of the GO-graph.

Next we visualize the $F_1$ scores of other methods.

```
walk(topgo_methods, ~print(plot_stat(
  full_sim_res_agg,
  .,
  'f1',
  'F1',
  'F1 score')))
```

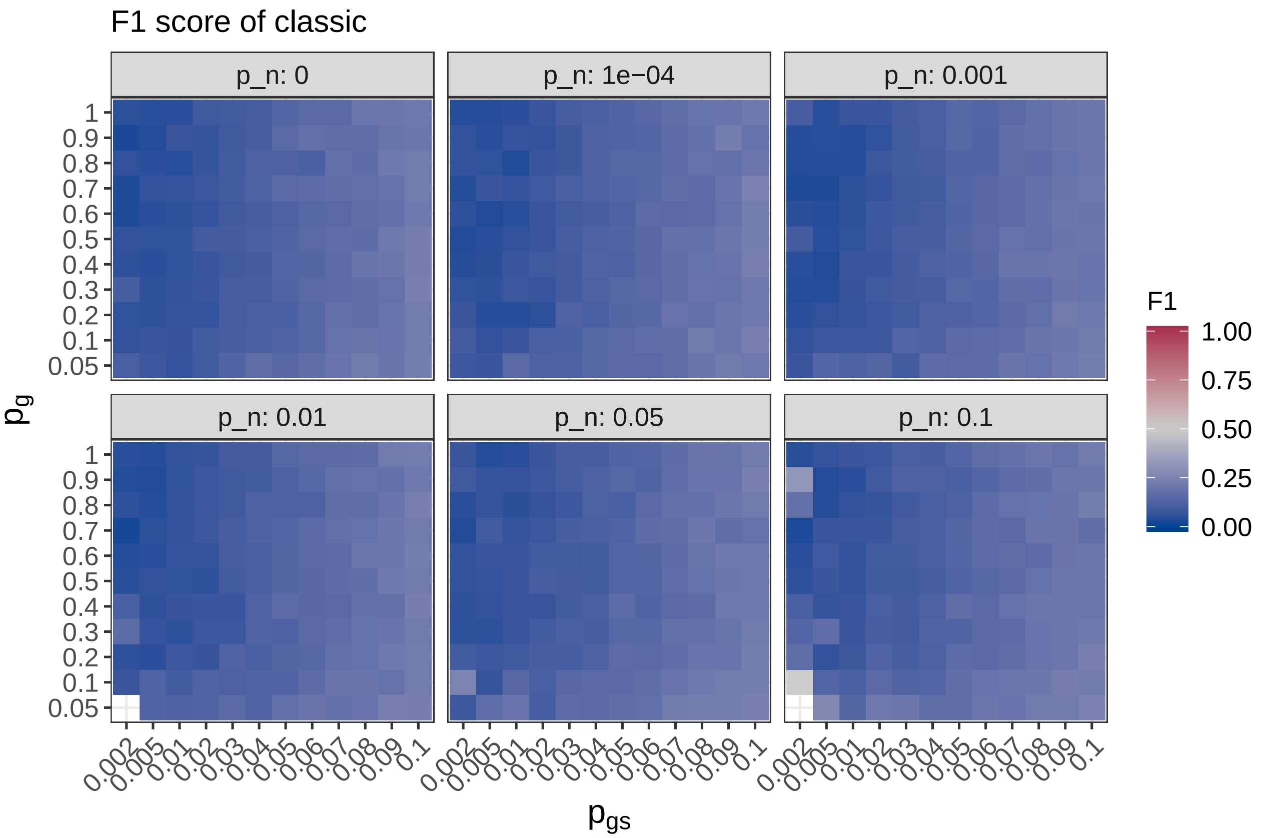
F1 score of classic
p_n: 0
p_n: 1e−04
p_n: 0.001
p_n: 0.01
p_n: 0.05
p_n: 0.1
$p_g$
$p_{gs}$
F1
1.00
0.75
0.50
0.25
0.00

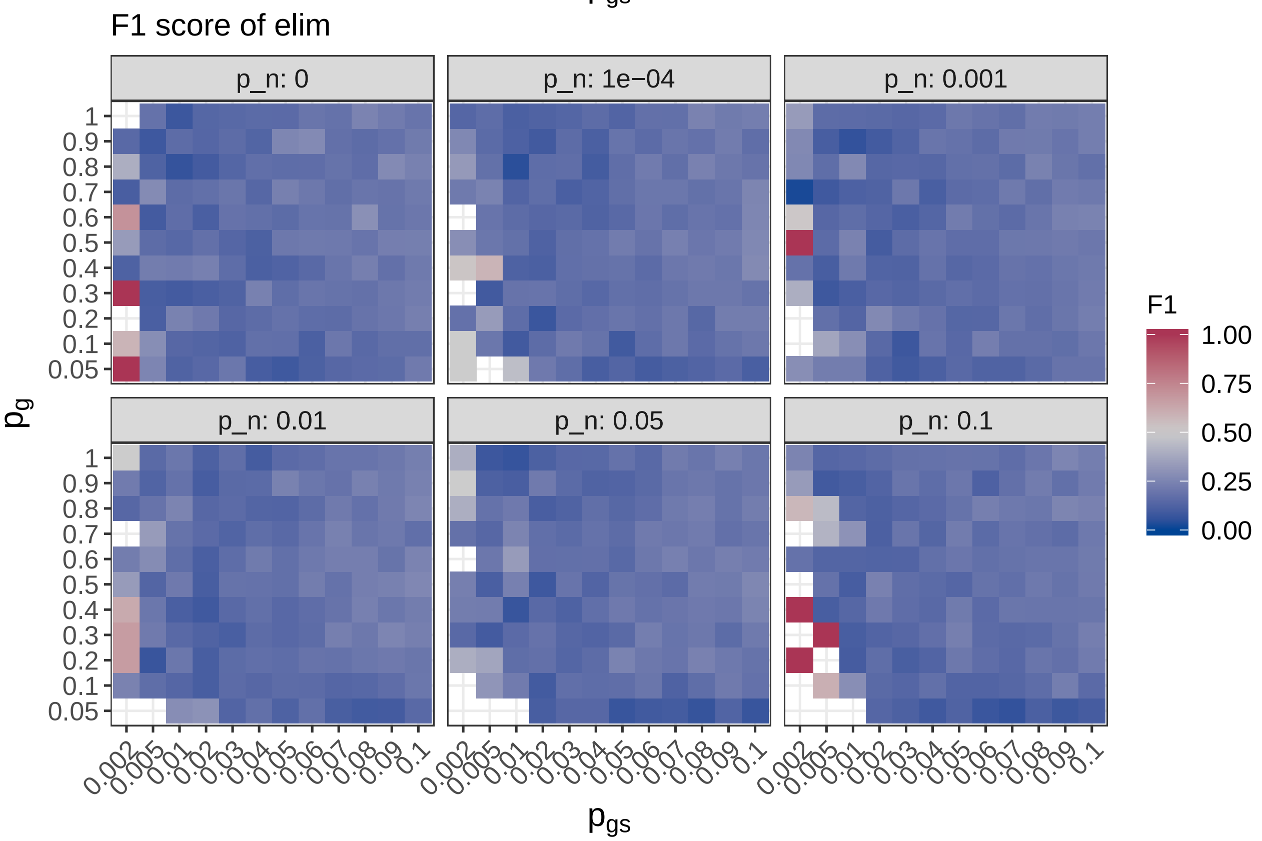
F1 score of elim
p_n: 0
p_n: 1e−04
p_n: 0.001
p_n: 0.01
p_n: 0.05
p_n: 0.1
$p_g$
$p_{gs}$
F1
1.00
0.75
0.50
0.25
0.00

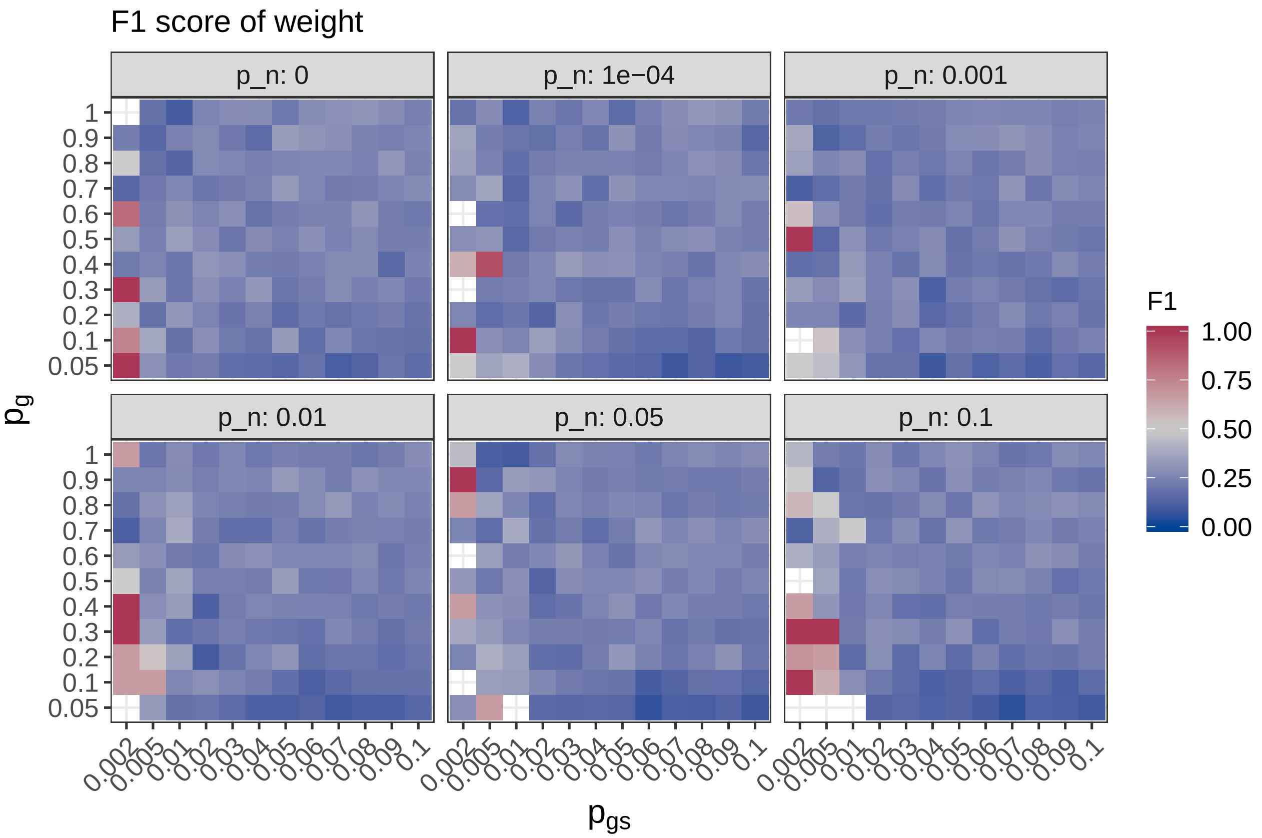
F1 score of weight
p_n: 0
p_n: 1e−04
p_n: 0.001
p_n: 0.01
p_n: 0.05
p_n: 0.1
1
0.9
0.8
0.7
0.6
0.5
0.4
0.3
0.2
0.1
0.05
0.002
0.005
0.01
0.02
0.03
0.04
0.05
0.06
0.07
0.08
0.09
0.1
$p_g$
$p_{gs}$
F1
1.00
0.75
0.50
0.25
0.00

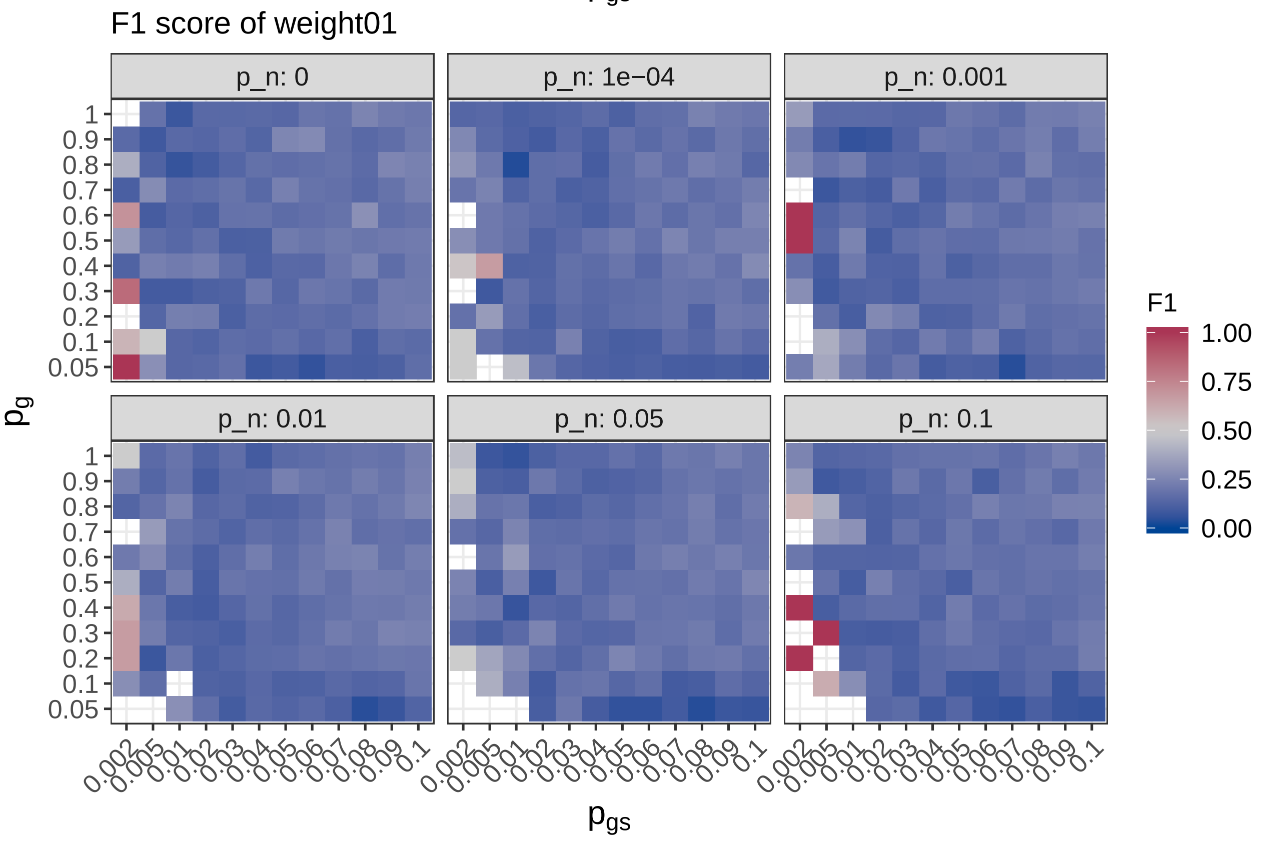
F1 score of weight01
p_n: 0
p_n: 1e−04
p_n: 0.001
p_n: 0.01
p_n: 0.05
p_n: 0.1
1
0.9
0.8
0.7
0.6
0.5
0.4
0.3
0.2
0.1
0.05
0.002
0.005
0.01
0.02
0.03
0.04
0.05
0.06
0.07
0.08
0.09
0.1
$p_g$
$p_{gs}$
F1
1.00
0.75
0.50
0.25
0.00

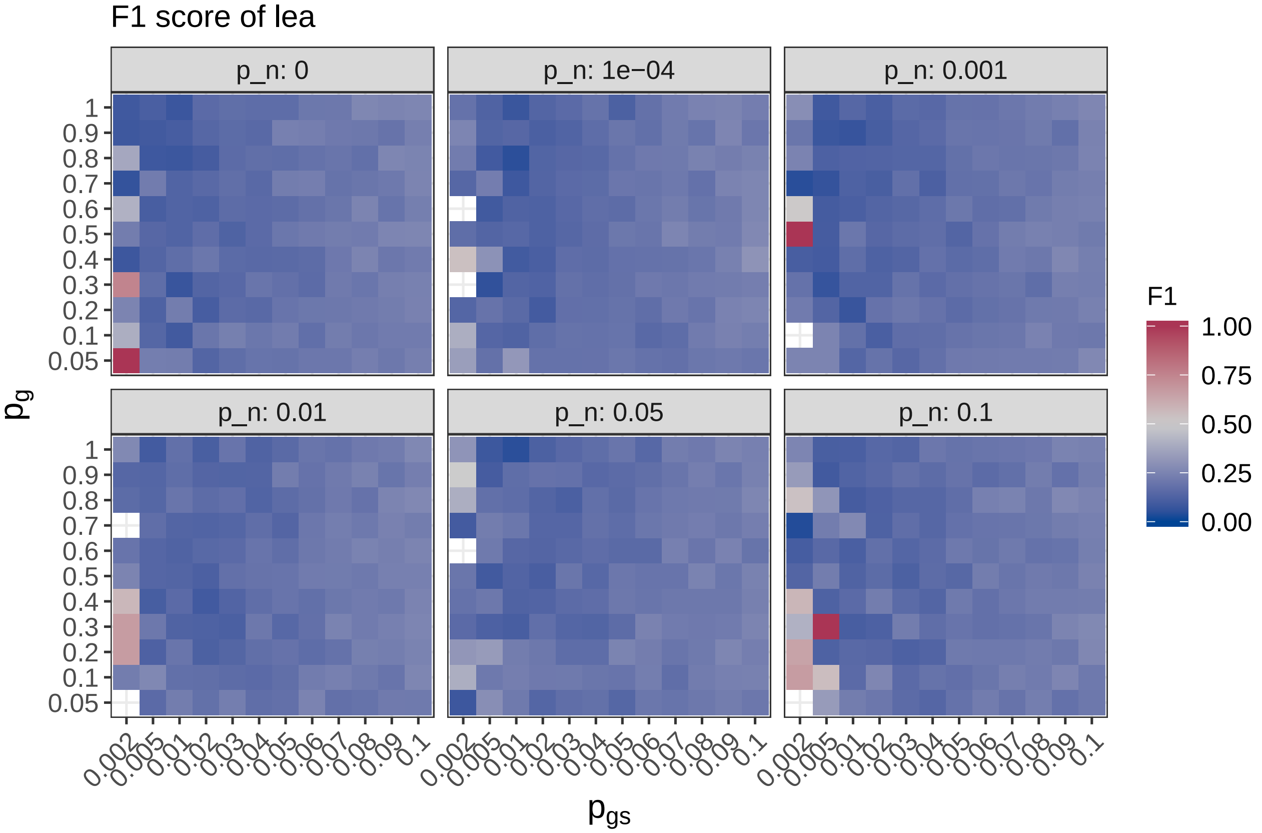

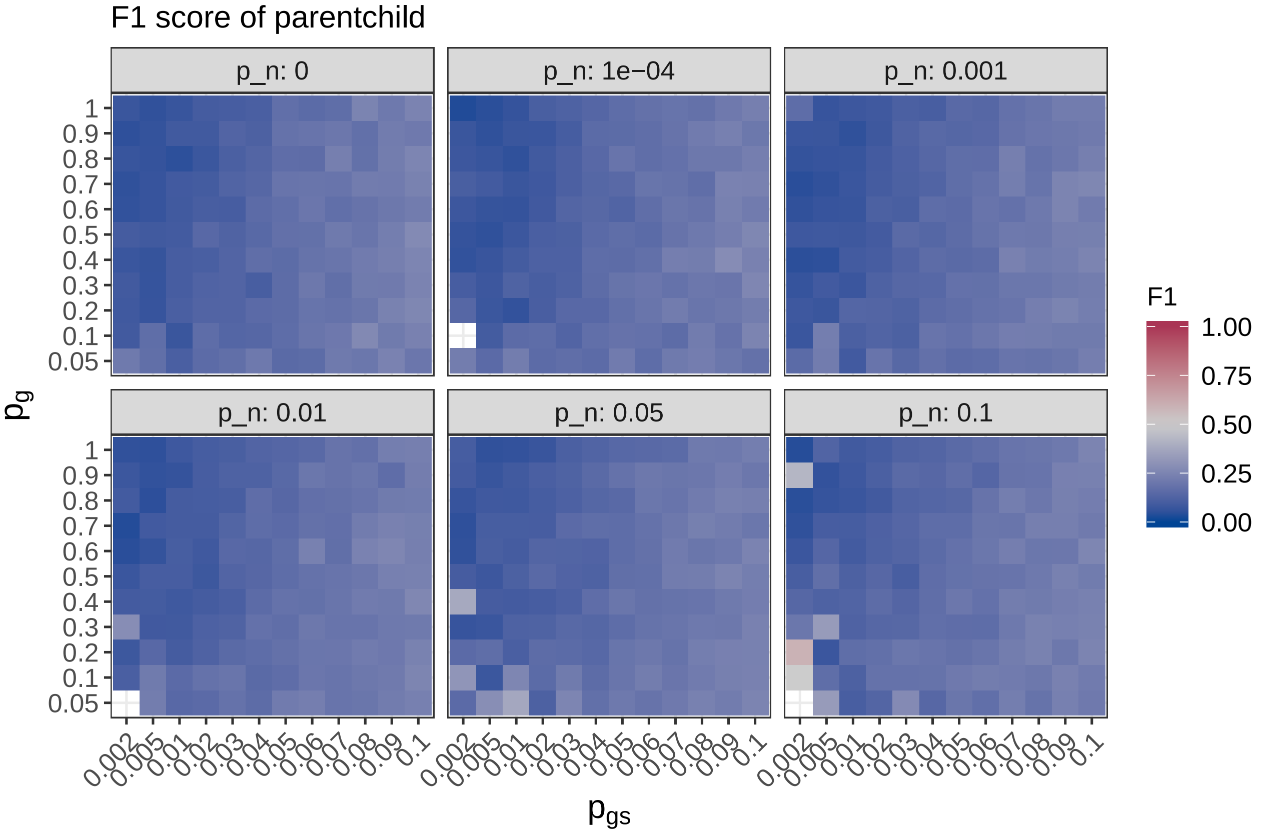

Next we calculate the difference of $F_1$ scores.

```r
walk(topgo_methods, ~print(plot_diff(
  full_sim_res_agg,
  .,
  'f1',
  'F1',
  'F1 score')))
```

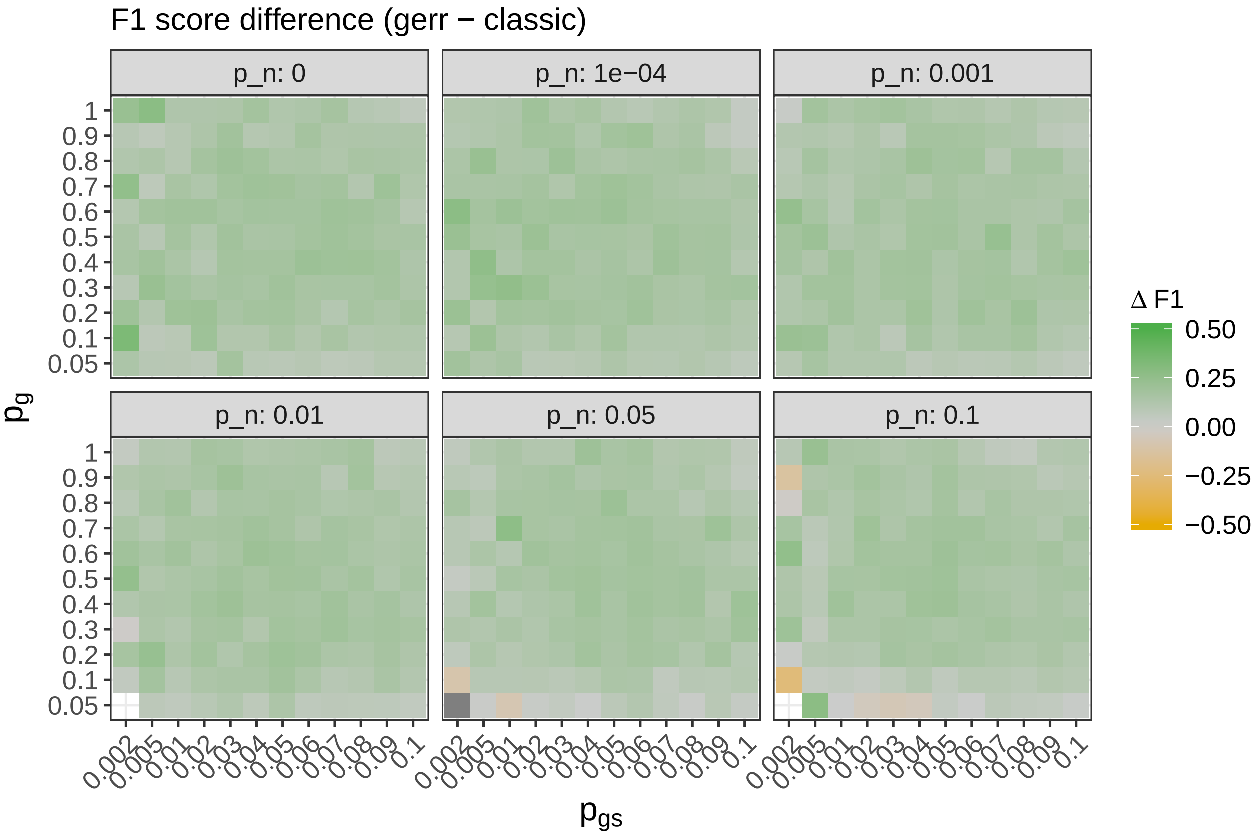

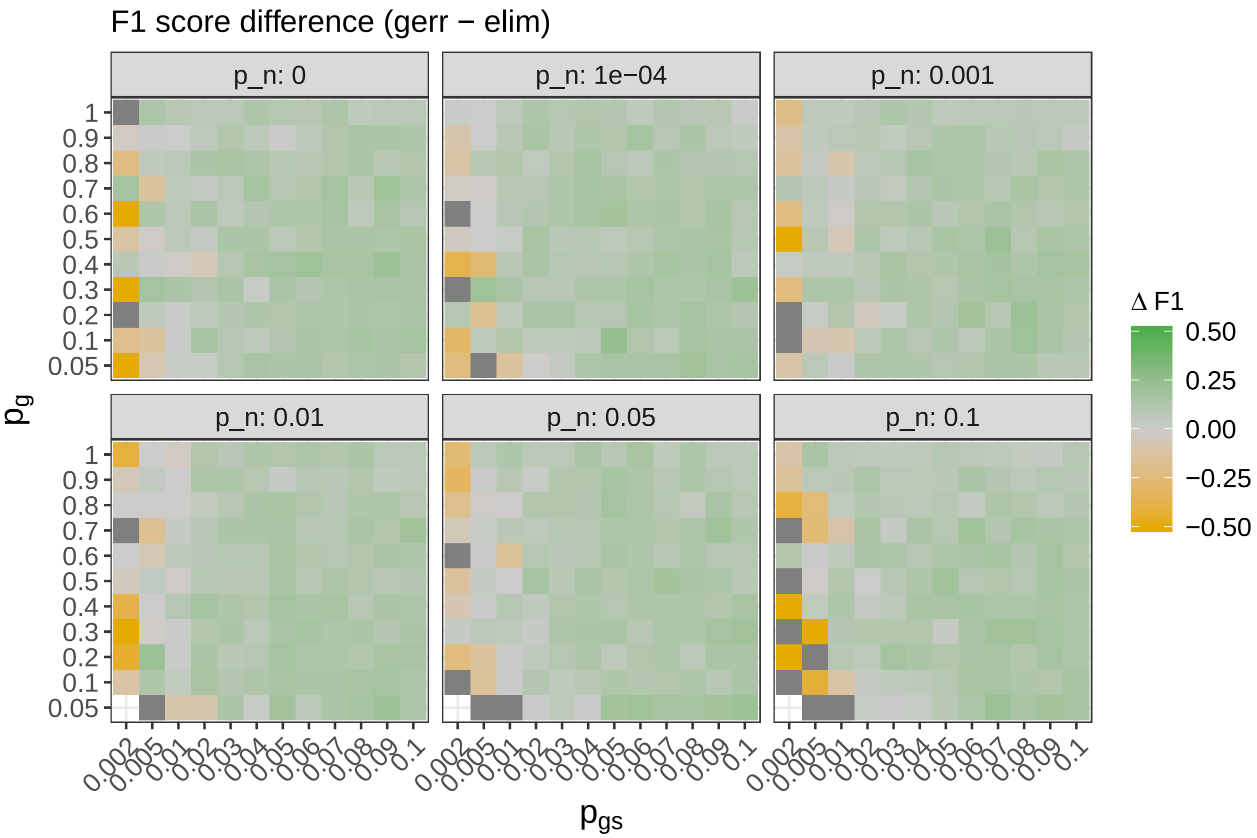
F1 score difference (gerr − elim)
p_n: 0
p_n: 1e−04
p_n: 0.001
p_n: 0.01
p_n: 0.05
p_n: 0.1
$p_g$
1
0.9
0.8
0.7
0.6
0.5
0.4
0.3
0.2
0.1
0.05
0.002
0.005
0.01
0.02
0.03
0.04
0.05
0.06
0.07
0.08
0.09
0.1
$p_{gs}$
Δ F1
0.50
0.25
0.00
−0.25
−0.50

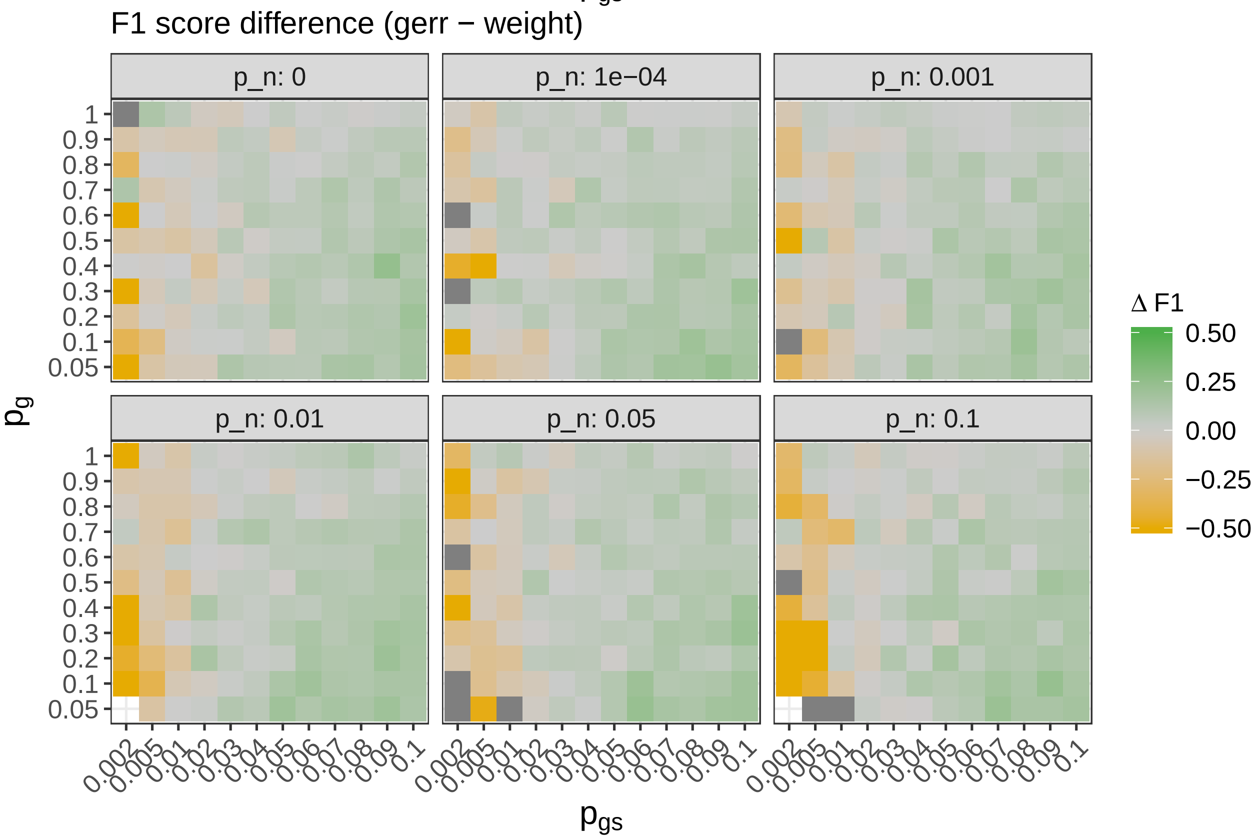
F1 score difference (gerr − weight)
p_n: 0
p_n: 1e−04
p_n: 0.001
p_n: 0.01
p_n: 0.05
p_n: 0.1
$p_g$
1
0.9
0.8
0.7
0.6
0.5
0.4
0.3
0.2
0.1
0.05
0.002
0.005
0.01
0.02
0.03
0.04
0.05
0.06
0.07
0.08
0.09
0.1
$p_{gs}$
Δ F1
0.50
0.25
0.00
−0.25
−0.50

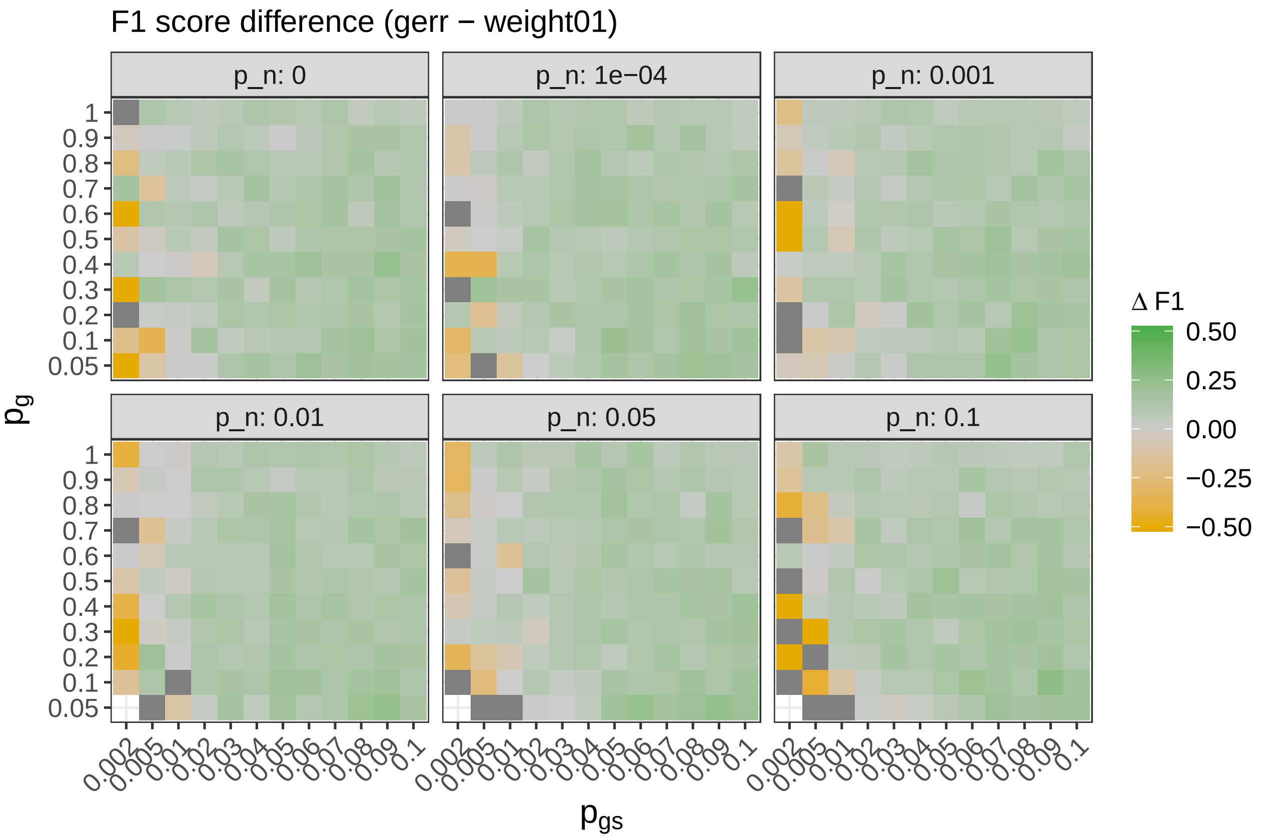
F1 score difference (gerr − weight01)
p_n: 0
p_n: 1e−04
p_n: 0.001
p_n: 0.01
p_n: 0.05
p_n: 0.1
1 0.9 0.8 0.7 0.6 0.5 0.4 0.3 0.2 0.1 0.05
0.002 0.005 0.01 0.02 0.03 0.04 0.05 0.06 0.07 0.08 0.09 0.1
$p_g$
$p_{gs}$
Δ F1
0.50 0.25 0.00 −0.25 −0.50

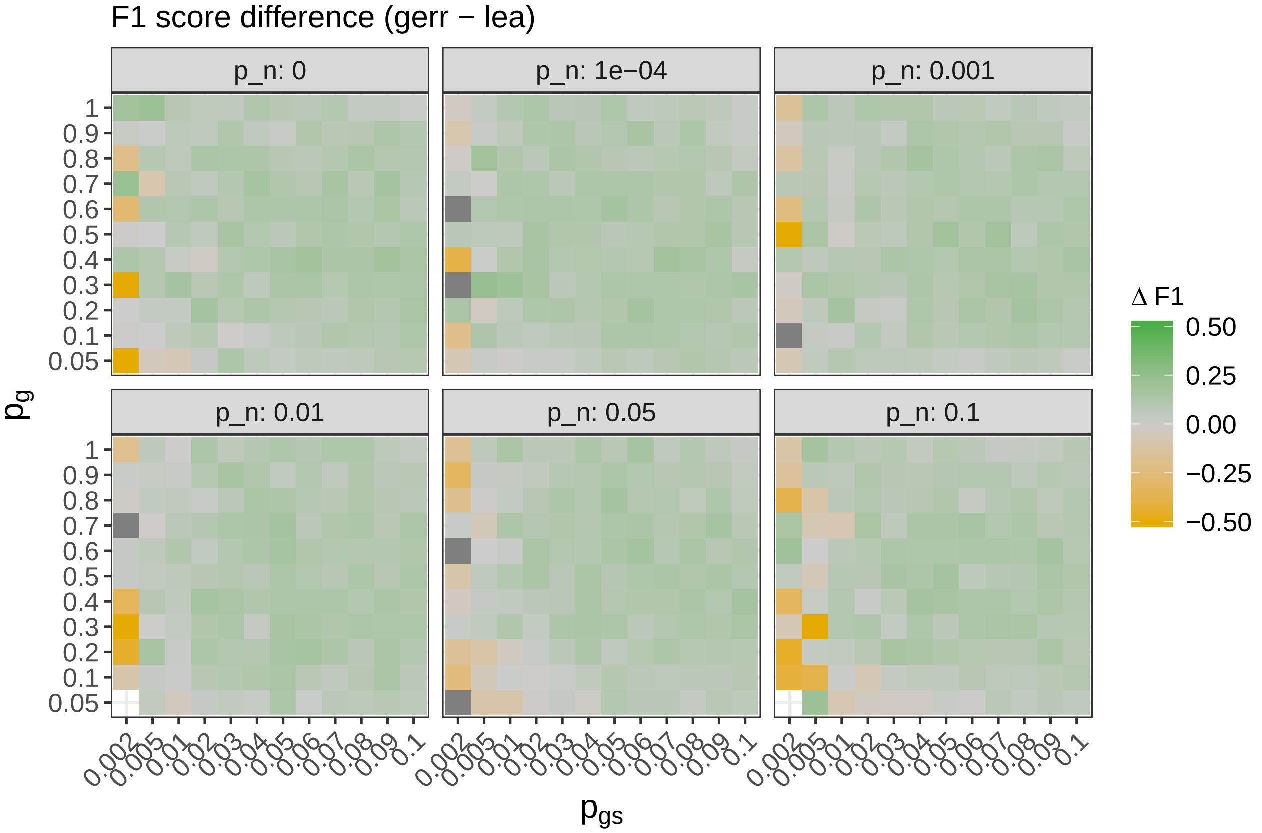
F1 score difference (gerr − lea)
p_n: 0
p_n: 1e−04
p_n: 0.001
p_n: 0.01
p_n: 0.05
p_n: 0.1
1 0.9 0.8 0.7 0.6 0.5 0.4 0.3 0.2 0.1 0.05
0.002 0.005 0.01 0.02 0.03 0.04 0.05 0.06 0.07 0.08 0.09 0.1
$p_g$
$p_{gs}$
Δ F1
0.50 0.25 0.00 −0.25 −0.50

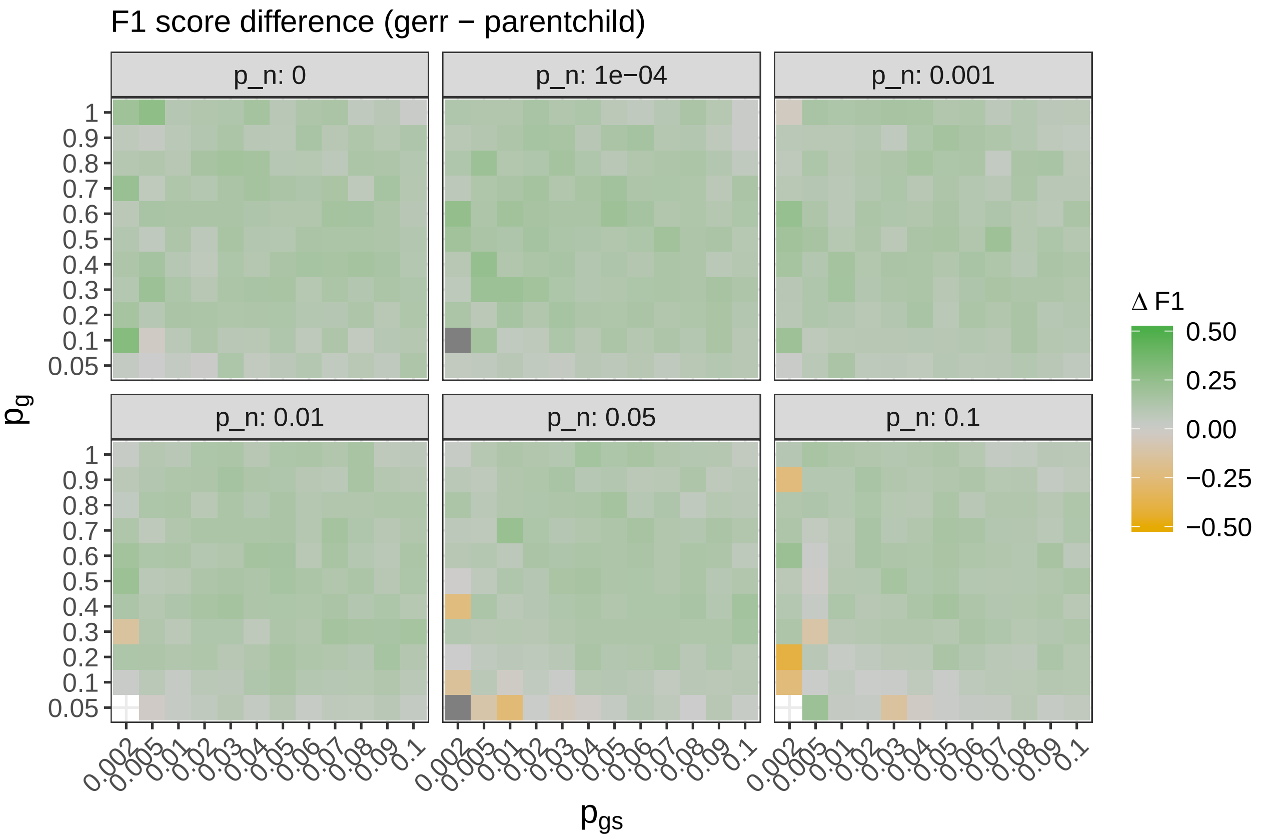

We observe that $F_1$ scores of `gerr` are higher than in all other methods when $p_{gs}$ is low. In other scenarios performance of `gerr` is higher than in other method.s

### Conclusions

We demonstrate that even when working on hierarchical gene sets, `gerr` demonstrates reasonable performance. It demonstrates a high true positive rate with one of the lowest false positive rates. We would like to emphasize that unlike other methods, `gerr` does not take the graph relationships into account.

### R session info

```r
sessionInfo()
```

```
## R version 3.5.1 (2018-07-02)
## Platform: x86_64-pc-linux-gnu (64-bit)
## Running under: CentOS Linux 7 (Core)
##
## Matrix products: default
## BLAS/LAPACK: /pstore/apps/OpenBLAS/0.2.13-GCC-4.8.4-LAPACK-3.5.0/lib/libopenblas_prescottp-r0.2.13.s
##
## locale:
##  [1] LC_CTYPE=en_US.UTF-8       LC_NUMERIC=C
##  [3] LC_TIME=en_US.UTF-8        LC_COLLATE=en_US.UTF-8
##  [5] LC_MONETARY=en_US.UTF-8    LC_MESSAGES=en_US.UTF-8
```

```
##  [7] LC_PAPER=en_US.UTF-8       LC_NAME=C
##  [9] LC_ADDRESS=C               LC_TELEPHONE=C
## [11] LC_MEASUREMENT=en_US.UTF-8 LC_IDENTIFICATION=C
##
## attached base packages:
## [1] stats4    parallel  stats     graphics  grDevices utils     datasets
## [8] methods   base
##
## other attached packages:
##  [1] org.Hs.eg.db_3.7.0   forcats_0.4.0        stringr_1.4.0
##  [4] dplyr_0.8.0.1        purrr_0.3.2          readr_1.3.1
##  [7] tidyr_0.8.3          tibble_2.1.1         ggplot2_3.2.0
## [10] tidyverse_1.2.1      topGO_2.32.0         SparseM_1.77
## [13] GO.db_3.7.0          AnnotationDbi_1.44.0 IRanges_2.16.0
## [16] S4Vectors_0.20.1     Biobase_2.42.0       graph_1.58.2
## [19] BiocGenerics_0.28.0  gridExtra_2.3        MASS_7.3-51.4
## [22] glmnet_2.0-18        foreach_1.5.1        Matrix_1.2-17
## [25] gerr_0.99.22         clustermq_0.8.8
##
## loaded via a namespace (and not attached):
##  [1] httr_1.4.0           bit64_0.9-8          jsonlite_1.6
##  [4] modelr_0.1.4         assertthat_0.2.1     blob_1.1.1
##  [7] cellranger_1.1.0     yaml_2.2.0           pillar_1.3.1
## [10] RSQLite_2.1.1        backports_1.1.4      lattice_0.20-38
## [13] glue_1.3.1           digest_0.6.18        RColorBrewer_1.1-2
## [16] rvest_0.3.2          colorspace_1.4-1     plyr_1.8.4
## [19] htmltools_0.3.6      pkgconfig_2.0.2      broom_0.5.1
## [22] haven_2.1.0          scales_1.0.0         generics_0.0.2
## [25] withr_2.1.2          lazyeval_0.2.2       cli_1.1.0
## [28] magrittr_1.5         crayon_1.3.4         readxl_1.1.0
## [31] memoise_1.1.0        evaluate_0.13        fansi_0.4.0
## [34] nlme_3.1-137         xml2_1.2.0           BiocInstaller_1.32.1
## [37] tools_3.5.1          hms_0.4.2            matrixStats_0.54.0
## [40] munsell_0.5.0        compiler_3.5.1       rlang_0.3.2
## [43] grid_3.5.1           iterators_1.0.11     rstudioapi_0.10
## [46] igraph_1.2.2         labeling_0.3         rmarkdown_1.10
## [49] gtable_0.3.0         codetools_0.2-16     DBI_1.0.0
## [52] reshape2_1.4.3       R6_2.4.0             lubridate_1.7.4
## [55] knitr_1.23           utf8_1.1.4           bit_1.1-14
## [58] rprojroot_1.3-2      stringi_1.4.3        Rcpp_1.0.1
## [61] tidyselect_0.2.5     xfun_0.7
```
