## Supplementary document 4 for "Gene-set Enrichment with Regularized Regression"

*Fang et al.*

### Supplementary Document 4

In this document, we provide two visual elements highlighting the performance of *gerr* when applied to the disease module detection DREAM challenge.

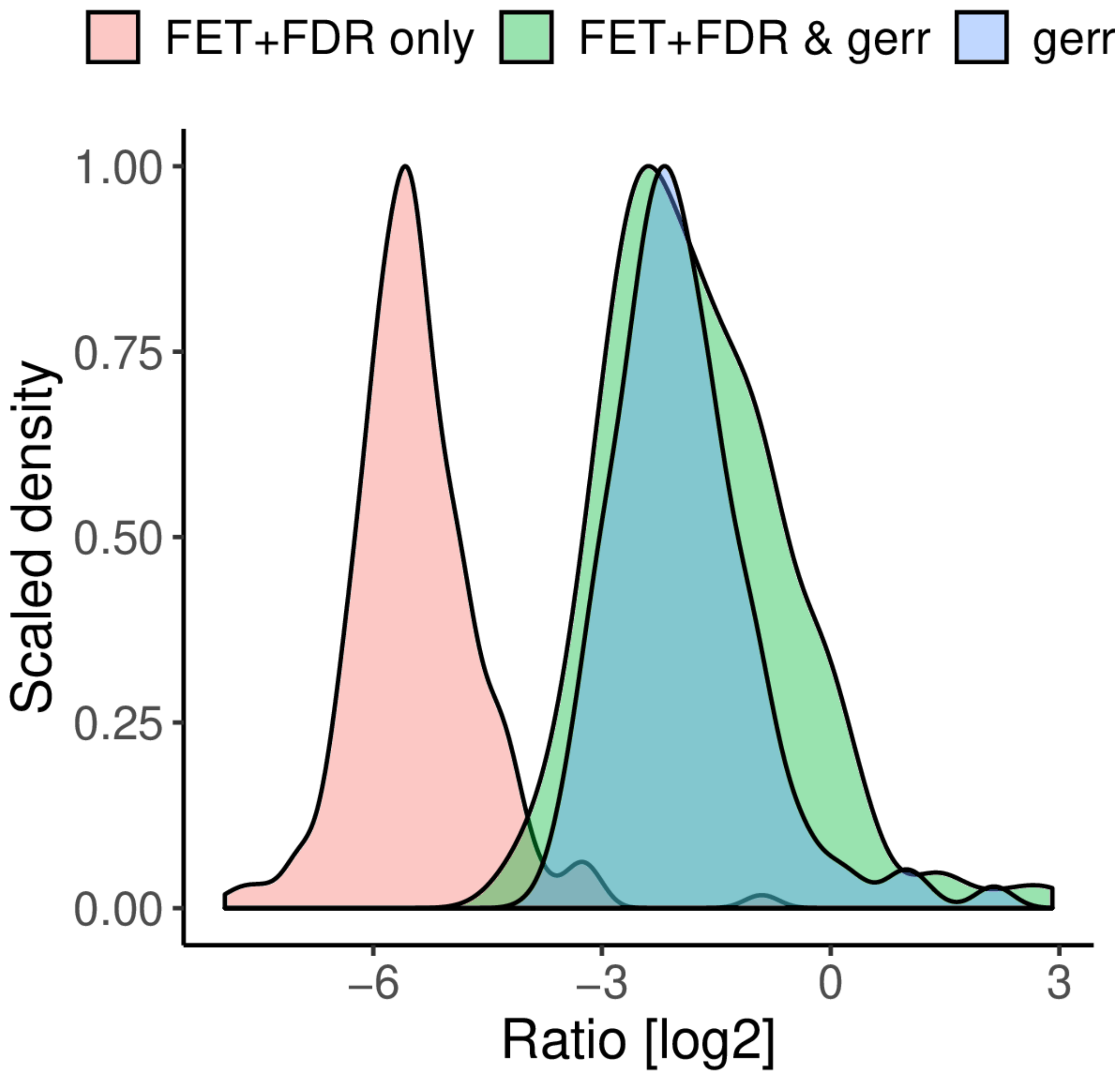


**Figure 1: Hits of *gerr* contain higher proportions of GoI genes compared with hits of FET+FDR**. Ratio (expressed in the logarithm of base 2) of GoI genes over the the number of non-GoI genes in gene-sets identified only by the FET+FDR procedure (*FET+FDR only*), compared with gene-sets identified only by *gerr* (*gerr only*) and gene-sets identified by both

methods (FET+FDR & gerr), across all modules. The x-axis indicates the logarithm ratio of base 2, and the y-axis represents scaled density.

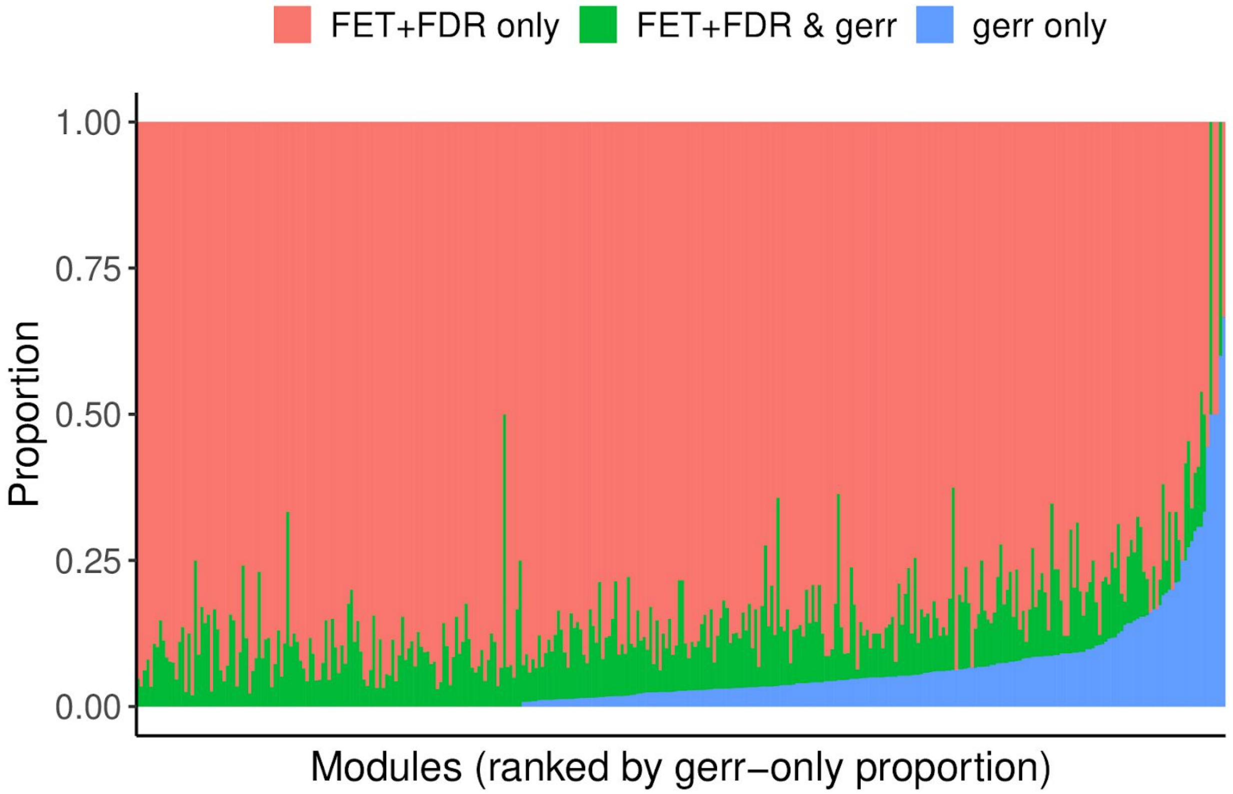


**Figure 2: *gerr* returns gene-sets that are not identified by the FET+FDR procedure in some modules.** Stacked bar plot of the number of unique gene-sets identified only by the FET+FDR procedure (*FET+FDR only*), only by the *gerr* method (*gerr only*), and by both methods (*FET+FDR & gerr*). The x-axis represents DREAM modules that are ordered by the number of unique gene sets that are identified only by the *gerr* method. The y-axis represents the proportion of each category, normalized to 1 by the number of unique gene-sets identified by either method.
